# Distinct extracellular matrix states uncouple collagen accumulation from pathological fibrosis in Duchenne muscular dystrophy

**DOI:** 10.64898/2026.08.11.739868

**Authors:** Pranav Kannan, Daniel Helzer, Ekaterina I. Mokhonova, George R. Marcotte, Tess S. Fleser, Mohammad H. Afsharinia, Joseph C. Reynolds, Jackson Walker, Wenbin Guo, Christina Y. Deng, Philip Farahat, Maxwell C. McCabe, Hannah Tamura, Dongping Qi, Thomas M. Vondriska, Kristen M. Stearns, Rachel Thompson, S. Armando Villalta, Kirk C. Hansen, Amy C. Rowat, Edoardo Malfatti, Valentina Taglietti, Eric J. Deeds, Rachelle H. Crosbie

## Abstract

Fibrosis severity is routinely inferred from collagen abundance, although whether collagen quantity determines pathological fibrosis remains unclear. In Duchenne muscular dystrophy (DMD), chronic muscle injury and inflammation drive extracellular matrix accumulation, making these processes difficult to disentangle. We exploit sarcospan overexpression in *mdx* mice, a model of DMD (*mdx*^TG^), which improves membrane integrity and muscle function despite persistent matrix remodeling. *mdx*^TG^ muscle accumulates more collagen than *mdx* yet lacks its dense macrophage-rich scars. Matrisome proteomics and spatial transcriptomics reveal compositionally and spatially distinct matrix states, while decellularized *mdx*^TG^ matrix protects myotubes from membrane damage relative to *mdx* matrix. Despite these differences, both dystrophic matrices remain stiff and induce nuclear YAP in fibro-adipogenic progenitors. Verteporfin suppresses collagen production and reduces fibrosis *in vivo*, while nuclear YAP is increased in FAPs from patients with DMD. Thus, collagen abundance alone does not define pathological fibrosis; matrix organization, biological activity, and mechanosignaling distinguish functionally distinct fibrotic states.

## Introduction

Fibrosis is commonly quantified by collagen abundance, yet whether extracellular matrix (ECM) quantity predicts its pathological consequences remains unresolved. The composition, crosslinking, architecture, spatial organization, and mechanical properties influence whether deposited ECM disrupts tissue function, supports repair, or alters the behavior of resident cells^1^. These features raise the possibility that pathological fibrosis reflects a qualitative tissue state rather than simply a quantitative increase in collagen. In dystrophic skeletal muscle, repeated cycles of myofiber degeneration and inflammation promote ECM deposition and impair tissue function^2–4^. Macrophage-derived factors, including transforming growth factor-β (TGF-β), activate fibro-adipogenic progenitors (FAPs), which are a principal source of collagen-rich matrix in injured and dystrophic muscle^5–14^. In *mdx* mice, a model of Duchenne muscular dystrophy (DMD), chronic inflammation is evident by four weeks of age and is characterized by sustained macrophage infiltration at sites of muscle injury^15,16^. Because tissue injury, inflammation, and collagen accumulation occur concurrently, it remains unclear whether all collagen-rich matrices are biologically equivalent or which matrix properties distinguish pathological scarring from adaptive tissue remodeling.

The ECM not only provides structural support, but also functions as an instructive component of the cellular niche. Changes in matrix composition, ligand presentation, crosslinking, topology, and mechanics regulate cytoskeletal tension and mechanosensitive transcription in stromal cells. ECM stiffness and cytoskeletal tension promote nuclear localization of the transcriptional co-activators YAP and TAZ and induction of matrix-producing gene programs^17^. In skeletal muscle, YAP/TAZ-associated activity is enriched in fibrotic regions and can be activated downstream of TGF-β signaling, linking biochemical and mechanical inputs in the regulation of FAP cell fate^18^. Fibroblasts can also retain mechanically induced activation after removal from a stiff environment, demonstrating that prior matrix exposure can maintain pro-fibrotic behavior^19^. These observations raise the possibility that a matrix may lack the dense organization of destructive scarring while remaining capable of sustaining stromal signaling and continued ECM production. However, these ECM-producing pathways have largely been studied in the context of ongoing pathological injury, making it unclear whether mechanical signaling actively sustains fibrosis or instead amplifies upstream inflammatory cues.

We previously demonstrated that sarcospan (SSPN), a component of the dystrophin-glycoprotein complex^20–26^, improves membrane stability, muscle function, and dystrophic pathology in *mdx* mice across skeletal, respiratory, and cardiac muscle^27–32^. Unexpectedly, ECM remodeling and collagen accumulation persist despite this broad physiological improvement^30^. Moreover, decellularized *mdx*^TG^ matrix supports muscle progenitor function more effectively than laminin-scarred *mdx* matrix, indicating that collagen abundance alone is insufficient to define pathological fibrosis and that matrix composition and organization contribute to functional outcomes^31^. The *mdx* and *mdx*^TG^ models therefore provide a unique opportunity to compare collagen-rich but biologically distinct ECM states and to separate matrix abundance from pathological fibrosis; we use the term ECM state to refer to the combined composition, spatial organization, and signaling properties of the matrix.

Here, we take an integrative approach spanning molecular to tissue scales: combining bulk, single-cell, single-nucleus, and spatial transcriptomics with quantitative matrisome proteomics, tissue mechanics, and cell-matrix functional assays to interrogate the cellular behaviors and matrix properties that distinguish these ECM states. We find that collagen in *mdx* muscle is concentrated within macrophage-rich necrotic lesions, whereas *mdx*^TG^ muscle contains greater total collagen that is more evenly distributed despite improved tissue integrity and reduced focal inflammatory pathology. FAPs remain the principal source of fibrillar collagen in both dystrophic models and adopt ECM-producing transcriptional programs. Although ECM states differ in spatial organization and inflammatory association, both *mdx* and *mdx*^TG^ matrices remain mechanically remodeled and induce nuclear YAP localization in wild-type FAPs. Pharmacological YAP blockade reduces collagen production by dystrophic FAPs and attenuates fibrosis *in vivo*. These findings demonstrate that collagen abundance alone does not define pathological fibrosis and identify ECM spatial organization and matrix-driven mechanosignaling as distinct features of the fibrotic state.

## Methods and materials

### Animal models

Wild-type (C57BL/6J and DBA/2J strains) and *mdx* mice (C57BL/10ScSn and DBA/2J strains) were obtained from Jackson Laboratories (Bar Harbor, ME, USA). The generation of *mdx*^TG^ mice expressing full-length SSPN cDNA has been previously described^27,33^. The SSPN transgenes were driven by the human skeletal *α*-actin promoter, resulting in approximately 3-fold overexpression of human SSPN (line 3, hSSPN). Male mice 7-20 weeks old were used for all experiments, except the verteporfin treatment study, which included male and female mice with treatment beginning at two weeks of age. For drug treatment experiments, pharmacological YAP1 inhibition was induced by three weekly intraperitoneal injections of verteporfin (Sigma Aldrich, SML0534) or DMSO (ATCC, 4X) vehicle control at a dosage of 50 mg/kg for a total of five weeks. All mice were housed and maintained in the vivarium according to approved guidelines by the UCLA Institutional Animal Care and Use Committee (protocol #2000-029).

### DMD human biopsies for proteomics analysis

Gastrocnemius muscle biopsies were obtained from 3 typically developing individuals (14-16 years old) with no history of neuromuscular disorders, and from 4 individuals with DMD (10-14 years old). 3 of the 4 DMD participants were non-ambulatory at the time of tissue collection. All participants were enrolled at a tertiary care academic referral center. All studies utilizing human samples were approved by the UCLA Institutional Review Board (IRB 21-000350). All methods were carried out in accordance with relevant guidelines and regulations and performed in accordance with the Declaration of Helsinki. Written informed consent was obtained from parents or caregivers, and age-appropriate assent was obtained from all participants.

### DMD human biopsies for immunofluorescence

Three DMD deltoid muscle biopsies were collected from 4-6 years old ambulant steroid free boys. Three histological normal age-matched biopsies have been used as control. The use of these samples for research purposes was approved by the ethical Committee issued by our institutions for myopathology in compliance with the Helsinki Declaration. Patients’ parents gave informed consent for muscle biopsy studies, according to French (Comité de Protection des Personnes Est IV DC 2012–1693).

### Bulk RNA sequencing

RNA sequencing of this dataset was first collected and published as described in previous studies^30^ (GEO accession: GSE262341). Briefly, tibialis anterior muscles were collected from 12-week-old wild-type (n=5), *mdx* (n=4), and *mdx*^TG^ (n=4) mice, immediately frozen in liquid nitrogen. The frozen tissue was powdered using a pre-chilled mortar and pestle, followed by homogenization in TRIzol reagent (Thermo Fisher Scientific, 15596026). RNA extraction was performed *via* TRIzol phase separation, with subsequent purification using the RNeasy column system (QIAGEN, 74104), following the manufacturer’s protocols. RNA concentration and quality were assessed using the TapeStation 4200 (Agilent Technologies). Libraries were constructed using the TruSeq Stranded mRNA Library Prep Kit (Illumina, San Diego, CA, USA), and sequenced with 75bp pair-end reads (targeting 55 million read pairs per sample) on Illumina HiSeq 4000 platform at the UCLA Technology Center for Genomics & Bioinformatics (TCGB). The raw sequencing data of the 13 samples underwent quality assessments with FastQC (version 0.11.9). Sequencing adapters and low-quality bases were removed using fastp (version 0.23.2). The processed reads were then aligned to the mouse reference genome (GRCm38, ensemble v93) using STAR (version 2.7.9a), and gene count matrices were generated with featureCounts from the Subread package (version 2.0.3). Genes with at least 10 counts in at least 2 samples are included for the downstream analysis. Gene expression counts normalized by sequencing depth (CPM, counts per million) were calculated using edgeR (version 3.38.4) and Log transformation with a pseudocount 1 was applied for principal component analysis (PCA) and heatmap. Top 2000 most variable genes were used for PCA plots. Differential gene expression analyses were carried out using edgeR. To control for false discovery rate (FDR), the Benjamini–Hochberg (BH) procedure was applied to adjust *P*-values, and genes with an adjusted *P*-value below the 0.05 threshold and absolute log2FoldChange greater than 1 were identified as significantly differentially expressed. Based on the expression levels in different sample groups, the differential expressed genes were classified into three categories. Gene Ontology over-representation analysis was performed on the gene subsets using clusterProfiler (version 4.4.4), and the dotplot is used to visualize the biological process pathways.

### Cytokine analysis

Serum from 7-week-old wild-type, *mdx*, and *mdx*^TG^ was collected and submitted to the UCLA Immune Assessment Core for cytokine and TGF-β1 ELISA analysis. For the collection, blood was collected directly into Microtainer Serum Separator Tubes (BD, 365967) by retro-orbital route. The tubes were then inverted 5 times to mix and allowed to incubate at room temperature for 30 mins to clot before being centrifuged at room temperature at 7000*g* for 7 mins. The serum supernatant was collected and stored at -80°C.

### Single cell isolation from skeletal muscle

A single cell suspension was prepared from 15-week-old male wild-type, *mdx*, and *mdx*^TG^ mice as described previously^34–36^ with modifications. Mice were euthanized with isoflurane followed by cervical dislocation. All hindlimb muscles were collected and stored in ice cold wash buffer (WB) (Dulbecco’s Modified Eagle Medium/Nutrient Mixture F-12 with 15 mM HEPES (StemCell Technologies, 36254) supplemented with 1% penicillin/streptomycin (Thermo Fisher, 15140122), and 10% heat-inactivated fetal bovine serum (FBS) (Thermo Fisher, 16140071). Muscles were trimmed of visible fat and tendon, then transferred into 1 mL of muscle digestion buffer (MDB) (wash buffer containing 700 U/mL collagenase II (Worthington; LS004176)) and minced into a slurry. The muscle slurry was then transferred to a 50 mL conical tube containing 10 mL of MDB and incubated horizontally in a 37°C water bath for 1 hr with agitation. MDB was diluted with 40 mL of ice-cold WB, centrifuged 500*g* for 5 min at 4°C. The supernatant was aspirated down to 8 mL, and 1 mL of a stock collagenase II solution (1000 U/mL in DPBS (Corning, 21-030-CM)) and 1 mL of stock dispase II solution (11 U/mL dispase (Thermo Fisher, 17105-041)) were added. The pellet was triturated with a 5 mL serological pipette to further breakdown the tissue. The suspension was incubated horizontally in a 37°C water bath for 30 min with agitation. The suspension was passed through a 10 mL syringe with a 20G 1-inch needle 10 times, diluted with 40 mL ice cold WB, and centrifuged 500*g* for 5 min at 4°C. The supernatant was removed down to 10 mL and the pellet was resuspended with a 10 mL pipette and filtered through a 40 μm cell strainer (Fisher Scientific, 22363547). The conical tube was washed twice with 10 mL of WB and filtered, then the filtered solution was diluted to 50 mL with WB and centrifuged 600*g* for 15 min at 4°C. The pellet was resuspended in 0.5 mL DPBS and transferred to a 15 mL conical tube for debris removal using Debris Removal Solution (Miltenyi Biotec, 130-109-398) following the manufacturer’s instructions. The pellet was resuspended in 5 mL of ammonium-chloride-potassium (ACK) lysing buffer (Thermo Fisher, A1049201), transferred to a 50 mL conical tube, and incubated for 5 min at room temperature. The suspension was then diluted to 50 mL with 10% FBS in DPBS and centrifuged 300*g* for 10 min at room temperature. Dead cells were removed using the Dead Cell Removal Kit (Miltenyi Biotec, 130-090-101) and the QuadroMACS separator (Miltenyi Biotec, 130-090-976) following the manufacturer’s instructions. The live cells were collected, centrifuged 300g for 10 min at 4°C, and resuspended in 0.5 mL of 0.04% bovine serum albumin (BSA) (UltraPure BSA, Thermo Fisher, AM2618) in Dulbecco’s Phosphate-Buffered Saline (DPBS). The suspension was centrifuged 300*g* for 5 min at 4°C, resuspended in 0.3 mL of 0.04% BSA, and the cells were counted. The suspension was then centrifuged 300g for 5 min at 4°C and resuspended to 1200 cells/μL.

### Single nucleus isolation from frozen skeletal muscle samples

Nuclei isolation was performed in accordance with the recommended 10x Genomics protocol and as described previously^37,38^ with modifications. Quadriceps muscles from wildtype, *mdx*, and *mdx*^TG^ mice were dissected, snap frozen in liquid nitrogen, and stored under liquid nitrogen until use. On the day of the experiment, frozen quadriceps muscles were placed on ice and transferred to a 2 mL microcentrifuge tube. 1 mL of NP40 lysis buffer (10 mM Tris HCl pH 7.4 (Sigma T2194), 10 mM NaCl (Sigma 59222C), 3 mM MgCl_2_ (Sigma M1028), 0.1% IGEPAL CA-630 (Sigma I8896)) and minced for 2-3 min on ice. Minced samples were then mixed and dounce homogenized using 10 strokes of a loose pestle and 20 strokes of a tight pestle (VWR 62400-595) followed by centrifugation at 500*g* for 5 min at 4°C. Supernatant was discarded and samples were resuspended in 1 mL of BSA solution (1% BSA (Miltenyi Biotec 130-091-376), 0.2 U/μL RNase inhibitor (Sigma 3335402001)) and centrifuged 100*g* for 1 min at 4°C to pellet large debris. The supernatant was filtered through 70 (Corning 352350) and 30 μm (Fisher scientific NC9682496) filters and centrifuged 500*g* for 5 min at 4°C. The supernatant was discarded and the resultant pellet was resuspended in 5 mL of BSA solution. 5 μL of 7AAD (final concentration 10 μg/mL) (Sigma SML1633-1ML) was added and the nuclei suspension was mixed and incubated on ice for 5 min protected from light. 1 mL of BSA solution was added and samples were centrifuged 500*g* for 5 min at 4°C. The supernatant was discarded and the nuclei pellet was resuspended in 0.3 mL of BSA solution. Nuclei were isolated by positive staining for 7AAD using FACS. Purified nuclei were centrifuged 500*g* for 5 min at 4°C and concentrated to 1200 cells/μL.

### Single cell library construction, sequencing, and analysis

The single cell/nucleus suspensions (described above) were submitted to the Technology Center for Genomics and Bioinformatics core at UCLA using the 10X Genomics platform. Single cell library was prepared using Chromium Single cell 3’ Library Construction. Single cell samples were then sequenced using Illumina NovaSeq S1 (2x50, 1 lane). Single nuclei samples were sequenced using Illumina NovaSeq SP (2x50, 2 lanes). The data was first processed using the 10X Genomics CellRanger pipeline. The resultant filtered count matrices were analyzed in Python (v3.9.7) using Scanpy^39^ (v1.9.3). For both datasets, we applied an initial filtering step to retain only cells expressing at least 200 genes and genes that were expressed in at least three cells. Additional quality control was performed by excluding cells with more than 6000 detected genes and cells where mitochondrial genes accounted for more than 10% of the total gene expression (only required for cell data). This was followed by data normalization to a total count of 10,000 molecules per cell and logarithmic transformation. For single cell sequencing, we next regressed out the effects of total counts per cell and the percentage of mitochondrial gene expression and performed data scaling to a maximum value of 10. The optimal number of principal components was determined, and a neighborhood graph was constructed using 40 principal components and 10 nearest neighbors. Clustering of cells was then performed on this graph using the Leiden algorithm^40^. Manual cluster annotation was performed using literature-established markers^41^. Differentially expressed genes (DEGs) were determined using the Wilcoxon rank-sum test. Gene set scoring was conducted using UCell^42–44^. YAP/TAZ target gene set was curated from published gene lists^44–46^. Pearson correlation coefficients were compared between genotypes by first converting to *r* value to *z*-scores using Fisher *r*-to-*z* transformation 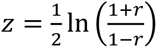 followed by standard error calculation as 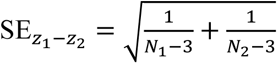 and a two-sided *z*-test using 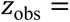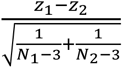 with Holm correction. Bootstrap standard errors were calculated to verify the accuracy of analytic standard errors.

### Flow cytometry

Single-cell isolation was performed as described previously^10^. Briefly, mice were subjected to whole body perfusion with PBS followed by hind-limb muscle dissection. Hind-limbs were minced and digested with 0.2 mg/mL collagenase P (Roche) and 20 μg/mL DNAse (Roche) in serum-free Dulbecco’s Modified Eagle Medium (DMEM) for 2, 30-minute rounds. Cells were filtered through 70 and 40 μm cell strainers, suspended over 5 mL of Lymphoprep (Stem Cell Technologies), and centrifuged at ∼315*g* for 30 minutes at room temperature. Cells in the interphase were collected and stained with with 100 μL of Zombie NIR viability dye (1:1000 in 1× PBS) for 15 minutes on ice (protected from light) for live/dead cell discrimination. An anti-CD16/32 antibody (clone 2.4G2) was used for Fc-receptor blocking before staining. Single-cell suspensions were stained with a previously established antibody panel^10^ followed by analysis on a BD FACSAria Fusion flow cytometer with FACSDiva software (BD Bioscience). Post-acquisition analysis was performed with Flowjo software version 10.8.

### Spatial RNA preparation and sequencing

Quadriceps muscles from 15-week-old wild-type, *mdx*, and *mdx*^TG^ mice were dissected, washed twice in DPBS, and submerged in a 2 mL microtube containing 4% PFA (Fisher Scientific, 50-980-494) for 24 hrs at 4°C in the dark. The tissues were washed 3× in DBPS and submerged in 70% ethanol. The fixed tissues were then delivered to the Translational Pathology Core Laboratory at UCLA for paraffin embedding and sectioning. The Technology Center for Genomics and Bioinformatics core at UCLA used the 10X Chromium Visium platform (V2 Visium kit) for library construction. The Visium spatial library was sequenced using Illumina Novaseq SP (2x50, 1 lane). The sequenced data was then processed using the 10X SpaceRanger pipeline.

### Spatial RNA sequencing analysis

The resulting filtered count matrices from SpaceRanger were used for data analysis with Seurat v5.0.1^47^ (R v4.3.1). Counts were normalized using the default parameters set in the NormalizeData function in Seurat. Pathway enrichment was performed using the DEenrichRPlot function in Seurat with the following parameters: DEGs identified using the Wilcoxon rank-sum test, a log fold change threshold of 0.25, a Benjamini-Hochberg corrected *P*-value cutoff of 0.05, the maximum number of genes input set to 1000, and the Enrichr^48^ query database set to “GO_Cellular_Component_2023”. Gene set scoring was conducted using UCell^42,43^. Patch size and number of patches were determined by first calculating local Moran’s I using the R package Voyager^49^ with statistical significance determined by a permutation test with 100,000 permutations for each genotype and gene set score followed by Benjamini-Hochberg correction. Spots with significant (FDR < 0.05) local Moran’s I were connected through the neighbor graph determined by Voyager, which uses the hexagonal lattice structure of the Visium platform. Directly adjacent significant spots, or those linked through a chain of adjoining significant spots, were grouped into a single patch. Patch size was calculated as the number of spots comprising each patch. Global Moran’s I (univariate spatial autocorrelation of each gene set) and global Lee’s L (bivariate spatial autocorrelation between gene sets) were calculated using the Voyager package. Global Moran’s I and Lee’s L were compared between genotypes using the observed statistics and jackknife leave-one-out standard errors. For each gene set, one gene was removed and Moran’s I was recalculated, iterating through all genes (*i* = 1 to *n*). Standard errors were calculated as 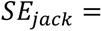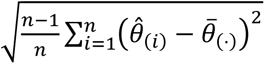, where θ̄_(⋅)_ is the mean Moran’s I from all leave-one-out estimates 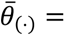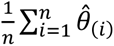 and *θ̂_i_* is the Moran’s I calculated with gene *i* removed from the gene set. For Lee’s L, this same approach was conducted by removing *i* in gene set 1 and calculating Lee’s L between the leave-one-out gene set 1 and the full gene set 2, iterating over all genes in gene set 1. This was repeated by calculating Lee’s L between the full gene set 1 and the leave-one-out gene set 2, iterating over all genes. *P*-values were determined using a two-sided *Z*-test with 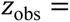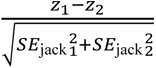, where *z*_1_ is the observed statistic in group 1 and *z*_2_ is the observed statistic in group 2. Pairwise z-test *P*-values were adjusted using Holm correction.

### Preparation of myoscaffolds

Muscle decellularizations were conducted as described previously^31^. In brief, transverse 50-micron cryosections from quadriceps muscle were placed on adhesive MAS slides (Newcomer Supply, 5077W90) and were dried at room temperature for 30 mins. Slides were then placed in a Coplin jar containing 1% sodium dodecyl sulfate (SDS) (Fischer Scientific, MP04811030) for 30 mins under constant rotation. Slides were then rinsed in 1X PBS for 30 mins, followed by two additional 30 mins rinses in diH_2_O followed by 1X PBS. Myoscaffolds were stored in PBS at 4°C until use. Pavone nanoindentation measurements to determine mechanical properties of myoscaffolds were performed within 48 hrs of decellularization. For the creatine kinase release assay, sections were attached to the bottoms of 12-well tissue culture plates. Sectioned muscles were exposed to 1% SDS for 10 minutes on a rotator at 50 rpm. SDS was removed, and slides were washed with PBS for 15 minutes. PBS wash was repeated two times, followed by a 30-minute wash in DI water. Water was removed, and a final 40-minute PBS wash was performed.

### Tissue stiffness measurements

Endomysium stiffness was measured using force probe indentation (Pavone, Optics11 Life, The Netherlands) with a probe tip stiffness of 0.40 N/m and tip radius of 26.5 μm. The instrument was calibrated with the probe tip submerged in PBS using a geometrical factor <5% variation than in air. Indentations (maximum indentation force: 0.1 μN, indentation speed: 10 μm/s) were performed across an array of selected endomysium regions on decellularized tissue samples. The Hertzian model fitting for indentation profiling (up to 16% probe tip radius according to the instrument manual, R^2^ > 0.90) was used to determine an effective Young’s modulus.

### Bone marrow-derived macrophage isolation

L929 fibroblasts were cultured in T175 flasks to 100% confluency in RPMI-C (Standard RPMI 1640 with 10% FBS, 1% penicillin/streptomycin, 1 mM Na^+^ Pyruvate, 2 mM L-glutamine, and 10 µM HEPES, 50 µM 2-mercaptoethanol) and cultured for an additional 7 days to obtain L929 conditioned media containing macrophage colony-stimulating factor (M-CSF). The media was collected, filtered through 0.22 µm, and aliquoted prior to storage at -80°C. Mouse femurs and tibia were sterilized with 70% ethanol, washed twice with RPMI-C, then crushed at a 90° angle in RPMI-C. The suspension was filtered through a 70 µm filter and collected. Femurs and tibias were crushed and filtered twice more in RPMI-C to maximize bone marrow cell yield. The final suspension was centrifuged at 1000 rpm (∼200*g*) and frozen in 1 mL aliquots of 12×10^6^ cells/mL in heat-inactivated fetal bovine serum (FBS) with 10% DMSO at -80°C. After three days, bone marrow cells were transferred to liquid nitrogen for long-term storage. Bone marrow cells were thawed in a 37°C water bath and plated into 10-cm non-treated petri dishes (∼3×10^6^ cells/dish). The cells were differentiated to macrophages for 7 days using macrophage growth media (RPMI-C supplemented with 25% L929 conditioned media), adding an additional 5 mL of media on day 5. On day 7, bone marrow-derived macrophages (BMDMs) were ready for experimentation.

### BMDM culture on myoscaffolds

BMDMs were removed from 10-cm plates using 5 mM EDTA in PBS and seeded onto slides at a density of 1.0×10^5^ BMDMs per well (2 wells per slide). For phagocytosis, BMDMs were cultured overnight on myoscaffolds followed by incubation for 1 hour (37°C, 5% CO_2_) with Zymozan A conjugated bioparticles (ThermoFisher Z23373) at a multiplicity of infection of 10^50^. Phagocytosis was stopped by ice cold PBS washes followed by fixation with 4% PFA for 10 min at room temperature. For YAP1 staining, BMDMs were stimulated with 100 ng/mL LPS or IL-4 to serve as positive controls for pro-inflammatory and anti-inflammatory phenotypes, respectively. Positive control groups and unstimulated negative control groups were seeded onto uncoated slides while experimental groups were seeded onto slides pre-coated with 50 µm decellularized WT, *mdx*, or *mdx*^TG^ myoscaffolds. BMDMs were incubated for 24 hours prior to fixation with 4% PFA. BMDMs were permeabilized with 0.3% Triton X-100 in PBS for 10 min then blocked with 3% BSA for 1 hr. Cells were incubated for 1.5 hours at room temperature with rabbit anti-YAP1 (Abcam AB205270, 1:250) primary antibody. Cells were then washed in PBS and incubated with anti-rabbit AF488 (Abcam, ab150077, 1:500) for YAP1 detection and phalloidin-AF594 conjugate (ThermoFisher A12381, 1:100) for 1 hour. All fluorescence images were then captured using an Axioplan 2 fluorescent microscope equipped with 20× differential interference contrast objectives and the Axiovision Rel 3.0 software. Length/width ratio measurements and cell counts were conducted in ImageJ.

### Creatine kinase release assay

Healthy H2K immortalized mouse myoblasts and H2K *mdx* myoblasts with a nonsense mutation in exon 23 of dystrophin^51^ were a gift from Terrance Partridge, Ph.D. (Children’s National Medical Center, Washington, D.C.). Cells were cultured in DMEM (ThermoFisher Scientific, 11965118) supplemented with 20% FBS (Invitrogen, 10082-147) and maintained at 33°C with 5% CO_2_ on Matrigel (BD Biosciences) coated plates. Cells were trypsinized (Fisher Scientific, MT25051CI) and seeded at 50,000 cells per well in growth media onto 12-well plates coated in 50 μm thick myoscaffolds. Cells were allowed to proliferate until they reached 80-90% confluency before switching to differentiation media. After five days of differentiation, a hypoosmotic salt solution (100 mosmol) was created by adding sucrose to a salt solution of 5 mM HEPES, 5 mM KCl, 1 mM MgCl_2_, 5 mM NaCl, 1.2 mM CaCl2, 1 mM glucose. Differentiation media was removed, and cells were exposed to 300 μL of the osmotic solution for 20 minutes at 37°C. After incubation, the supernatant was isolated from each well and kept on ice. Cells were then removed from the myoscaffolds using trypsin, centrifuged, and resuspended in DI water. Cells were pipetted to break apart cell clusters and lysed by rapid freezing in liquid nitrogen and warming in a water bath. This freeze/thaw cycle was repeated three times. This final lysate was stored on ice. The percentage of creatine kinase in the supernatant versus lysate was determined using the Creatine Kinase-SL kit (Sekisui Diagnostics, 326-10) according to the manufacturer’s instructions.

### FAP isolation and culture

Primary FAPs were isolated using the single cell isolation method described in “Single cell isolating from skeletal muscle” section above, with the following additions. After live cell collection, PDGFRα^+^ cells were positively selected using anti-CD140a (PDGFRα) microbeads (Miltenyi Biotec, 130-101-502) and MACS separation^52^ following the manufacturer’s instructions. The PDGFRα^+^ cells (FAPs) were centrifuged at 300*g* for 5 min and plated in 12-well plates at a density of 10,000 cells/cm^2^ in FAP expansion media (high glucose DMEM (Fisher Scientific, 11-965-118) supplemented with 10% FBS, 1% penicillin/streptomycin, 2.5 ng/mL bFGF (Thermo Fisher; 450-33))^41,53–55^.

### Cell-matrix functional assays

Primary FAPs isolated from wild-type mice were seeded on 50 μm myoscaffolds prepared by decellularizing quadriceps muscle sections derived from wild-type, *mdx*, and *mdx*^TG^ mice and allowed to adhere for 24 hrs in the presence of DMSO or 5 µM verteporfin. Following this period, cells were rinsed with 1X PBS and fixed with 4% paraformaldehyde for 10 mins. Following fixation, cells were permeabilized with 0.3% Triton-X for 10 mins, followed by two 5 min PBS washes. The cells were then blocked with 3% BSA for 1 hour and incubated with rabbit-anti YAP1 antibody (Abcam, AB205270, 1:250) overnight at 4°C. Following overnight incubation, cells were washed in PBS 3 X and incubated in anti-rabbit Alexa-488 secondary antibody (Abcam, ab150077, 1:500) diluted in 3% BSA for 1 hour at room temperature. Cells were washed three times with PBS and cover-slipped with Prolong Gold Antifade Mountant (Fischer Scientific, P36982).

### Mechanically tunable gels and cell seeding

Polydimethylsiloxane (PDMS) gels were prepared according to previously established protocols^56,57^. The gels were prepared using published ratios of components part A and part B of Dow Corning Sylgard 527 silicone dielectric gel (Ellsworth Adhesives, 1696742) and were subsequently used to coat 6-well plates by overnight incubation at 60°C. The stiffness of the hydrogels was confirmed by force probe indentation (Pavone, Optics-11 Life). To minimize probe tip-sample adhesion effects, the tip was immersed in 1% Pluronic solution for 2 hours and then washed twice with deionized water prior to measurements. Force probe indentation was performed under the following conditions: probe stiffness, 0.54 N/m; tip radius, 3 μm; and indentation speed, 10 μm/s. The gels were then sterilized by ultraviolet irradiation and coated with 1 μg/ml bovine fibronectin in PBS (Sigma, F1141) for 4 hrs at 37°C followed by PBS washes. For cell seeding experiments, primary FAPs obtained from *mdx* mice (n=2) were seeded at a density of 10,000 cells/cm^2^ and treated with DMSO or 5 µM verteporfin cultured in the presence or absence of 1 ng/ml TGF-β1 (Thermo Fisher, 100-21) for 14 days.

### Hydroxyproline assays

Following 14 days of treatment, collagen content was measured by a hydroxyproline assay adapted from Creemers *et al.* 1997 and Cissell *et al.* 2017^58,59^. Briefly, five wells per experimental condition were decellularized with 10 mM NH_4_OH (Sigma, 09859), 0.25% Triton-X (Sigma, T8787) in PBS for three min^60^ followed by three PBS washes. After decellularization, 6N HCl (Fisher Scientific, SA49) was added to each well followed by gentle scraping using cell scrapers to remove deposited ECM. The ECM was then transferred to a 2 ml tube and hydrolyzed in HCl for 3.5 hrs at 135°C. Samples were evaporated completely dry and dissolved in 200 μL of deionized water followed by centrifugation at 9470*g* for 7 mins. Chloramine-T solution (1.6% w/v chloramine T (Thermo Fisher Scientific, A12044.30), 190 mM citric acid (Sigma, C2404-100G), 610 mM sodium acetate (Sigma, S2889), 150 mM acetic acid (Fisher Scientific, A38-212), 650 mM NaOH (Fisher Scientific, S320-1), 30% v/v 2-propanol (Fisher Scientific, A416-1)) was added to each sample and mixed, and incubated for 20 mins at room temperature. Erlich’s solution (25% w/v 4-dimethylaminobenzaldehyde (Sigma, 109762), 70% v/v 2-propanol, 30% v/v 10N HCl) was added to each sample, mixed, and incubated for 30 mins at 60°C. Final solutions were aliquoted in a 96-well plate in duplicate and absorbance was measured at 560 nm. Hydroxyproline content was determined using the standard curve generated with 0-50 μg/mL *trans*-4-hydoxy-L-proline (Fisher Scientific, AAA1185109). Hydroxyproline quantification of mouse muscle followed the same procedure as deposited ECM samples. Diaphragms were dissected, frozen in liquid nitrogen, and pulverized. 5-10 mg of diaphragm samples were carefully weighed and hydrolyzed in HCl as described above.

### Proteomics sample preparation and analysis

Proteomics analysis of wild-type, *mdx*, and *mdx*^TG^ mice was first published and has previously been described in Stearns-Reider *et al.* 2023 and McCourt *et al.* 2023^30,31^. This quantitative proteomics method yielded 79 ECM proteins, represented as absolute quantities in nmol/g of muscle tissue, making standard statistical methods designed for 1000s of proteins and intensity values unfit. Therefore, pairwise comparisons were conducted using Welch’s *t*-test followed by Benjamini-Hochberg correction. Proteins represented as fractions within functional class were compared between genotypes by first transforming values by centered log ratio followed by pairwise permutation analysis of variance (PERMANOVA) tests using the vegan and RVAideMemoire R packages.

### Immunohistochemistry and histological analysis

Indirect immunofluorescence staining was performed using 10 μm quadriceps muscle cryosections. Sectioned tissue was fixed and permeabilized in ice cold acetone or fixed with 4% formaldehyde followed by permeabilization with 0.3% Triton-X solution for 15 min. Tissue sections were rinsed with PBS and blocked with 3% BSA in PBS for 30 mins at room temperature. Sections were then incubated in primary antibody overnight at 4°C with the following antibodies: rabbit anti-collagen type I (Cedarlane Labs, CL50151AP-1, 1:250), rat anti-CD68 (Serotec, MCA1957, 1:100), rat anti-CD206 (Serotec, MCA2235, 1:50), goat anti-PDGFRα (Thermo Fisher, AF1062SP; 1:200), goat anti-human PDGFRα (biotechne, AF-307-NA, 1:100), rabbit anti-YAP1 (AB205270; 1:100; Abcam). The sections were then washed with PBS and detected by the following secondary antibodies: anti-rabbit Alexa Fluor 488 (Thermo Fisher, A-21121; 1:200), anti-rabbit Alexa Fluor 594 (Thermo Fisher, A-21207, 1:200), anti-goat Alexa Fluor 488 (Thermo Fisher, A-11055; 1:200), anti-rat Alexa Fluor 647 (Thermo Fisher, A-21247, 1:200). All sections were mounted with Vectashield Antifade Mounting Medium (Vector Laboratories, H-1200-10) or Prolong Gold Antifade Mountant. Hematoxylin & Eosin protocols have been previously described^29^. Briefly, sections were acclimated at room temperature for 30 mins and stained with hematoxylin for 5 mins. The samples were then rinsed with ddH_2_O, stained with eosin for 5 mins, and dehydrated in a series of ethanol rinses of 70, 80, 90 and 100% before final incubation in xylene. For Picrosirius red staining, tissues were acclimated to room temperature for 30 mins. Samples were then rehydrated by incubating in 100%, 80% and 50% ethanol solutions following by soaking in ddH_2_O. The tissue sections were then stained with hematoxylin for 5 mins, washed three times with ddH_2_O, and stained with Picrosirius red for one hour. The tissues were subsequently dehydrated in a series of ethanol rinses of 70, 80, 90 and 100% before final incubation in xylene. All images were captured using an Axioplan 2 fluorescence microscope with Axiovision 3.0 software (Carl Zeiss Inc., Thornwood, NY, USA) or LSM 880 (Carl Zeiss Microscopy, Germany) on Zeiss Axio Observer Z1 inverted microscope stand running Zen Black 2.3. All quantifications were performed using ZenBlue (Carl Zeiss), imageJ (NIH) and QuPath^61^. Quantification methods are further described in respective figure legends.

### Pearson dispersion for macrophage spatial distribution analysis

Wild-type, *mdx*, and *mdx*^TG^ quadriceps muscle sections were probed for CD68 as described above. Dispersion was calculated using multiple individual images per sample. We assumed positive cell counts follow a Poisson process where variance is equal to the mean. Under complete spatial randomness (CSR; in this context, random arrangement of positive cells throughout a tissue section), we can expect dispersion (variance of positive cell counts divided by mean positive cell counts across images in a given sample) to be approximately equal to 1, and dispersion > 1 indicates clustering. Formally, Pearson dispersion is defined as 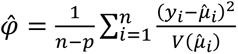 is the variance function implied by the assumed distribution and *p* is the number of fitted parameters. For each genotype *g*, we fit a Poisson generalized linear model with a separate intercept for each sample (animal), *s* = 1, …, *k_g_*, where *k_g_*is the number of animals in genotype *g*. Under this saturated model, *V*(*μ*) = *μ* and the maximum-likelihood fitted value for every image in sample *s* is exactly that sample’s own mean positive cell count, 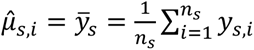 for all *i* ∈ *s* where *y_s_*_,*i*_ is the observed positive cell count in the *i*-th image in sample *s*, and *n_s_* is the number of images taken in sample *s*. Substituting these two properties into the general Pearson dispersion formula yields a closed-form expression for genotype-level dispersion

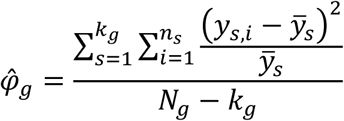

where *N_g_* = ∑*_s_n_s_* is the total number of images pooled across all samples in genotype *g*. This closed-form expression is identical to the Pearson dispersion statistic obtained by the generalized linear model (glm) function in R but avoids the computational cost of iteratively reweighted least squares. Pearson dispersion distributions were calculated by bootstrap resampling (10,000 resamples) at the image level, resampling each animal’s own images independently with replacement and preserving sample labels within genotype.

### Index of dispersion from whole quadriceps images

Whole tissue images were imported into QuPath for positive cell detection. Detection coordinates (total cells and CD68^+^ cells) and regions of interest (free of image artifacts) were exported and analyzed in R and tissue windows were built using the spatstat R package. Positive cell density was visualized using Kernel density estimation (KDE) with a shared bandwidth between all samples. A shared bandwidth was estimated using the method defined by Berman and Diggle^62,63^ and implemented in the spatstat R package^64^. Only tissue samples with >20 positive cells were used for bandwidth estimation to avoid estimation instability due to low cell density, and the median bandwidth across all samples was used for KDE and visualization. Dispersion analysis was conducted by first overlaying grids with varying bin sizes (50, 60, 70, 80, 90, and 100 μm) onto each tissue section and counting positive cells within each bin. Index of dispersion *D* was calculated as 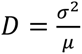, where *σ*^2^ is the variance of positive cell counts between bins and *μ* is mean positive cell counts across all bins. *D* ≈ 1 indicates CSR, while *D* > 1 suggests clustering. *D* was calculated for all bin sizes, and average *D* was used as a summary statistic per sample. Bins falling outside the tissue region of interest, including internally excluded regions, were excluded from analysis. Bootstrap standard errors were calculated for each sample using 10,000 resamples. For each bin size and tissue sample, sampling with replacement was conducted on the observed bin counts, drawing a number of values equal to the number of bins at that bin size. *D* was calculated in each bootstrap resample identically to the observed test static (averaging over all bin sizes), and bootstrap standard errors per sample were calculated as the standard deviation of all 10,000 *D* values per sample. These bootstrap standard errors and observed *D* values were used to compare genotypes by fitting a random-effects meta-regression with the Hartung-Knapp small-sample adjustment followed by Holm multiple comparisons correction.

### Identification of spatially clustered macrophages in quadriceps cross-sections

Spatial cluster analysis was conducted using cell coordinates. To identify CD68^+^ cells with statistically significant spatial clustering, we first implemented Ripley’s *K* function. Briefly, Ripley’s *K* function is defined as

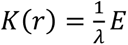 [number of additional points within *r* of a randomly chosen point],

where *λ* is the number of points per unit area^65^. Here, the estimator is defined (in the R package spatstat) as 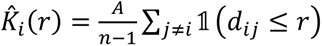 where *A* is the tissue area, *n* is the total number of positive cells, *d_ij_* is the distance between positive cells *i* and *j*, and 1(⋅) is the indicator function. Under a homogenous Poisson process (i.e., for spatial data, CSR), *E*[*K*(*r*) ∣ CSR] = *πr*^2^^65^. The most common use of Ripley’s *K* to test CSR is through the transformation 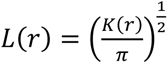 since 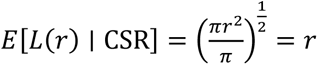; that is, under CSR, the number of positive cells within radius *r* of a given randomly chosen positive cell is *r*^65^. Based on this, for each positive cell *i*, we calculated the excess number of cells above expected as excess*_i_*(*r*) = *L_i_*(*r*) − *r*. A cell was only tested at radius *r* if its distance to the nearest excluded region was at least *r*. For each positive cell, we tested radii ℛ = {50, 60, 70, 80, 90, 100} μm and determined the maximum test statistic 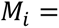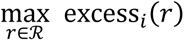. Next, for each tissue section, we simulated 999 random spatial arrangements of positive cells *i*^∗^ using the rpoint function in spatstat and calculated excess*_i_*∗ (*r*) for each simulated positive cell. Excess values were collapsed to a single maximum across all simulated positive cells per simulation *s* as 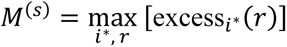. We compared the observed *M_i_* values to these simulation maxima and observed CD68^+^ cells were considered significant if they fell above the 99^th^ percentile threshold *T* from all simulations *T* = *Q*_0.99_({*M*^(1)^, . . ., *M*^(999)^}). Lastly, cells identified as significant were clustered using density-based clustering with applications with noise (DBSCAN). DBSCAN was applied to the coordinates of significant CD68^+^ cells, with minimum points equal to 1 and epsilon values of 50, 60, 70, 80, 90, and 100 μm tested. For each epsilon value, we recorded patch size and number of patches and took the average across all tested values.

### Statistical analysis

Data are presented as mean ± standard deviation or boxplots. Analysis was performed in R (version 4.3.1), Python (version 3.11.9) and GraphPad Prism 11. Permutation tests with step-down maximum T *P*-value adjustment or Benjamini-Hochberg correction were applied to control family-wise error rate or false discovery rate, respectively. When sample sizes were sufficiently large, linear regression with multiple comparisons correction was applied, and results were validated with permutation approaches. For clustered data, we fit generalized linear models with cluster-robust standard errors followed by Tukey’s multiple comparison correction. Statistical analyses of RNA sequencing data are described in their corresponding sections and in figure legends as appropriate. A *P*-value < 0.05 was considered statistically significant.

### Data availability

The data generated and reported in this paper are available from the corresponding author upon request. Bulk RNA sequencing (GEO accession: GSE262341), scRNA-seq, snRNA-seq, and spatial data (GEO accession: GSE343269) that support the findings of this study have been deposited in GEO and are available at the time of publication. DMD FAP datasets were obtained from Single cell Broadinstitute.org repository, accession number SCP2678 (reference^66^). All analytic code is available at https://github.com/dphelzer/mechanobiology_2026 (commit ID a967389).

## Results

### Extracellular matrix remodeling persists despite partial normalization of inflammatory transcriptional programs

To determine whether fibrotic remodeling remains coupled to inflammatory pathology, we leveraged the *mdx*^TG^ model in which membrane integrity and muscle function are improved despite persistent ECM remodeling^28,29,32^, providing an opportunity to distinguish persistent matrix remodeling from overall dystrophic disease severity. We first performed bulk RNA sequencing on skeletal muscle samples from 12-week-old wild-type, *mdx*, and *mdx*^TG^ mice. Principal component analysis (PCA) revealed distinct clustering of the three genotypes (**Fig. 1A**). Differential gene expression (DEG) analysis showed the greatest number of DEGs between *mdx*^TG^ and wild-type mice (2484), followed by *mdx* and wild-type mice (1627) (**Fig. 1B**). Although *mdx*^TG^ mice exhibit broad physiological improvement relative to *mdx* (**Supplementary Table 1**), their transcriptional profiles remain distinct from wild-type muscle^28,29,32,67,68^, suggesting that SSPN activates dystrophin-independent gene-regulatory networks that contribute to functional rescue^30^. Differentially expressed genes were therefore classified into two categories: unrestored genes for which *mdx*^TG^ expression resembled *mdx* but differed from wild-type and restored genes for which *mdx*^TG^ expression resembled wild-type but differed from *mdx* (**Fig. 1C**). ECM remodeling pathways were significantly enriched among unrestored genes, indicating that matrix-associated transcriptional programs remain elevated in *mdx*^TG^ muscle relative to wild-type (**Fig. 1D-E**). Conversely, immune-related Gene Ontology terms, including inflammation, cytokine production, and leukocyte migration/chemotaxis, were predominantly enriched among restored genes (**Fig. 1F-G**). Together, these data demonstrate that SSPN shifts bulk inflammatory transcriptional signatures towards wild-type, while ECM remodeling programs remain transcriptionally distinct from wild-type. Because bulk sequencing reflects shifts in overall tissue composition, we next used single-cell and spatial approaches to determine what this transcriptional shift reflects at cellular resolution.

**Figure 1.**
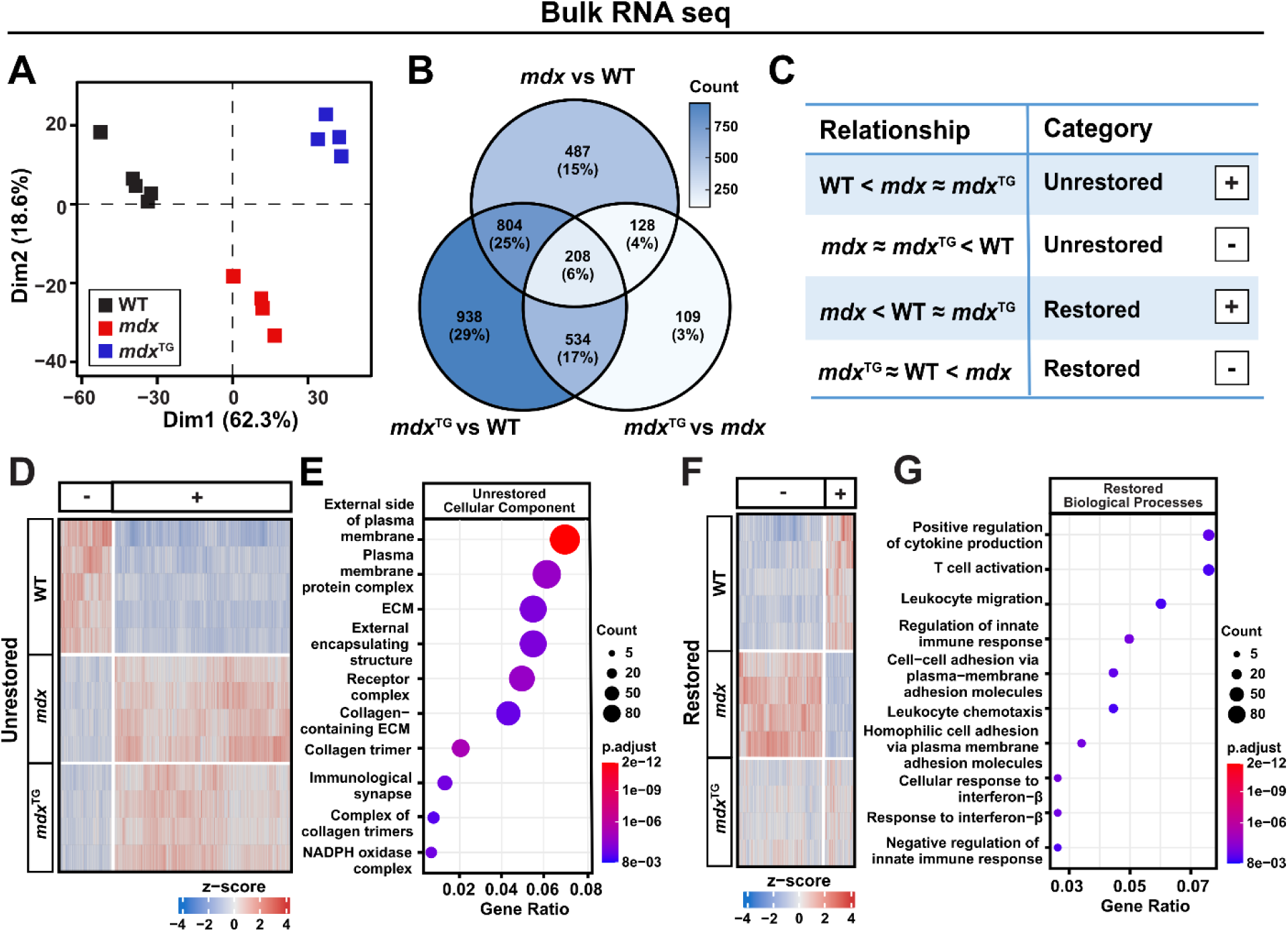
Extracellular matrix remodeling persists despite partial normalization of inflammatory transcriptional programs in *mdx*^TG^ muscle. (**A**) Principal component analysis (PCA) of bulk RNA sequencing data of 12-week, wild-type (n=5), *mdx* (n=4), and *mdx*^TG^ (n=4) tibialis anterior muscles. (**B**) Venn diagram illustrating differential gene expression (DEG) analysis comparing wild-type, *mdx*, *mdx*^TG^ genotypes. (**C**) Table summarizing gene relationships categorized as unrestored (*mdx*^TG^ significant difference from wild-type, no significant difference from *mdx*), restored (*mdx*^TG^ no significant difference from wild-type, significant difference from *mdx*), “+” and “-” indicate whether gene expression was upregulated or downregulated within each category, respectively. (**D**) Heatmap displaying the expression of unrestored genes. (**E**) Gene ontology (GO) terms enriched in unrestored component in *mdx*^TG^ mice, ranked by gene count and adjusted *P*-value. (**F**) Heatmap displaying the expression of restored genes. (**G**) GO terms enriched in restored component in *mdx*^TG^ mice, ranked by gene count and adjusted *P*-value.

### Interstitial cells account for the major transcriptional changes in *mdx*^TG^ muscle

To identify the cellular populations responsible for the transcriptional differences observed by bulk RNA-sequencing, we performed single-cell (scRNA-seq) and single-nucleus (snRNA-seq) RNA sequencing of wild-type, *mdx*, and *mdx*^TG^ skeletal muscle. scRNA-seq primarily captured interstitial and immune cells, while snRNA-seq predominantly captured mature myonuclei (**Fig. S1A**), with scRNA-seq capturing greater total RNA and gene detection than snRNA-seq (**Fig. S2A-C**)^69^.

Because myonuclei represented the largest proportion of nuclei in snRNA-seq, we examined this population first. DEG analysis revealed substantial transcriptional differences across all genotype comparisons, with the greatest number of DEGs in *mdx* versus wild-type (**Fig. S1B**), and myonuclei were highly heterogeneous between genotypes (scPerturb energy distance)^70^ (**Fig. S1C-D**). Given enrichment of ECM and membrane-adhesion pathways among unrestored genes in bulk RNA-seq, we examined cell-ECM adhesion gene expression and found that *Sgcg*, *Sgcd*, *Utrn*, *Itgav*, and *Itgb1* were upregulated in *mdx*^TG^ myonuclei relative to *mdx*, with *Utrn* and *Itgb1* also elevated above wild-type (**Fig. S1E-F**). This is consistent with SSPN’s known role in increasing abundance of the utrophin-glycoprotein and α7β1-integrin complexes at the sarcolemma, where they compensate for dystrophin loss (**Table 1**)^27,71^. GO analysis confirmed enrichment of focal adhesion and cell-substrate junction terms in *mdx*^TG^ relative to both *mdx* and wild-type (**Fig. S1G**). Despite these changes, the strongest DEGs identified in myonuclei and bulk RNA-seq showed little overlap (Jaccard distances 0.89–0.95 across comparisons), indicating that most bulk transcriptional differences arise from non-myonuclear, interstitial cell populations rather than the myonuclei compartment itself, motivating a closer examination of the immune landscape.

### Inflammatory remodeling is attenuated but not eliminated in *mdx*^TG^ muscle

To determine how SSPN alters the inflammatory environment associated with persistent ECM remodeling, we next analyzed the immune landscape in *mdx* and *mdx*^TG^ muscle using the scRNA-seq dataset. Differential gene-expression analysis identified >1,000 DEGs detected in nearly all genotype comparisons (**Fig. 2A**). Macrophages from both *mdx* and *mdx*^TG^ muscle retained expression of ECM-associated genes, including fibrillar collagens (*Col1a1, Col1a2, Col3a1*), TGF-β1 (*Tgfb1*), matrix remodeling enzymes (*Lgmn, Ctss, Ctsl, Mmp19*), and platelet-derived growth factor gene expression (*Pdgfa, Pdgfc*), consistent with continued stromal remodeling activity^11^ (**Fig. 2B**). Recent studies identified a distinct macrophage subpopulation associated with skeletal muscle fibrosis that is characterized by high *Lgals3* expression and elevated osteopontin (*Spp1*)^10^. Bulk RNA sequencing, spatial transcriptomics, and scRNA-seq revealed a significant reduction in expression of genes associated with this *mdx*-enriched, pro-fibrotic macrophage program in *mdx*^TG^ muscle, including *Spp1*, a key mediator of macrophage-FAP interactions^10^, while *Lgals3* expression remained elevated (**Fig. 2B, Fig. S3A-B, Fig. S4**). In parallel, expression of osteopontin receptors was significantly reduced in *mdx*^TG^ FAPs relative to *mdx* FAPs (**Fig. S3C**), supporting attenuation of osteopontin-dependent macrophage–FAP signaling in *mdx*^TG^ muscle. Consistent with these observations, the pro-fibrotic macrophage gene-set score^10^ remained elevated in *mdx* compared to wild-type and slightly reduced in *mdx*^TG^ macrophages compared with *mdx* (**Fig. 2B-C**). Macrophages from both dystrophin-deficient genotypes (*mdx* and *mdx*^TG^) remained transcriptionally more similar to pro-fibrotic macrophages than to classically polarized M1 (pro-inflammatory) or M2 (anti-inflammatory) macrophages^72^ (**Fig. 2C**), indicating that SSPN attenuates selected inflammatory macrophage programs without completely normalizing macrophage state.

**Figure 2.**
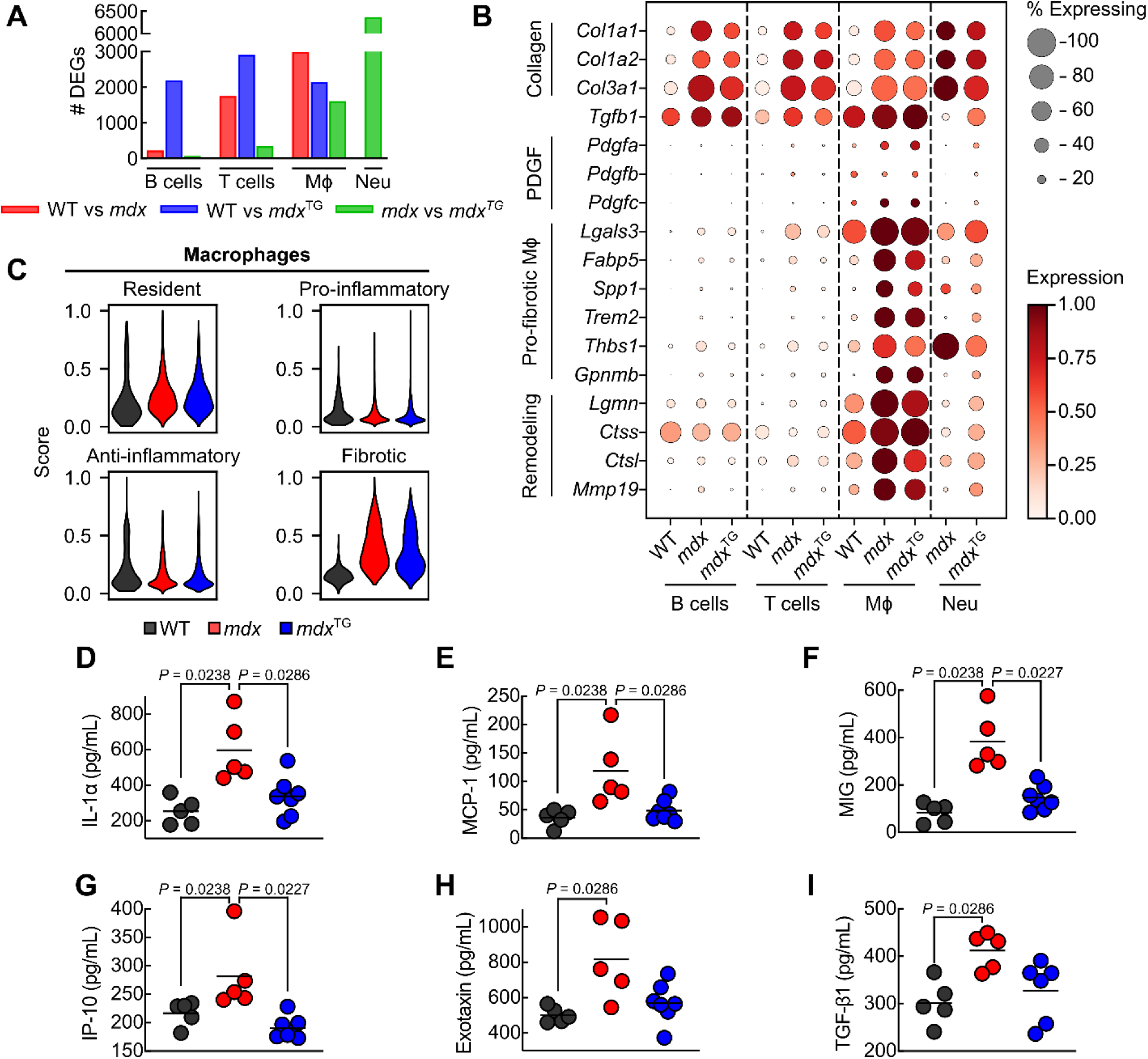
The inflammatory response is reduced in *mdx*^TG^ muscle. (**A**) Number of differentially expressed genes (DEGs) between wild-type, *mdx*, and *mdx*^TG^ immune cells annotated from scRNA-seq. Mϕ, macrophages; Neu, neutrophils. Few differentially expressed genes for B cells in wild-type versus *mdx* and *mdx* versus *mdx*^TG^ is likely related to the low number of B cells (cell count: 84, 0.74% of all cells) in *mdx* (**B**) Dot plot of collagens, platelet-derived growth factor (PDGF), pro-fibrotic Mϕ genes, and ECM remodeling genes. (**C**) Violin plot of macrophage gene set scores. Serum levels of inflammatory cytokines and chemokines, including (**D**) IL-1α, (**E**) MCP-1, (**F**) MIG, (**G**) IP-10, (**H**) Eotaxin, and (**I**) TGF-β1 (n=5-7 biological replicates). Statistical analysis by permutation test.

To determine whether these cellular changes were reflected systemically, we measured circulating cytokines in wild-type, *mdx*, and *mdx*^TG^ mice. Among the analytes measured, the pro-inflammatory cytokine IL-1α, a key regulator of leukocyte trafficking^73^, along with its downstream chemokines MCP-1, MIG, and IP-10, were reduced in *mdx*^TG^ serum relative to *mdx* (**Fig. 2D-I**). TGF-β1, produced predominantly by anti-inflammatory macrophages during muscle repair^74^, was highest in *mdx*. Because these circulating cytokines originate from multiple cell types (**Fig. S4D**), they do not directly define macrophage phenotype but instead reflect the overall inflammatory environment. Together, these findings indicate that inflammatory signaling is substantially attenuated, but not eliminated in *mdx*^TG^ muscle.

### Inflammation is spatially reorganized despite persistent macrophage abundance

Although inflammatory transcriptional programs were partially restored in *mdx*^TG^ muscle, it remained unclear whether this reflected reduced macrophage infiltration or reorganization of the inflammatory landscape. We therefore quantified macrophage spatial organization across entire quadriceps cross-sections. Spatial transcriptomic analysis revealed that the pan-leukocyte marker *Ptprc* and macrophage genes *Cd68* and *Mrc1* were highly concentrated within discrete inflammatory lesions in *mdx* muscle, but present at lower levels and more diffusely distributed in *mdx*^TG^ muscle (**Fig. 3A**). Gene set enrichment analysis (GSEA) of *Ptprc*⁺ and *Cd68*⁺ regions in *mdx* muscle demonstrated significant enrichment of fibrosis, inflammation, and macrophage activation pathways compared to wild-type and *mdx*^TG^ counterparts (**Fig. S5A-F**).

**Figure 3.**
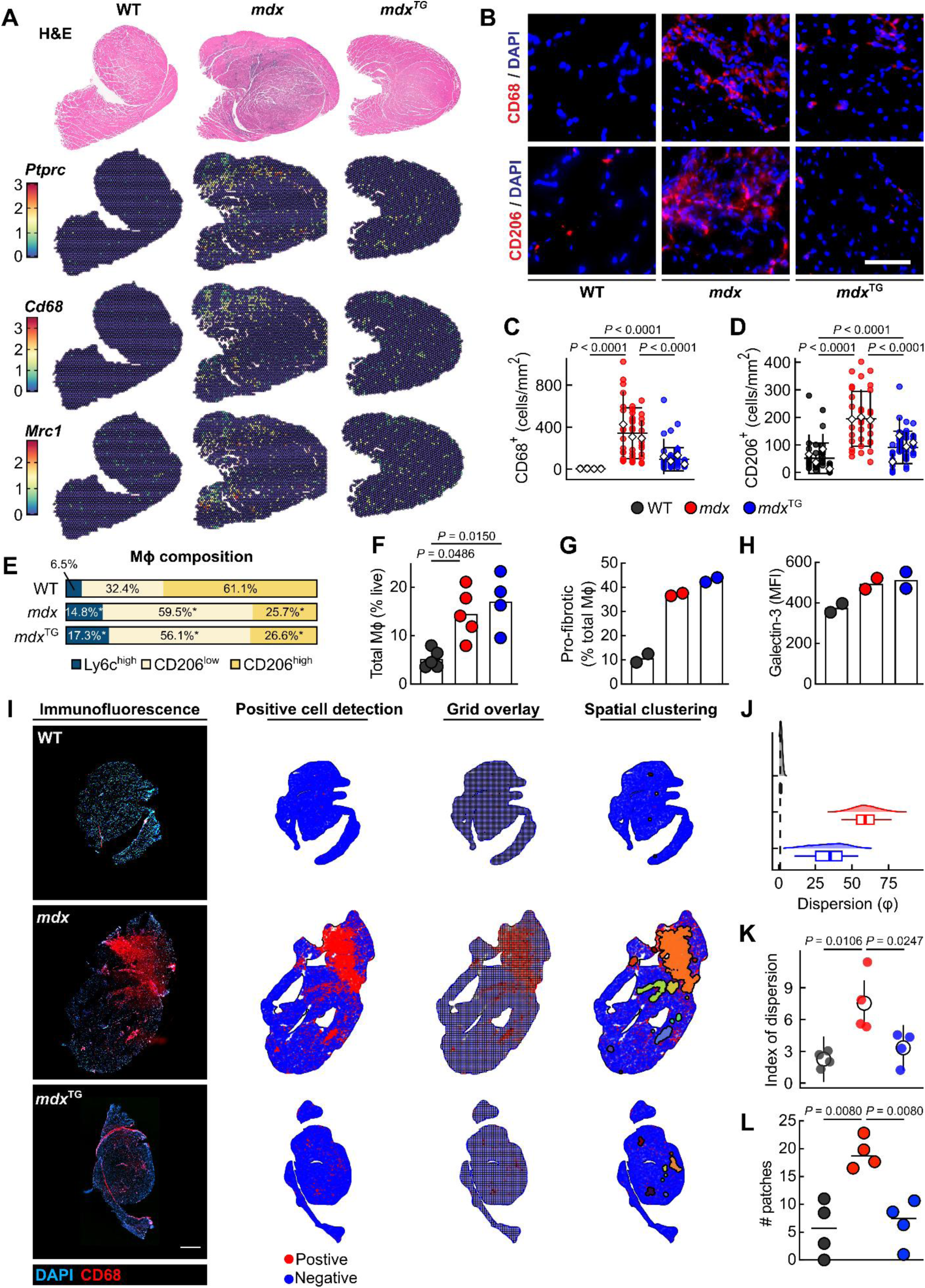
The inflammatory landscape is reorganized in *mdx*^TG^ muscle. (**A**) Representative H&E staining and corresponding spatial RNA sequencing plots showing the expression of leukocyte and macrophage genes (*Ptprc*, *Cd68*, *Mrc1*) in 15-week-old cross-sections of wild-type, *mdx*, *mdx*^TG^ quadriceps muscle. (**B**) Representative immunofluorescence images of macrophage markers CD68 and CD206, and DAPI, with (**C**) quantification of CD68^+^ and (**D**) CD206^+^ macrophages. At least ten micrographs were analyzed per biological replicate in 12-week wild-type (n=4), *mdx* (n=3), and *mdx*^TG^ (n=4) mice. Cell density per image is shown for each biological sample. Mean cell density per sample are represented by a white diamond. Scale bar = 200 μm. (**E**) Composition of macrophage subsets, expressed as percentage of total macrophages. The three macrophage populations shown are a subset of total macrophages and were re-normalized to 100% for visualization. (**F**) Flow cytometry analysis of macrophage abundance (as percentage of total live cells) in wild-type (n=5), *mdx* (n=5), and *mdx*^TG^ (n=4) muscle. (**G**) Pro-fibrotic macrophage abundance and (**H**) Galectin-3 (*Lgals3*) mean fluorescence intensity (MFI). *Indicates statistically significant difference compared to wild-type. Statistical analysis by permutation test with step-down max T *P*-value adjustment. (**I**) Immunofluorescence images of whole quadriceps cross-sections in wild-type, *mdx*, and *mdx*^TG^ probed with anti-CD68 antibodies. Scale bar, 1000 μm. After image acquisition, CD68^+^ cells were identified by positive cell detection using QuPath. Each section was overlaid by a grid of varying sizes (bin size of representative image was set to 75 μm for illustrative purposes only). An example of CD68^+^ cell spatial clusters identified by the transformed Ripley’s K function (defined in methods) followed by DBSCAN is shown. (**J**) Dispersion of macrophage counts estimated from individual images per sample (images from data presented in B-D). Density plots show dispersion parameters from 10,000 bootstrap resamples with boxplots showing median, 1^st^ and 3^rd^ quartile, and 95% confidence intervals. (**K**) Index of dispersion for each whole quadriceps tissue cross-section (n = 4 wild-type, *mdx*, and *mdx*^TG^ quadriceps). Samples are represented as individual points. Statistical analysis by fitting a random-effects meta-regression using estimates and bootstrapped standard errors with the Knapp-Hartung small-sample adjustment followed by Holm multiple comparisons correction. Mean and 95% confidence intervals from meta-regression estimates are shown for each genotype. (**L**) Total number of inflammatory patches identified from whole quadriceps images (n = 4 wild-type, *mdx*, and *mdx*^TG^ quadriceps).

Immunofluorescence analysis revealed a reduced density of both CD68^+^ and CD206^+^ (*Mrc1*)^75^ macrophages in *mdx*^TG^ muscle compared to *mdx* (**Fig. 3B-D**), whereas total whole-muscle macrophage abundance and macrophage composition, assessed by flow cytometry, was largely preserved in *mdx*^TG^ relative to *mdx* (**Fig. 3E-H**). To determine if these findings reflected differences in macrophage infiltration patterns, we quantified macrophage spatial distribution across quadriceps cross-sections, revealing that macrophages in *mdx* muscle were concentrated within large, highly clustered inflammatory lesions, whereas *mdx*^TG^ muscle exhibited diffuse CD68^+^ macrophage distribution with a significantly reduced number of macrophage aggregates (**Fig. 3I-L**). Thus, SSPN expression reorganizes the inflammatory landscape rather than simply reducing immune cell infiltration. Consistent with this spatial reorganization, regions enriched for macrophage and pro-fibrotic macrophage gene expression remained associated with collagen type I staining and elevated *Col1a1* and *Col1a2* expression in both dystrophin-deficient genotypes, indicating that the association between macrophage activity and matrix remodeling is preserved even as macrophages shift from a clustered to a diffuse spatial distribution (**Fig. S4, Fig. S6**). These findings indicate that SSPN does not eliminate macrophage-associated remodeling activity but instead results in the redistribution of inflammatory cells from dense necrotic lesions into a more diffuse interstitial pattern.

Collectively, these observations demonstrate that the inflammatory landscape is reorganized rather than resolved in *mdx*^TG^ muscle. Whereas *mdx* muscle contains dense macrophage-rich inflammatory lesions, macrophages in *mdx*^TG^ muscle are distributed more diffusely, with substantially fewer focal aggregates despite continued macrophage-associated matrix remodeling.

### Fibro-adipogenic progenitors maintain collagen production despite reduced inflammatory pathology

To determine how the reorganized inflammatory environment influences ECM-producing stromal cells, we next examined FAPs using scRNA-seq. FAPs were the predominant source of fibrillar collagen expression in all genotypes, with highest expression in *mdx* FAPs (**Fig. 4A**). The strongest differentially expressed genes in *mdx* FAPs included *Postn, Thbs4,* and *Tnc*, consistent with fibroblast activation and differentiation^76–78^ (**Fig. 4B**). *mdx*^TG^ FAPs similarly maintained high *Postn* expression relative to wild-type FAPs, although reduced compared to *mdx* (**Fig. 4C-D, Fig. S5G**). Pathway analysis identified collagen-containing ECM as the most significantly enriched Gene Ontology term in both *mdx* and *mdx*^TG^ FAPs compared with wild type (**Fig. 4E-F**). FAPs comprised a similar proportion of total muscle-resident cells across wild-type, *mdx*, and *mdx*^TG^ muscle (**Fig. 4G**). Thus, persistent ECM remodeling in *mdx*^TG^ muscle is associated with sustained matrix-producing programs within FAPs rather than an increased relative representation of this population.

**Figure 4.**
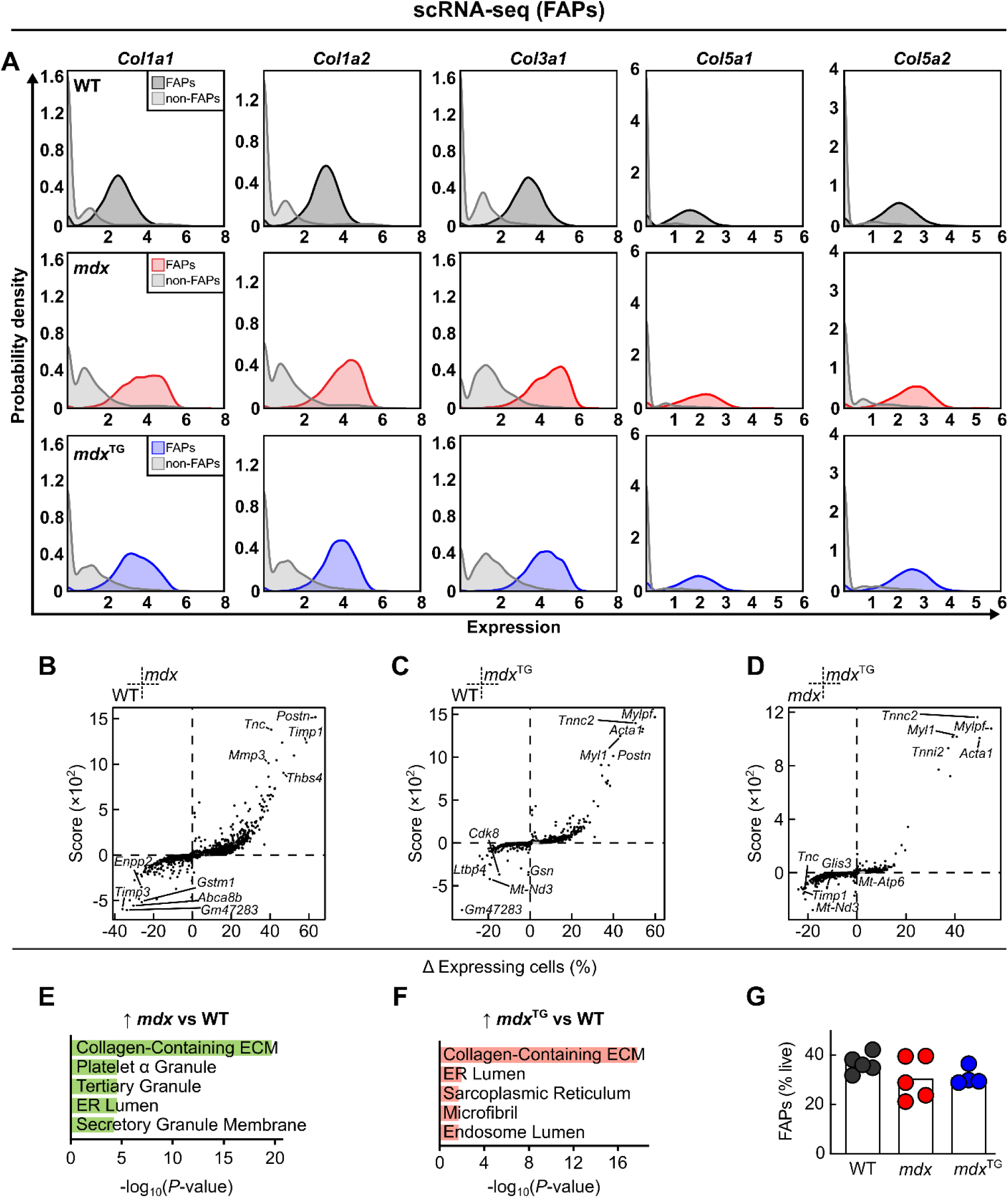
Single-cell analysis identifies FAPs as the primary source of fibrillar collagen expression in *mdx* and *mdx*^TG^ muscle. (**A**) Fibrillar collagen gene expression distributions across wild-type, *mdx*, and *mdx*^TG^ from scRNA-seq. FAPs exhibit predominant expression of *Col1a1*, *Col1a2*, *Col3a1*, *Col5a1*, and *Col5a2* compared to other cell types, highlighting their central role in collagen production within the muscle niche. DEGs between (**B**) wild-type and *mdx*, (**C**) wild-type and *mdx*^TG^, and (**D**) *mdx* and *mdx*^TG^. The x-axis represents the difference in the percentage of expressing cells and the y-axis is the product of -log_10_(adjusted *P*-value) and log_2_(FC). (**E**) GO term enrichment analysis of genes upregulated and enriched in FAPs from *mdx* versus wild-type and (**F**) *mdx*^TG^ versus wild-type comparisons. (**G**) Flow cytometric quantification of FAP abundance in wild-type (n = 5), *mdx* (n = 5) and *mdx*^TG^ (n = 3) muscle (expressed as a percentage of live cells).

### Collagen-rich *mdx* and *mdx*^TG^ muscles represent distinct extracellular matrix states

To determine whether persistent ECM transcription in *mdx*^TG^ muscle is reflected at the protein level, we quantified the muscle matrisome using a previously validated quantitative proteomic approach using radiolabeled Quantitative conCATamers (QconCATs) as internal standards for absolute quantification of ECM proteins^30,79^. Total fibrillar collagens, basement membrane proteins, FACIT collagens, matricellular proteins, and structural ECM proteins were all increased in *mdx*^TG^ relative to *mdx* muscle (**Fig. 5A-B**). Additional proteomic analyses revealed extensive remodeling of basement membrane-associated proteins, ECM-cytoskeletal interfaces, and secreted ECM modulators (**Fig. 5A-B**, **Fig. S7-S15**). Thus, improved dystrophic pathology in *mdx*^TG^ muscle occurs despite extensive ECM remodeling and greater accumulation of multiple matrix components, demonstrating that ECM abundance is not proportional to disease severity.

**Figure 5.**
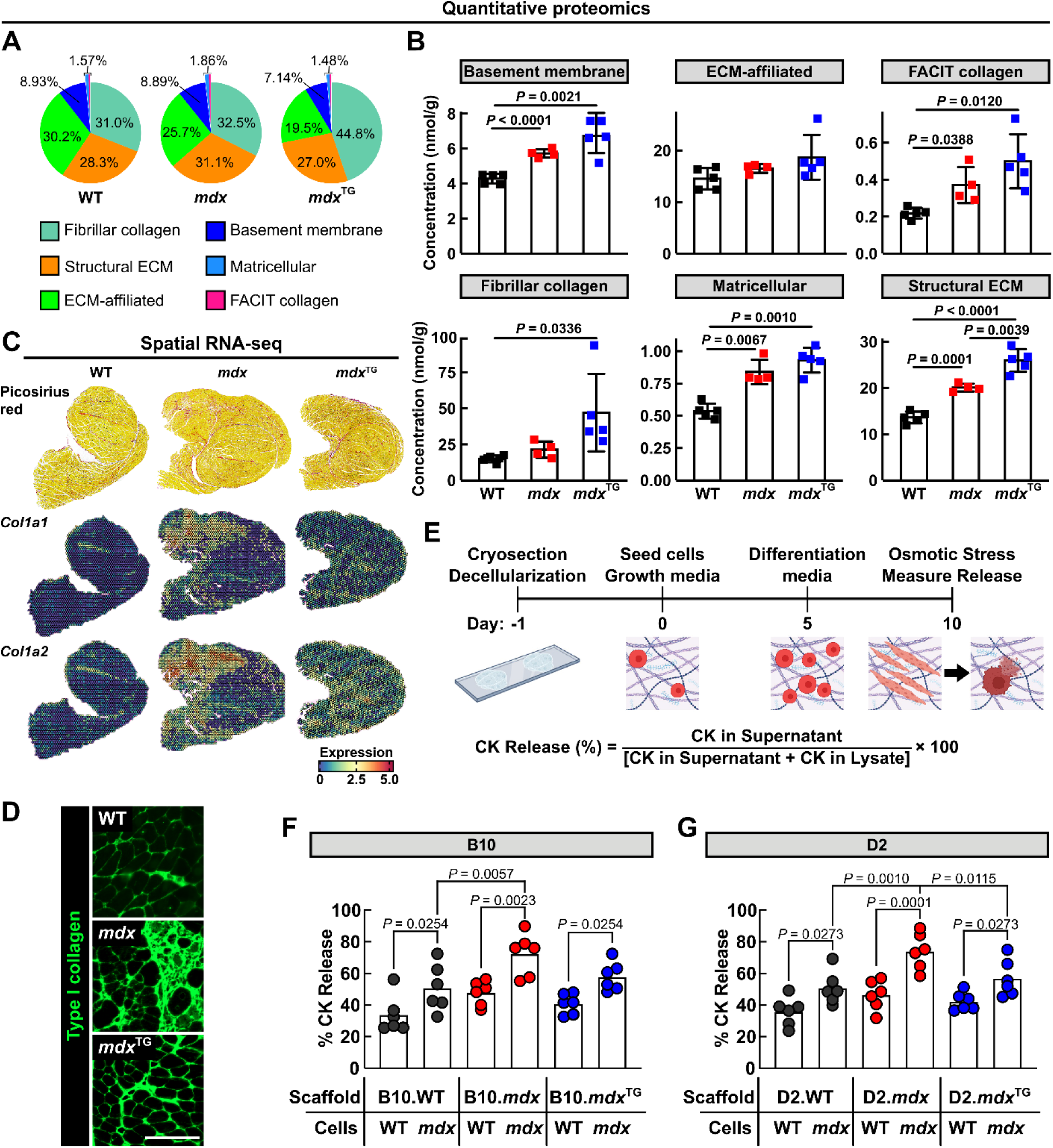
Greater collagen abundance in *mdx*^TG^ muscle yields a diffuse, membrane-protective matrix rather than dense scar. Despite increased collagen abundance, *mdx*^TG^ muscle lacks the dense inflammatory scars characteristic of *mdx* muscle. (**A**) Proportions of ECM protein functional classes in wild-type, *mdx*, and *mdx*^TG^ quadriceps muscle. (**B**) Total abundance of ECM proteins in six functional classes in wild-type, *mdx*, and *mdx*^TG^ quadriceps muscle. (**C**) Picrosirius red (PSR) and spatial RNA-seq analysis showing *Col1a1* and *Col1a2* expression in wild-type, *mdx*, and *mdx*^TG^ quadriceps muscle. (**D**) Type I collagen staining in wild-type, *mdx*, and *mdx*^TG^ quadriceps muscle. Scale bar, 200 μm. Statistical analysis by permutation test with maxT *P*-value adjustment. (**E**) Overview of experimental design for culturing wild-type or dystrophic H2K cells on WT, *mdx,* or *mdx*^TG^ myoscaffolds followed by osmotic stress and creatine kinase (CK) quantification. Healthy and dystrophic H2K cells cultured on (**F**) B10 myoscaffolds and (**G**) D2 myoscaffolds. Each data point represents the mean of triplicates of a single well. Statistical analysis by linear model with pairwise comparisons and Benjamini-Hochberg correction.

We next queried differences in the spatial organization of the ECM in *mdx* and *mdx*^TG^ muscle. Picrosirius red staining demonstrated that collagen deposition in *mdx* muscle was concentrated within inflammatory lesions, whereas *mdx*^TG^ muscle exhibited broad, diffuse ECM deposition together with more uniform *Col1a1/Col1a2* expression and a notable reduction of dense, pathological fibrotic scarring (**Fig. 5C-D**). Consistent with these observations, GO term analysis of *Col1a1*⁺ regions revealed enrichment of macrophage activation pathways in *mdx* but not *mdx*^TG^ muscle, indicating that collagen deposition in *mdx* muscle remains tightly associated with focal inflammatory lesions, whereas *mdx*^TG^ matrix remodeling occurs in a substantially less inflammatory environment (**Fig. S16**). Thus, collagen abundance and pathological scarring are distinguishable: increased collagen can reflect diffuse matrix remodeling rather than dense inflammatory scar formation, indicating that collagen abundance alone does not differentiate pathological fibrosis from ECM remodeling. Although *mdx*^TG^ muscle contains greater total collagen than *mdx* muscle, its matrix lacks the dense inflammatory scars that typify dystrophic muscle pathology, indicating that *mdx* and *mdx*^TG^ muscles represent distinct ECM states rather than different degrees of the same fibrotic response.

To determine whether these structurally distinct matrices differ in their biological consequences, we differentiated wild-type and *mdx* myoblasts into myotubes on decellularized *mdx*-derived scaffolds from two genetic backgrounds, C57BL/10 and the more severely fibrotic DBA/2J, and subjected myotubes to osmotic stress. *mdx* cells were more susceptible to stress on *mdx* scaffolds relative to wild-type or *mdx*^TG^ scaffolds in both backgrounds, despite the *mdx*^TG^ matrix containing more collagen than *mdx* matrix (**Fig. 5E–G**). Thus, the pathological consequences of the dystrophic matrix depend not only on collagen abundance, but also on its spatial organization and biological activity.

### YAP/TAZ signaling is enriched in fibrotic regions and maintained in dystrophic FAPs

SSPN expression alters ECM architecture and mechanotransduction signaling in dystrophic muscle^30^. Because ECM-derived mechanical cues regulate Hippo pathway signaling through focal adhesions^80,81^, and YAP/TAZ promotes fibroblast activation and ECM production^82–84^, we next asked whether fibrotic regions were enriched for YAP/TAZ-associated transcripts. Consistent with this, we observed increased expression of focal adhesion- and Hippo pathway-associated genes^30^ in *mdx* and *mdx*^TG^ muscle relative to wild-type (**Fig. S17**). Given the established role of the Hippo pathway in regulating fibroblast activation and ECM protein production^85^, we next examined whether highly fibrotic and inflamed regions were also enriched for YAP/TAZ target genes. Spatial transcriptomics revealed higher YAP/TAZ gene set scores in *mdx* and *mdx*^TG^ relative to wild-type (**Fig. 6A-F**). Inflammatory signatures were reduced in *mdx*^TG^ compared to *mdx*, whereas fibrotic signatures remained highest in *mdx*^TG^. In *mdx* muscle, inflammatory, fibrotic, and YAP/TAZ-associated transcriptional programs were concentrated within discrete lesions. In contrast, *mdx*^TG^ muscle exhibited a more diffuse spatial distribution with substantially fewer high-density foci, resulting in significantly smaller inflamed and fibrotic regions (**Fig. 6G, Fig. S18-S19**). Together, these data demonstrate that localized, inflammatory lesions coincide with strong fibrotic and YAP/TAZ-associated transcriptional programs in *mdx* muscle, whereas *mdx*^TG^

**Figure 6.**
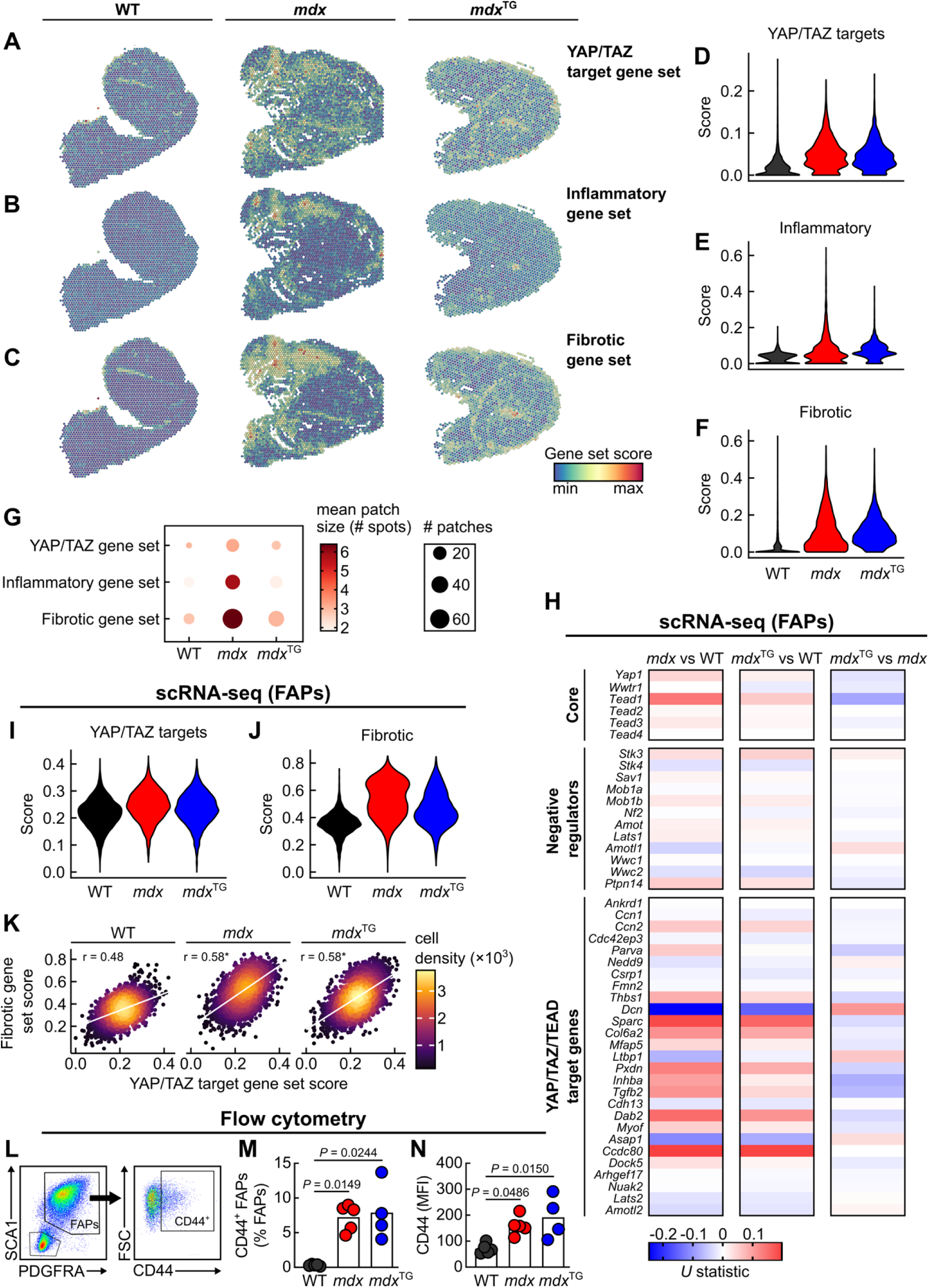
Hippo mediators are increased in *mdx* and *mdx*^TG^ FAPs. (**A**) Spatial RNA-seq showing gene set scores for YAP/TAZ target genes, (**B**) inflammatory genes, and (**C**) fibrotic genes in 15-week cross-sections of wild-type, *mdx*, *mdx*^TG^ quadriceps muscles. Violin plots of gene sets scores for (**D**) YAP/TAZ target genes, (**E**) inflammatory genes and (**F**) fibrotic genes. (**G**) Mean patch size (number of spots per patch) and number of patches of spatially connected, significant spots (FDR < 0.05 local Moran’s I) for YAP/TAZ, inflammatory, or fibrotic gene set scores. (**H**) Heatmap of YAP/TAZ pathway genes in FAPs from scRNA-seq. Heatmap color scale represents Wilcoxon rank-sum *U*-statistic. (**I**) Violin plot of gene set scores for YAP/TAZ target genes and (**J**) fibrotic genes in FAPs from scRNA-seq. (**K**) Correlation between gene set scores for YAP/TAZ target genes and fibrotic genes in FAPs from scRNA-seq. *Indicates statistically significantly different Pearson correlation coefficient compared to wild-type. Statistical analysis using Fisher r-to-z transformation followed by Holm multiple-comparisons correction. (**L**) Representative flow cytometry panel for FAPs selection and CD44 gating. (**M**) Percentage of CD44^+^ FAPs (relative to total FAPs) and (**N**) CD44 mean fluorescence intensity (MFI) in wild-type (n = 5), *mdx* (n = 5), *mdx*^TG^ (n = 4) muscle. Statistical analysis by permutation test.

exhibits a more diffuse distribution of these signatures despite persistent ECM remodeling. scRNA-seq analysis revealed that *Yap1* and *Ccn2* were expressed in FAPs across all genotypes (**Fig. S20**), and YAP/TAZ/TEAD core genes, target genes, and fibrotic genes were most highly expressed in *mdx* FAPs, with *mdx*^TG^ FAPs retaining elevated fibrotic gene expression relative to wild-type (**Fig. 6H-J**). Because many YAP/TAZ target genes also participate in ECM remodeling, we examined the relationship between these transcriptional programs. Correlation between YAP/TAZ target genes and fibrotic genes was stronger in both *mdx* and *mdx*^TG^ than in wild-type FAPs (**Fig. 6K**). Lastly, we confirmed higher CD44 expression, a cell adhesion protein upstream of YAP/TAZ activation^86,87^, in *mdx* and *mdx*^TG^ FAPs by flow cytometry (**Fig. 6L-N**). Taken together, these findings show that YAP/TAZ-associated signaling remains elevated in dystrophin-deficient muscle despite marked differences in the spatial organization of inflammatory and fibrotic pathology.

### A stiff dystrophic extracellular matrix promotes YAP-dependent fibroblast activation

To determine whether the mechanical properties of the dystrophic ECM are altered, we generated myoscaffolds from wild-type, *mdx*, and *mdx*^TG^ muscles and measured matrix stiffness by nanoindentation (**Fig. 7A-B, Fig. S21A-B**). The endomysium of both *mdx* and *mdx*^TG^ myoscaffolds exhibited significantly higher effective Young’s moduli than wild-type scaffolds, confirming increased matrix stiffness of the dystrophic ECM (**Fig. 7C**). To determine whether this altered ECM was sufficient to activate YAP signaling, wild-type FAPs were seeded onto myoscaffolds from each genotype in the presence or absence of verteporfin, which inhibits YAP nuclear localization^88^ (**Fig. 7D**). FAPs cultured on *mdx* and *mdx*^TG^ myoscaffolds exhibited increased nuclear YAP1 compared with those cultured on wild-type scaffolds, whereas verteporfin markedly reduced nuclear YAP1 localization on both dystrophic matrices (**Fig. 7E-F**).

**Figure 7.**
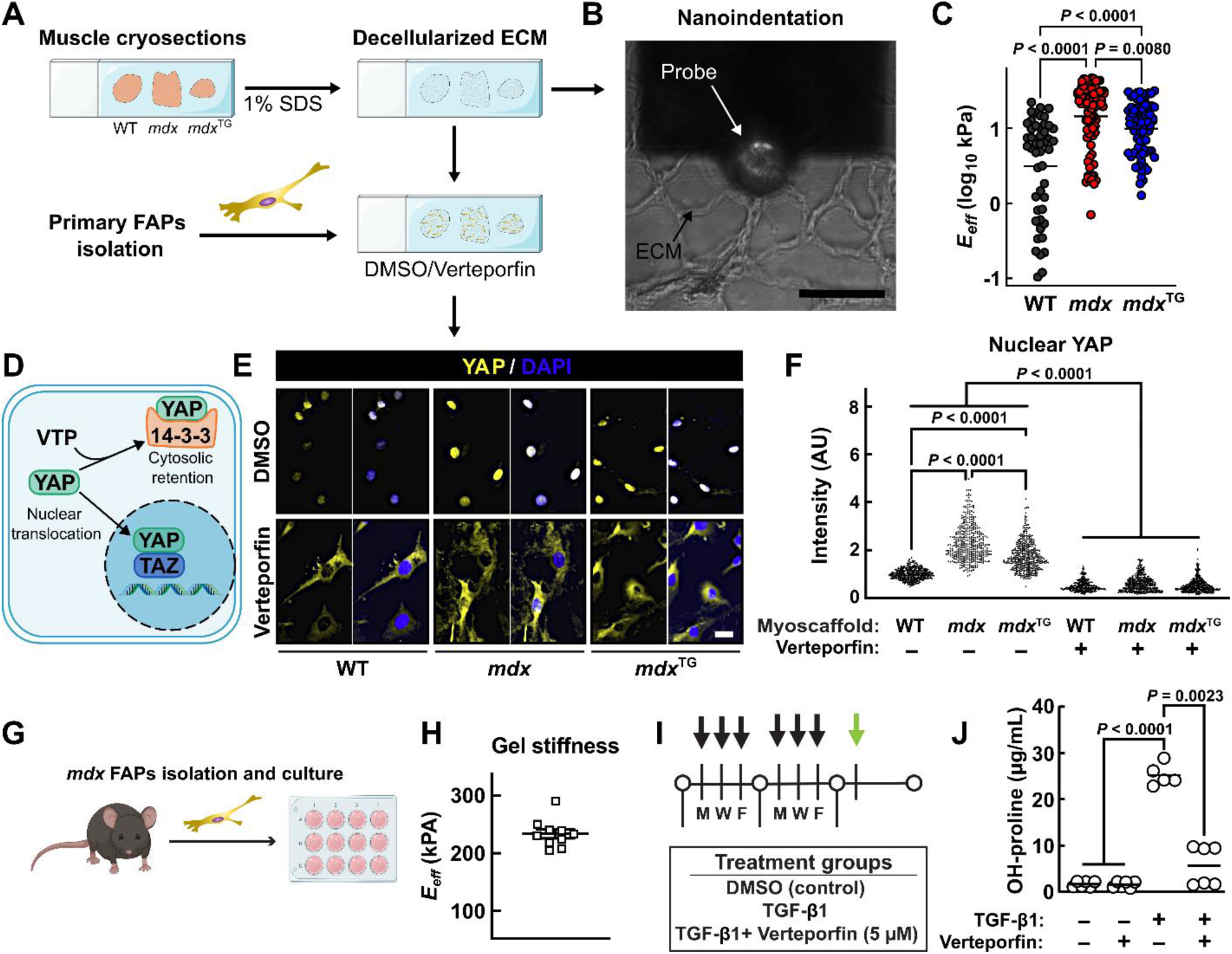
YAP1 inhibition reduces collagen production in *mdx* FAPs. (**A**) Schematic representation of myoscaffold generation and subsequent nanoindentation or primary FAPs culture. 50 μm wild-type, *mdx*, and *mdx*^TG^ quadriceps cryosections were decellularized using 1% SDS followed by (**B**) ECM stiffness measurements by nanoindentation. (**C**) Quantification of ECM stiffness measurements. *E_eff_*, effective Young’s modulus. Statistical analysis by permutation test. (**D**) Diagram of YAP nuclear localization and cytosolic retention with verteporfin treatment. Primary FAPs were isolated from wild-type mice and cultured on myoscaffolds (shown in (A)) with or without verteporfin, a pharmacological inhibitor of YAP1 nuclear localization. (**E**) Representative images of YAP1 staining in FAPs cultured on wild-type, *mdx*, and *mdx*^TG^ myoscaffolds after 24-hours of treatment with DMSO or verteporfin. Scale bar = 20 μm. (**F**) Quantification of nuclear YAP1 intensity in primary FAPs. Myoscaffolds were generated from n=4 mice per genotype. Results represent the average fluorescence intensity (in arbitrary units, a.u.) of 100 cells analyzed per biological replicate (representing one myoscaffold). Statistical analysis by generalized linear model with cluster-robust standard errors, followed by Tukey’s *post hoc* test. (**G**) Schematic diagram of experimental design for *in vitro* collagen production and quantification. (**H**) Stiffness measurements of synthetic gel coating on 6 well plates. (**I**) Overview of FAPs treatment strategy and *in vitro* generation of collagen scaffolds. FAPs were treated with DMSO, TGF-β1, verteporfin, or verteporfin and TGF-β1 for two weeks. Hydroxyproline assay was performed the week following the final treatment, as indicated by the green arrow. (**J**) Quantification of deposited hydroxyproline content (μg/mL). Statistical analysis by permutation test with step-down maxT *P*-value adjustment.

Because macrophages are key drivers of dystrophic pathology and their inflammatory response is influenced by mechanotransduction and YAP signaling^89^, we next asked whether myoscaffolds influence macrophages *in vitro*. Bone marrow-derived macrophage culture on myoscaffolds induced genotype-dependent differences in morphology and phagocytic capacity (**Fig. S21C-E**). In contrast to FAPs, macrophage nuclear YAP1 localization was unchanged across scaffold genotypes (**Fig. S22**), suggesting that ECM stiffness preferentially regulates YAP signaling in stromal rather than immune cells, and the observed functional effects occur independently of YAP signaling in macrophages.

Given the effect of myoscaffold genotype on YAP signaling in FAPs, and because YAP signaling regulates fibroblast activation and ECM production^18,90–92^, we next investigated whether pharmacological inhibition of YAP alters collagen deposition by dystrophic FAPs. *mdx* FAPs were cultured on hydrogels mimicking the physiological stiffness of the *mdx* ECM^93^ (**Fig. 7G-H**) and treated with vehicle, TGF-β1, verteporfin, or TGF-β1 plus verteporfin three times per week for two weeks (**Fig. 7I**) after which deposited collagen was quantified by hydroxyproline assay. TGF-β1 significantly increased collagen deposition in *mdx* FAPs, whereas verteporfin markedly attenuated this response (**Fig. 7J**). Together, these findings demonstrate that a stiff dystrophic ECM is sufficient to promote nuclear YAP localization in FAPs and that pharmacologic YAP blockade suppresses collagen production even in the presence of TGF-β1. These results support a model in which matrix-derived mechanical signaling reinforces the collagen-producing state of dystrophic FAPs alongside canonical pro-fibrotic signaling.

### YAP blockade uncouples fibrosis from ongoing inflammatory signaling *in vivo*

To determine whether YAP1 signaling is required to maintain fibrosis *in vivo*, we treated *mdx* mice with 50 mg/kg verteporfin from 2 to 6 weeks of age (**Fig. 8A**), a dosage previously reported to reduce collagen deposition after acute muscle injury^18^. Treatment began at 2 weeks of age, while *mdx* muscle is still pre-necrotic, and continued through the period of peak necrosis, inflammation, and TGF-β1 release^15,16,94–99^, allowing us to test whether YAP1 inhibition is sufficient to limit collagen deposition during peak inflammation. Following 5 weeks of treatment, heart and hindlimb muscle masses were unchanged between treatment groups, indicating no detectable adverse effects on muscle growth^100^ (**Fig. 8B**). Since the diaphragm develops rapid and severe fibrosis in *mdx* mice, we measured hydroxyproline content and found diaphragms from verteporfin-treated mice had lower total collagen content (**Fig. 8C**). Additionally, type I collagen quantification in quadriceps muscles revealed a decrease in total collagen area as well as less collagen scarring (**Fig. 8D-F**). Together, these findings demonstrate that pharmacological YAP blockade limits collagen accumulation and scar formation during active dystrophic pathology, supporting a functional role for YAP-associated mechanotransduction in fibrotic remodeling *in vivo*. We next asked whether this mechanosensitive program is similarly engaged in human DMD disease.

**Figure 8.**
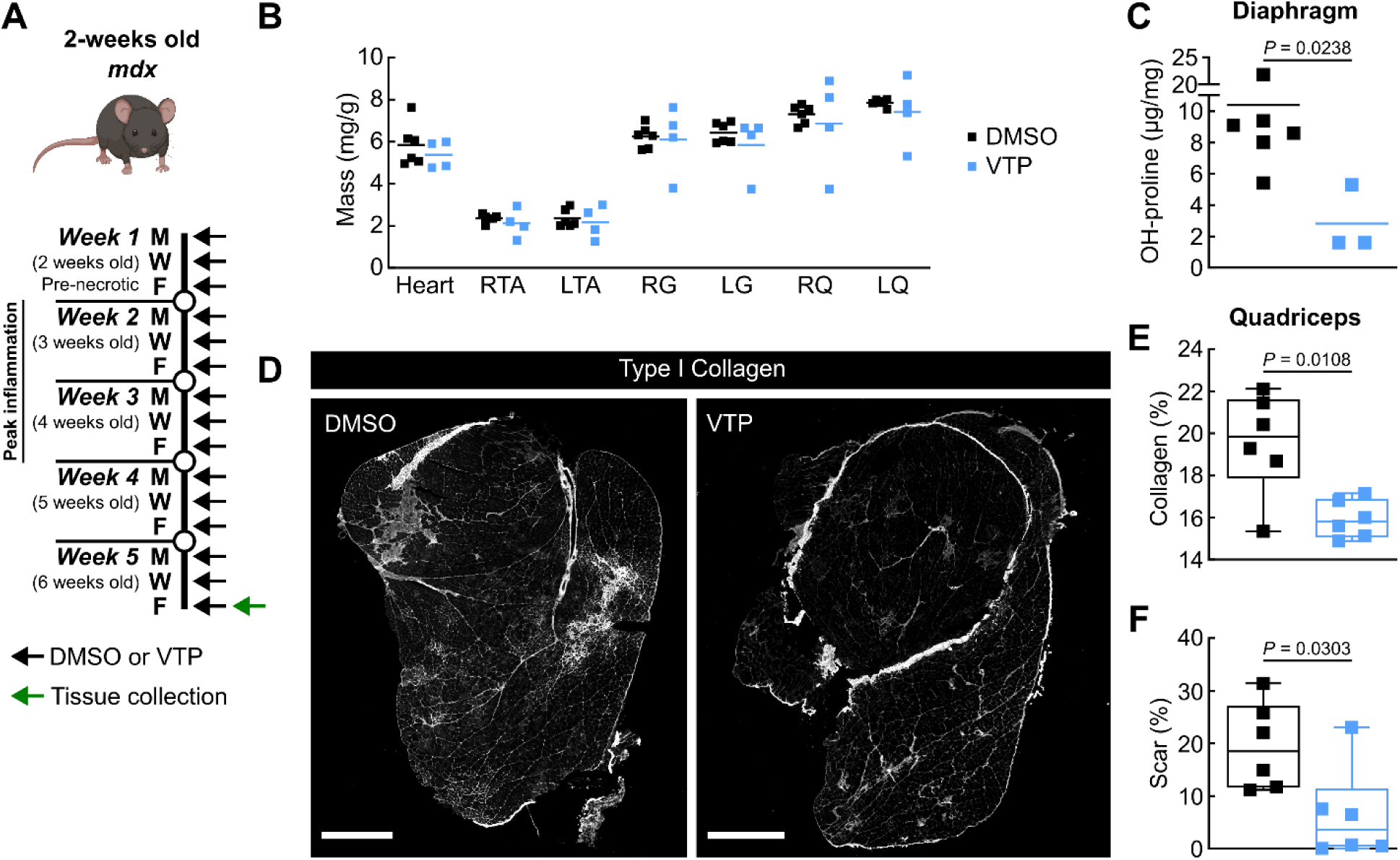
YAP blockade limits fibrosis during active dystrophic inflammation *in vivo*. (**A**) Schematic diagram of verteporfin treatment. Young (2-weeks old) *mdx* mice were intraperitoneally injected with 50 mg/kg verteporfin or DMSO 3× per week (black arrows) for 5 weeks. Muscle was collected at the end of the 5^th^ week of treatment (green arrow). (**B**) Relative heart and muscle mass (mg muscle per gram body mass) at the end of treatment (n = 6 DMSO, n = 4 verteporfin). RTA, right tibialis anterior; LTA, left tibialis anterior; RG, right gastrocnemius; LG, left gastrocnemius; RQ, right quadriceps; LQ, left quadriceps. (**C**) Hydroxyproline (OH-proline) content in diaphragm muscles following verteporfin treatment (n = 6 DMSO, n = 3 verteporfin). (**D**) Representative images of collagen type I staining in quadriceps muscles from DMSO and verteporfin (VTP) treated *mdx* mice. Scale bar, 1000 μm. (**E**) Quantification of type I collagen area. (**F**) Quantification of scar area as a percentage of the total collagen area (n = 6 per group). Statistical analysis by permutation test.

### Hippo pathway signaling remains elevated in DMD patient FAPs despite corticosteroid therapy

To determine whether Hippo pathway signaling is similarly altered in human disease, we analyzed publicly available single-cell RNA-seq data from Fernandez-Simon and colleagues^66^ and compared FAPs from DMD patients with healthy controls. Expression of *YAP1* and *WWTR1* (TAZ) was significantly elevated in DMD FAPs relative to control (**Fig. 9A-B**). Genes associated with FAP activation, including *PDGFRA*, *CD44*, mechanosensitive integrins, collagens, and collagen cross-linking enzymes, remained elevated despite ongoing corticosteroid therapy (**Fig. 9A-D**). In contrast, several inflammatory mediators, including *TGFB1*, chemokines, and cytokine receptors, were reduced (**Fig. 9A**), indicating that inflammatory suppression does not normalize the broader fibroblast activation program. Consistent with transcriptomic data, the proportion of nuclear YAP⁺PDGFRα⁺ cells was increased in DMD FAPs (**Fig. 9E-F**). Proteomic analyses likewise demonstrated persistent accumulation of fibrillar collagens in DMD muscle relative to healthy controls despite ongoing corticosteroid treatment (**Fig. 9G**). Together, these findings demonstrate that YAP/TAZ-associated activation and ECM remodeling persist in human DMD despite corticosteroid therapy and reduced expression of multiple inflammatory mediators. This persistence parallels the separation between inflammatory signaling and matrix remodeling observed in *mdx*^TG^ muscle and identifies YAP activation as a conserved feature of dystrophic fibrosis across mouse and human muscle (**Fig. 10**).

**Figure 9.**
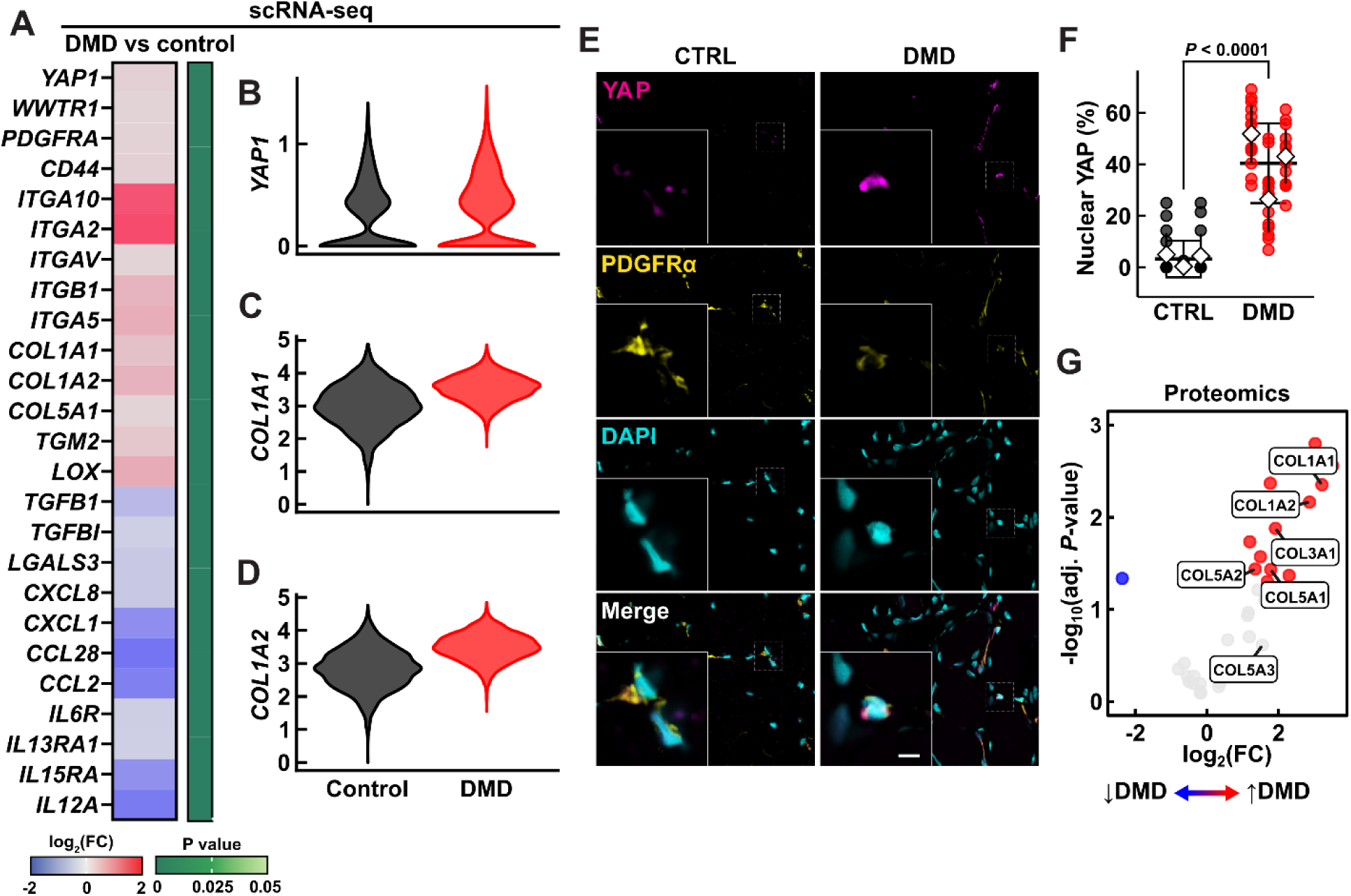
Elevated nuclear YAP in DMD FAPs. (**A**) Heat map showing differential gene expression in FAPs from DMD versus healthy individuals. Genes with an adjusted *P* < 0.05 were considered significantly differentially expressed. (**B**) Violin plot comparing *YAP1,* (**C**) *COL1A1*, and (**D**) *COL1A2* expression levels in FAPs from control and DMD. (**E**) Representative immunofluorescence images showing increased nuclear YAP1 localization in PDGFRA⁺ FAPs from DMD muscle biopsies. Scale bar, 50 μm. (**F**) Quantification of nuclear YAP1⁺ FAPs in control and DMD muscle biopsies (n = 3 per group). The percentage of FAPs with nuclear YAP1 in each captured image are represented as individual data points, with white diamonds indicating the mean of all captured images per independent patient sample. Statistical analysis by permutation test. (**G**) Volcano plot showing increased fibrillar collagen protein expression (log₂ fold change) in DMD patients relative to healthy controls. Statistical analysis by permutation test.

**Figure 10.**
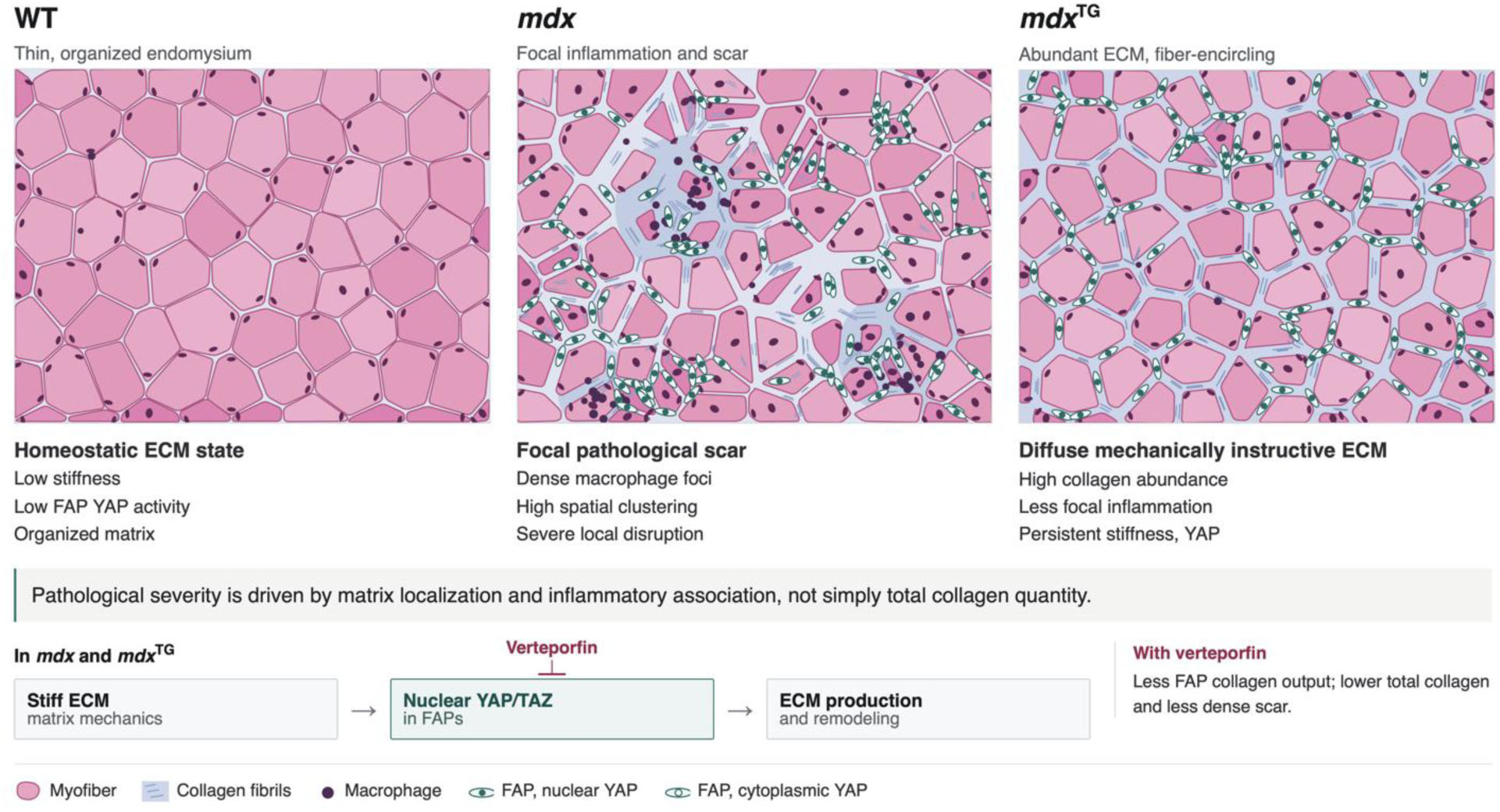
Bulk collagen abundance does not predict pathological fibrosis: matrix localization and inflammatory association define distinct dystrophic ECM states. Schematic of myofiber cross-sections, endomysial ECM containing collagen fibrils, and FAPs with nuclear or cytoplasmic YAP in wild-type, *mdx* and SSPN-transgenic *mdx* (*mdx*^TG^) muscle. Wild-type muscle shows a thin, organized endomysium, low matrix stiffness and predominantly cytoplasmic YAP in FAPs. In *mdx* muscle, collagen accumulates focally within macrophage-dense lesions, producing spatially clustered scar, severe local distortion of myofiber geometry and nuclear YAP in lesion-associated FAPs. In *mdx*^TG^ muscle, total collagen is high but deposited diffusely as a continuous fiber-encircling network with little focal inflammation, while matrix stiffness and nuclear YAP in FAPs persist. An equal or greater collagen burden therefore corresponds to markedly different pathological states, with severity tracking matrix localization and inflammatory association rather than total collagen quantity. Lower panel, proposed feed-forward loop in which stiff ECM promotes nuclear translocation of YAP/TAZ in FAPs and drives further ECM production and remodeling; verteporfin blocks YAP/TAZ activity, lowering FAP collagen output, total collagen and scar density.

## Discussion

Fibrosis is widely viewed as a downstream consequence of chronic injury and inflammation. Here, we show that fibrotic remodeling can persist as a mechanically reinforced tissue state even when inflammatory pathology is substantially reduced. The *mdx*^TG^ model uncouples collagen accumulation from pathological scar formation: despite extensive ECM accumulation, *mdx*^TG^ muscle lacks the dense macrophage-rich lesions characteristic of *mdx* muscle. Nevertheless, both dystrophin-deficient matrices remain stiff and induce nuclear YAP localization in FAPs. Pharmacological YAP blockade reduces collagen production in vitro and fibrosis in vivo, supporting a feed-forward model in which remodeled ECM reinforces mechanosensitive stromal activation.

In dystrophic muscle, inflammatory signaling, particularly TGF-β, has been considered the dominant driver of fibrosis through activation of FAPs^5–7,9,101–103^. However, because of the tight coupling between muscle degeneration, inflammation, and ECM deposition, it has remained unclear whether fibrosis is continuously driven by upstream signals or becomes self-reinforcing once established. By separating inflammatory pathology from fibrotic remodeling in the *mdx*^TG^ model, our findings reveal that collagen deposition can persist despite reduced injury-associated signaling, exposing a mechanosensitive program within FAPs that reinforces pro-fibrotic collagen production. This distinction reframes fibrosis not solely as a response to damage, but as a state shaped by ECM-dependent signaling alongside ongoing injury. In this context, *mdx*^TG^ muscle represents an adaptive ECM remodeling state that distinguishes persistent fibrotic remodeling from the severity of ongoing tissue injury and inflammatory pathology. Notably, a similar dissociation is evident in patients with DMD. In human DMD, *YAP1* and *WWTR1* expression and the broader FAP activation program remained elevated despite ongoing corticosteroid therapy, even as several inflammatory mediators were reduced, indicating that pharmacological suppression of inflammation does not normalize the underlying mechanosensitive fibrotic program.

Importantly, our findings challenge the use of collagen abundance alone as a surrogate for fibrosis severity. Although *mdx*^TG^ muscle contained greater total collagen than *mdx* muscle, this matrix lacked the dense inflammatory scars that typify dystrophic pathology. Collagen burden can therefore correspond to biologically distinct ECM states depending on its spatial organization, inflammatory landscape, and mechanical activity. These findings argue that pathological fibrosis should be defined not solely by the quantity of deposited matrix, but by the biological state that matrix creates and sustains.

Our findings extend prior work linking YAP/TAZ activity to fibrotic environments. YAP/TAZ regulates fibroblast activation and myofibroblast differentiation across tissues, often in coordination with TGF-β/SMAD signaling^18,83,85,104–109^. In skeletal muscle specifically, YAP activity is enriched in fibrotic regions and correlates with FAP activation, including co-localization with collagen-producing, pro-fibrotic gene expression^18^. However, these roles have largely been established in the context of ongoing injury and interpreted as downstream of inflammatory signaling. Here, we show that stiff dystrophic ECM is sufficient to promote nuclear YAP localization in FAPs, and that YAP blockade attenuates collagen production even in the presence of the pro-fibrotic cytokine TGF-β1, indicating that mechanical inputs act in parallel with, rather than merely downstream of, inflammatory signaling. Together, these findings support a model in which inflammatory cues initiate and amplify fibroblast activation, while ECM mechanics provide a reinforcing signal that sustains a collagen-producing state. This pro-fibrotic FAP state also persists *in vivo* even as inflammatory cues in *mdx*^TG^ muscle are reduced relative to *mdx*, positioning YAP as a convergence point between biochemical and mechanical inputs in fibrotic remodeling.

These observations have important implications for therapeutic strategies targeting fibrosis in DMD and related disorders. Current approaches primarily focus on modulating inflammation or inhibiting TGF-β signaling, but such strategies can disrupt regenerative processes and have shown limited durability^110–115^. Our findings suggest that therapies directed solely at inflammatory pathways may be insufficient once a mechanically reinforced fibrotic state has become established. Consistent with this concept, pharmacologic inhibition of YAP with verteporfin reduced collagen deposition *in vivo*, supporting mechanotransduction as a therapeutic target for limiting fibrosis.

Although verteporfin has recognized off-target activities, the convergence of ECM stiffness measurements, FAP mechanosignaling assays, pharmacological intervention in mice, and human DMD data implicates YAP-associated mechanotransduction in the maintenance of dystrophic fibrosis. More broadly, these findings suggest that effective anti-fibrotic strategies may require targeting not only the signals that initiate fibrosis, but also the mechanically reinforced stromal states that sustain it^18,83,85,116^.

This study has several limitations. Our verteporfin treatment experiments included male and female mice but did not directly examine sex differences. Homozygous female *mdx* mice display dystrophic pathology, and whether the ECM states and mechanosignaling responses described here differ by sex remains untested. Additionally, single-cell, single-nucleus, and spatial sequencing were performed on one animal per genotype, and we therefore treat these datasets as descriptive. The spatial reorganization of inflammatory pathology, however, was independently validated by whole-section imaging in four mice per genotype.

## Ethics declarations

RHC is a co-founder of Pagoda Bio and holds equity and a position on the scientific advisory board in the company, which is developing small-molecule sarcospan enhancers. RHC, EIM, GRM, and ACR are inventors on patents covering sarcospan-based technologies assigned to the University of California, Los Angeles. The remaining authors declare no competing interests.

## Funding

This study was supported by the National Institutes of Health (NINDS R01 NS120060 to SAV, NIGMS R01 GM143378 to EJD, NIH-NIGMS R35 GM 161448 to ACR, 5T32AR065972-08 to PK, DH, and JR, NIAID, T32 AI177324 to PF), the National Science Foundation (BRITE Fellow Award CMMI-2135747 to ACR) and a Hyde Fellowship (UCLA Department of Integrative Biology and Physiology) to PK and DH.

## Author contributions

PK, DH, and RHC conceived and designed the study. EJD provided single cell and statistical analysis expertise, VT and EM provided DMD clinical expertise as well as DMD animal model expertise, SAV and PF provided DMD inflammation expertise and conducted flow cytometry experiments, KCH provided global and matrisome proteomics expertise, and RHC provided SSPN and DMD pathophysiology expertise. WG, GRM, PK and DH performed bulk RNA sequencing analysis and prepared figures. PK, DH, and EIM conducted all *in vitro* and *in vivo* experiments. Force probe indentation measurements were carried out by PK with support from DQ. Human DMD and control biopsies were generously contributed by EM and VT and stained for mechanical signaling markers by TSF, DH, and PK. EIM, JW and DH contributed to mouse colony maintenance, genotyping, and collection of tissues and blood from all murine samples. DH developed isolation protocols for spatial, single-nucleus, and single-cell RNA sequencing. Computational analysis of scRNA-seq datasets was supported by EJD and performed by MHA, HT, PK, and DH. DH analyzed scRNA-seq, snRNA-seq, and spatial RNA-seq datasets. DH and TSF adapted the protocol for FAP isolation and culture. DH adapted the hydroxyproline assay, and DH, PK and EIM performed OH-proline quantification. PK contributed to the immunofluorescence staining and quantification in Figs. 7 and 9, and DH conducted all statistical analysis under EJD’s guidance. KSR contributed to the original development of the muscle decellularization protocols and worked with RT on the collection of human control and DMD biopsies; proteomics was performed by MM and KCH. PK, DH, and RHC wrote the manuscript with input from all authors. All authors reviewed and approved the final version of the manuscript.

