## Supplementary Information for "Distinct extracellular matrix states uncouple collagen accumulation from pathological fibrosis in Duchenne muscular dystrophy"

**Supplemental Table 1. Sarcospan improves dystrophic muscle pathology and function despite persistent extracellular matrix accumulation.**

| System | Finding (magnitude relative to *mdx*) | Evidence (assay, muscle, age) | Citation |
|---|---|---|---|
| **Dystrophic pathology is broadly improved** | | | |
| Systemic | Reduced muscle damage marker: serum creatine kinase reduced by roughly 60% (~1,200 to ~500 U/L; WT ~250 U/L) reflecting improved membrane stability | Serum creatine kinase, 20 wk | 27, 28 |
| Skeletal muscle | Improved sarcolemmal integrity: EBD-positive fibers reduced from ~10% to ~1.5%, not significantly different from WT | Evans blue dye uptake, quadriceps, 6 wk | 27, 28 |
| Skeletal muscle | Reduced degeneration and regeneration cycling: central nuclei reduced from ~50% to ~23% (hSSPN, 4 to 6 wk), ~48% to ~19% (hSSPN, 6 wk), ~75% to ~20% (hSSPN, 10 to 13 wk; $P < 0.0001$) and ~45% to ~17% (mSSPN, 6 wk). eMHC-positive fibers trend lower | Central nucleation and eMHC immunofluorescence, H&E, quadriceps, 4 to 13 wk | 28, 29, 32, 33 |
| Skeletal muscle | Normalized fiber size distribution: peak cross-sectional area shifts from 500 to 1,000 μm² and mean area falls from 2,200–2,250 μm² to 1,200–1,400 μm², with loss of the small-regenerating and large-hypertrophic tails. Transgenic mice weigh less than non-transgenic littermates | Myofiber cross-sectional area binned per 500 μm², quadriceps, 10 to 12 wk; body weight, 10 to 11.5 wk | 29 |
| Skeletal muscle | Improved sarcolemmal integrity by a second, dye-independent measure: IgG-positive fibers reduced from ~3.3% to ~0.7% ($P < 0.0001$), indistinguishable from WT | Serum IgG infiltration, immunofluorescence, quadriceps, 10 to 13 wk | 29 |
| Skeletal muscle | Regenerative reserve is preserved: Pax7-positive satellite cell density comparable to WT, so reduced | Pax7 immunofluorescence, cells per myofiber, | 29 |

| System | Finding (magnitude relative to *mdx*) | Evidence (assay, muscle, age) | Citation |
|---|---|---|---|
| | regeneration reflects less damage rather than stem cell depletion | quadriceps, 10 to 13 wk | |
| Skeletal muscle | Increased resistance to contraction-induced injury: ~95% of initial force retained after five eccentric contractions vs ~66% in *mdx*, comparable to WT. Isometric specific force is not improved | Eccentric contraction protocol, isolated EDL, 5 mo | 28 |
| Skeletal muscle | Reduced fatigue following exercise: post-exercise activity restored to WT levels, ~4-fold greater distance travelled than *mdx* over 6 min. Sarcolemmal nNOS is not restored | Open-field activity after grip-strength exercise, 6 mo | 28 |
| Whole animal | Increased strength: forelimb grip force approximately 2-fold over *mdx* on the fifth consecutive trial (~0.013 to ~0.026 N/g), remaining below WT (~0.06 N/g) | Forelimb grip strength normalized to body weight, 6 wk | 28 |
| Whole animal | Improved ambulation after grip challenge: distance travelled not significantly different from WT; no genotype difference in unexercised mice | Open-field video tracking, 6 min (same protocol as the fatigue row above), 6 mo | 28 |
| Diaphragm | Reduced dystrophic histopathology: thickness normalized to WT (~520 to ~240 µm; $P < 0.0001$), central nuclei ~62% to ~28%, EBD-positive fibers ~45% to ~11% | H&E, Masson trichrome, EBD uptake, diaphragm, 6 wk | 28 |
| Pulmonary | Improved ventilatory function under hypercapnia: minute ventilation (~142 to ~154 mL/min; WT ~176) and peak expiratory flow (~8.2 to ~9.3 mL/s; WT ~9.8) increased. Respiratory frequency and peak inspiratory flow are not improved | Whole-body plethysmography, 8% $CO_2$ challenge, 26 wk | 28 |
| Cardiac | Improved sarcolemmal integrity: EBD-positive cardiomyocytes | Evans blue dye uptake, heart, 13 to 15 mo | 67 |

| System | Finding (magnitude relative to *mdx*) | Evidence (assay, muscle, age) | Citation |
|---|---|---|---|
| | reduced from 13.3% to 1.3%, restored to WT levels | | |
| Cardiac | Reduced myocardial damage and fibrosis: decreased collagen and fat deposition; serum CK-MB reduced ($P = 0.009$); ANP transcript reduced ($P = 0.036$) | H&E, Masson trichrome, Oil Red O, serum CK-MB, ANP qRT-PCR, 4 mo | 67, 68 |
| Cardiac | Improved systolic performance: ESPVR slope +16.3%. By echocardiography, ejection fraction 59.1 ± 5.0 to 71.7 ± 4.9% and fractional shortening 27.4 ± 1.3 to 36.4 ± 3.4%, both restored to WT | Hemodynamic pressure-volume analysis (2019); echocardiography, 10 to 11 mo (2015) | 67, 68 |
| Cardiac | Reduced hypertrophic remodeling: heart weight/body weight 6.17 ± 0.15 to 5.42 ± 0.25 mg/g (WT 5.33 ± 0.11); LV mass 81.4 ± 8.8 to 65.8 ± 7.0 mg (trend, not significant) | Heart weight/body weight and echocardiographic LV mass, 10 to 12 mo | 67 |
| Cardiac | Partially restored beta-adrenergic responsiveness. SSPN loss blunts the isoproterenol response (ejection fraction 64.5 ± 3.8% vs 80.6 ± 6.9% in WT), with increased LV mass (116.1 ± 4.8 vs 82.6 ± 8.7 mg) and fibrosis (12% vs 4.6% of area) | Isoproterenol 0.8 mg/day for 2 wk, echocardiography and histology, 12 mo | 67, 68 |

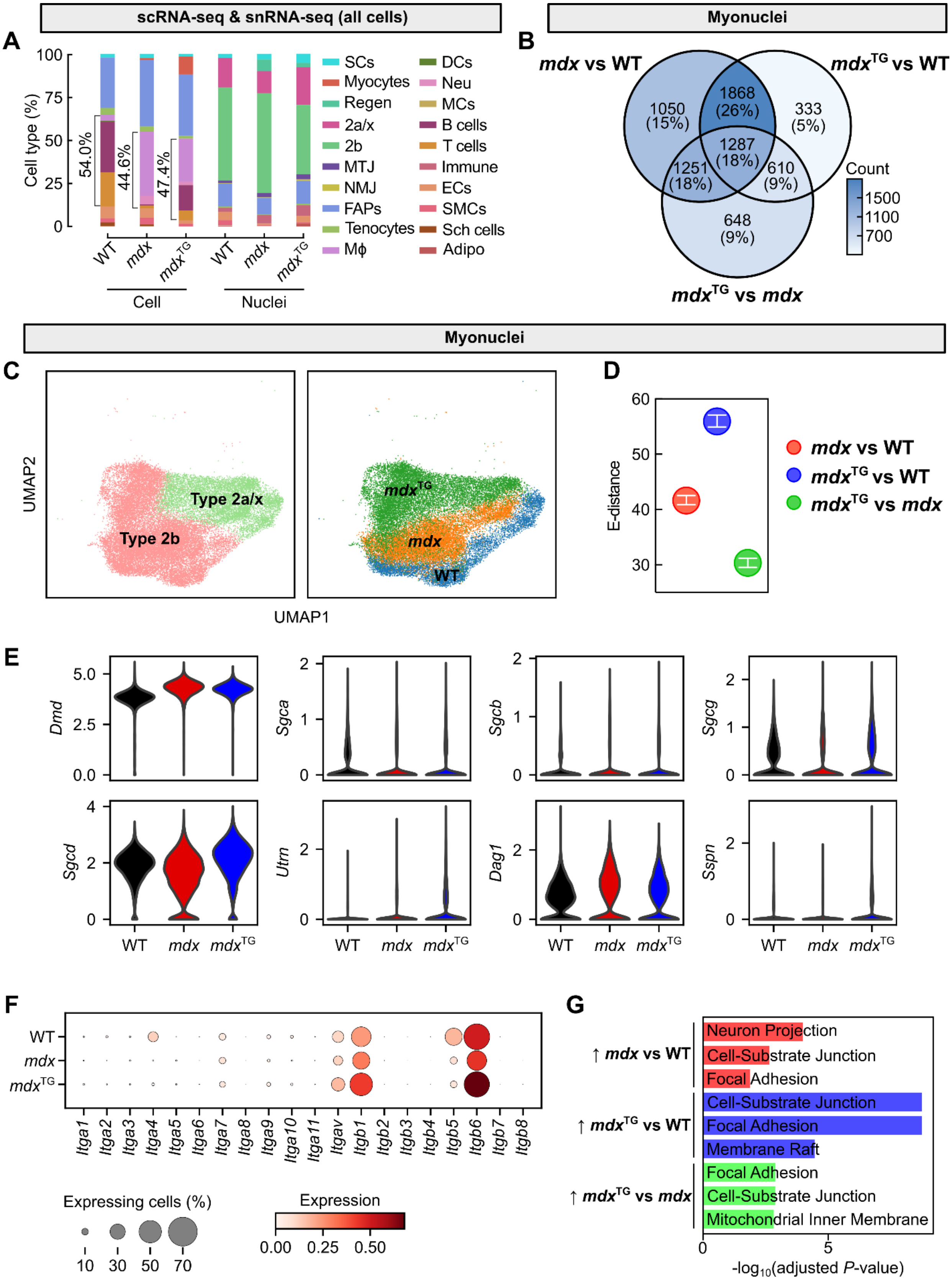
scRNA-seq & snRNA-seq (all cells)
Myonuclei
A
B
Cell type (%)
54.0%
44.6%
47.4%
WT
mdx
mdxTG
Cell
Nuclei
SCs
Myocytes
Regen
2a/x
2b
MTJ
NMJ
FAPs
Tenocytes
Mϕ
DCs
Neu
MCs
B cells
T cells
Immune
ECs
SMCs
Sch cells
Adipo
mdx vs WT
mdxTG vs WT
mdxTG vs mdx
1050 (15%)
1868 (26%)
333 (5%)
1287 (18%)
1251 (18%)
610 (9%)
648 (9%)
Count
Myonuclei
C
D
UMAP2
UMAP1
Type 2a/x
Type 2b
mdxTG
mdx
WT
E-distance
mdx vs WT
mdxTG vs WT
mdxTG vs mdx
E
Dmd
Sgca
Sgcb
Sgcg
Sgcd
Utrn
Dag1
Sspn
F
G
Itga1
Itga2
Itga3
Itga4
Itga5
Itga6
Itga7
Itga8
Itga9
Itga10
Itga11
Itgav
Itgb1
Itgb2
Itgb3
Itgb4
Itgb5
Itgb6
Itgb7
Itgb8
Expressing cells (%)
Expression
↑ mdx vs WT
↑ mdxTG vs WT
↑ mdxTG vs mdx
Neuron Projection
Cell-Substrate Junction
Focal Adhesion
Cell-Substrate Junction
Focal Adhesion
Membrane Raft
Focal Adhesion
Cell-Substrate Junction
Mitochondrial Inner Membrane
-log10(adjusted P-value)

**Supplemental Figure 1. SSPN increases focal adhesion gene expression in myonuclei.** (**A**) Proportion of cell types identified by single cell (scRNA-seq) and single nucleus (snRNA-seq) RNA sequencing in wild-type, *mdx*, and *mdx*$^{TG}$ muscle. SCs, satellite cells; Regen, regenerating myonuclei; MTJ, myotendinous junction; NMJ, neuromuscular junction; FAPs, fibroadipogenic progenitors; M$\phi$, macrophages, DCs, dendritic cells; Neu, neutrophils; MCs, mast cells; ECs, endothelial cells; SMCs, smooth muscle cells; Sch cells, Schwann cells; Adipo, adipocytes. (**B**) Venn diagram of differentially expressed genes between myonuclei annotated in snRNA-seq. (**C**) UMAP visualization of myonuclei separated by type 2a/2x and type 2b and genotype. (**D**) Energy distances calculated on myonuclei. Error bars represent 95% confidence interval determined by bootstrapping. (**E**) Dystrophin-glycoprotein complex gene expression in all wild-type, *mdx*, and *mdx*$^{TG}$ myonuclei. (**F**) Dotplot of integrins in all wild-type, *mdx*, and *mdx*$^{TG}$ myonuclei**.** (**G**) GO-term enrichment (cellular component) from wild-type, *mdx*, and *mdx*$^{TG}$ myonuclei**.**

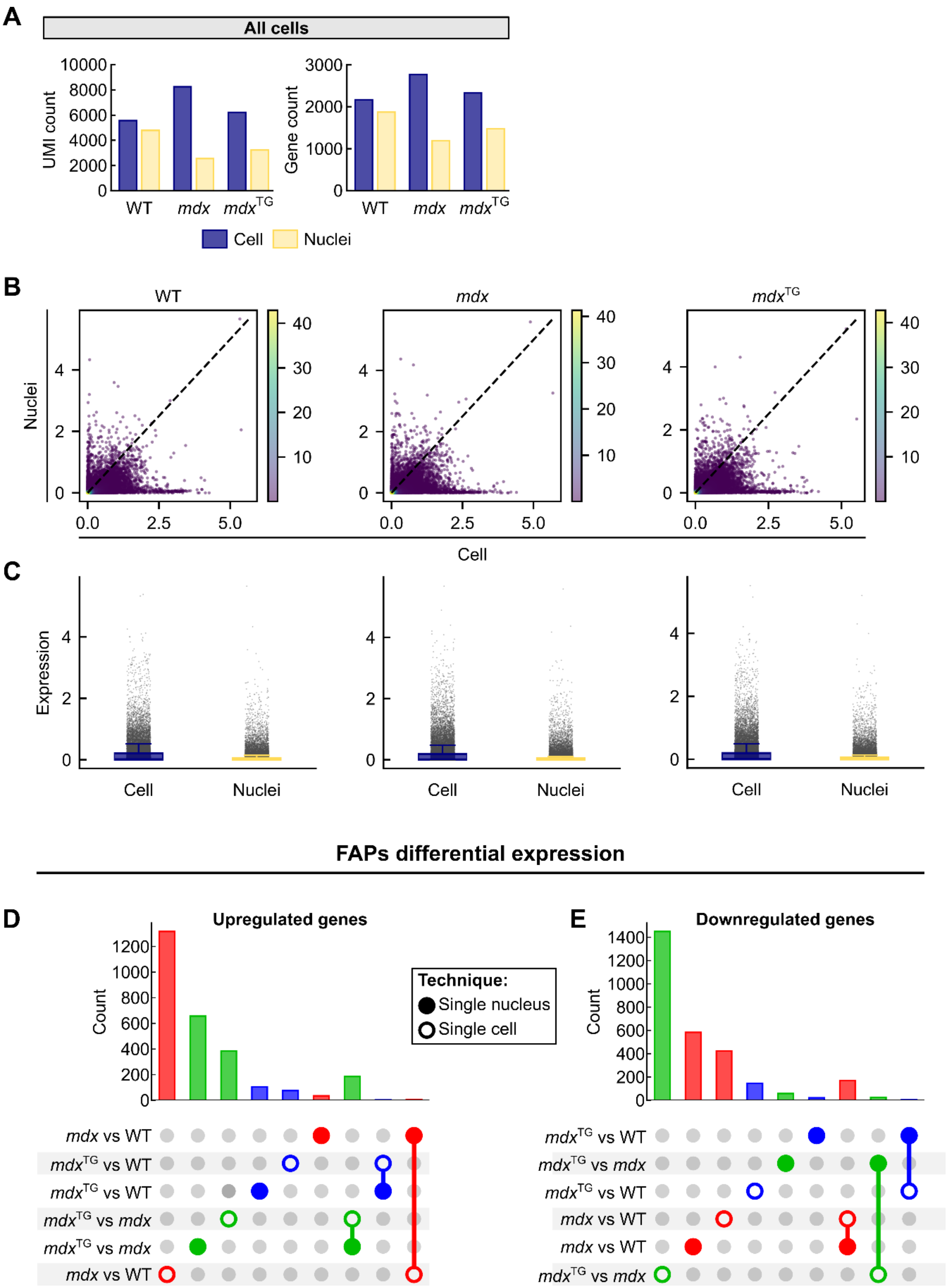


**Supplemental Figure 2. Comparison of scRNA-seq and snRNA-seq datasets.** (A) Comparison of median UMI counts and gene counts across all cells in wild-type, *mdx*, *mdx*$^{TG}$ in scRNA-seq and snRNA-seq. (**B**) Correlation of normalized gene expression (normalized mean expression of each gene across all cells) between scRNA-seq and snRNA-seq separated by genotype. Color map indicates density. (**C**) Mean expression of all genes represented as a boxplot. All cells were used in scRNA-seq and snRNA-seq. Expression profiles are separated by genotype. (**D**) Comparison of upregulated and (**E**) downregulated genes identified in FAPs between scRNA-seq and snRNA-seq.

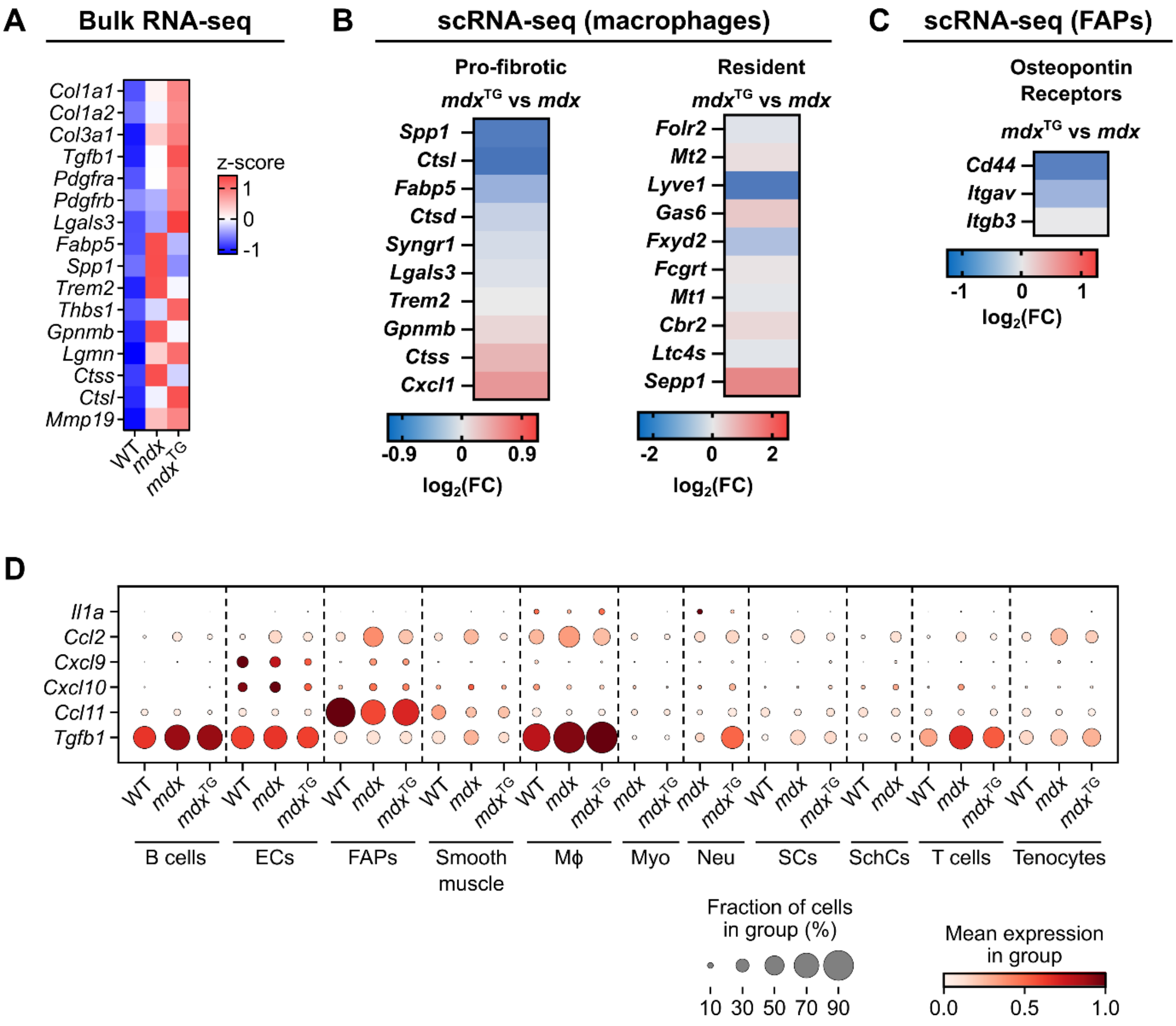


**Supplemental Figure 3. SSPN reduces expression of pro-fibrotic macrophage genes and osteopontin receptors.** (**A**) Heatmap of collagens, platelet-derived growth factor genes, pro-fibrotic macrophage genes, and ECM remodeling genes in bulk RNA-seq. (**B**) Heatmap of pro-fibrotic macrophage and resident macrophage genes in macrophages from scRNA-seq. (**C**) Osteopontin receptors in FAPs from scRNA-seq. (**D**) Expression of cytokine genes in all cell types identified by single cell RNA-seq.

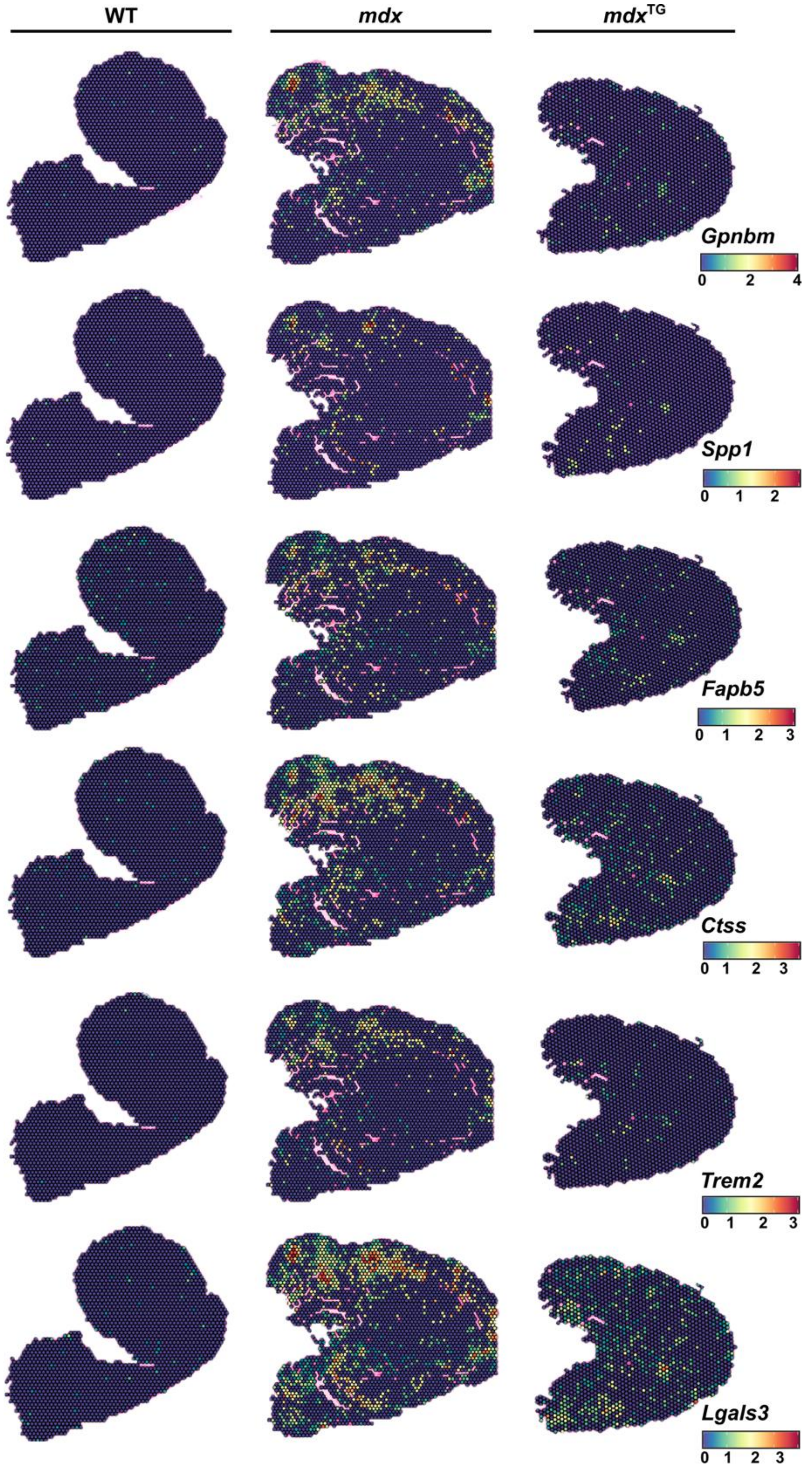


**Supplemental Figure 4. Pro-fibrotic macrophage genes localize to inflammatory and fibrotic regions.** Spatial expression of pro-fibrotic macrophage genes in 15-week-old wild-type, *mdx*, and *mdx*$^{TG}$ quadriceps muscle using spatial RNA-seq.

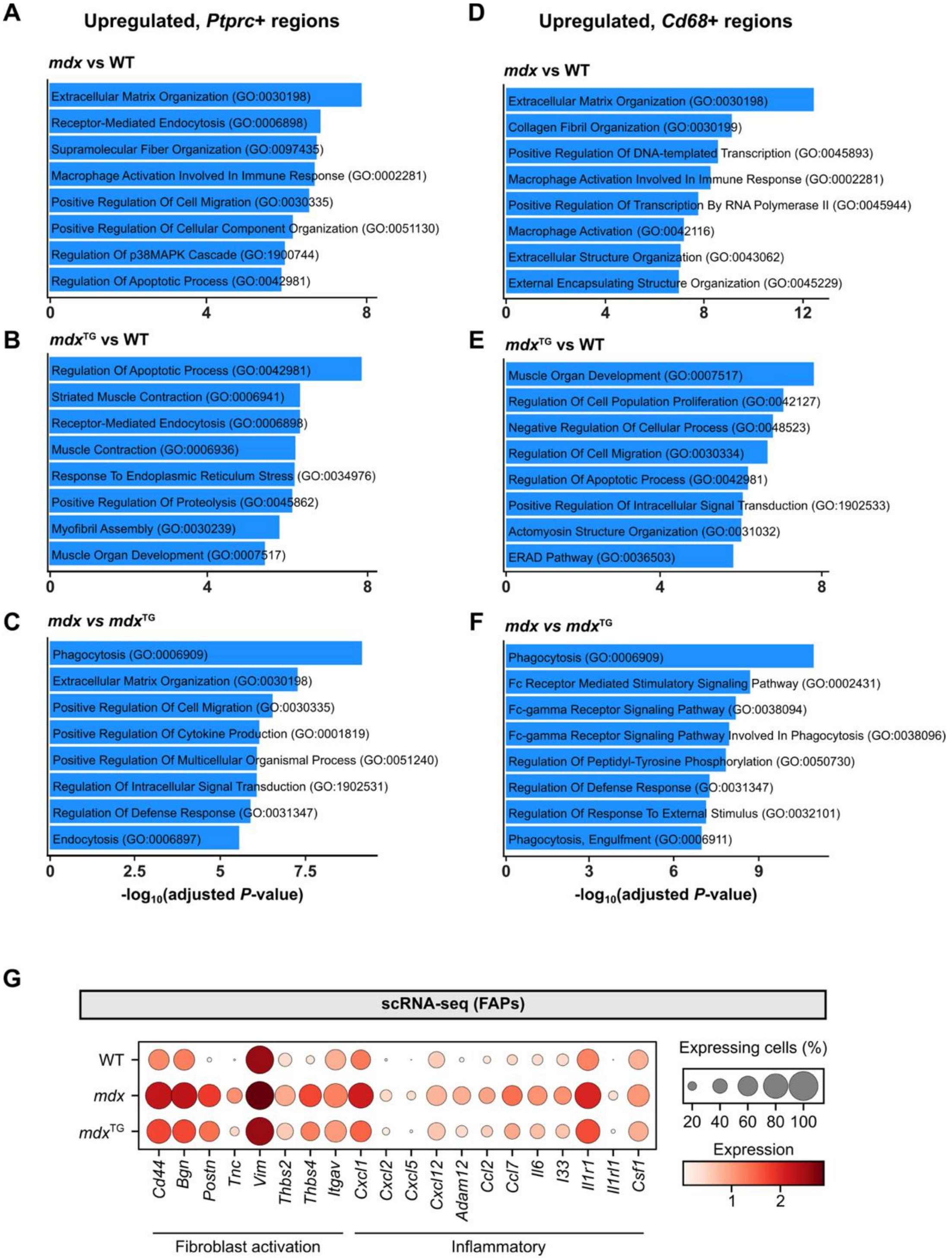


**Supplemental Figure 5. Inflammatory regions are associated with ECM production in *mdx* muscle.** GO term enrichment in regions highly expressing *Ptprc* in spatial RNA-seq comparing (**A**) *mdx* versus wild-type (**B**) *mdx*$^{TG}$ versus wild-type and (**C**) *mdx* versus *mdx*$^{TG}$. GO term enrichment in regions highly expressing *Cd68* in spatial RNA-seq comparing (**D**) *mdx* versus wild-type, (**E**) *mdx*$^{TG}$ versus wild-type and (**F**) *mdx* versus *mdx*$^{TG}$. (**G**) Dotplot from scRNA-seq showing expression of fibroblast activation markers and inflammatory genes in FAPs.

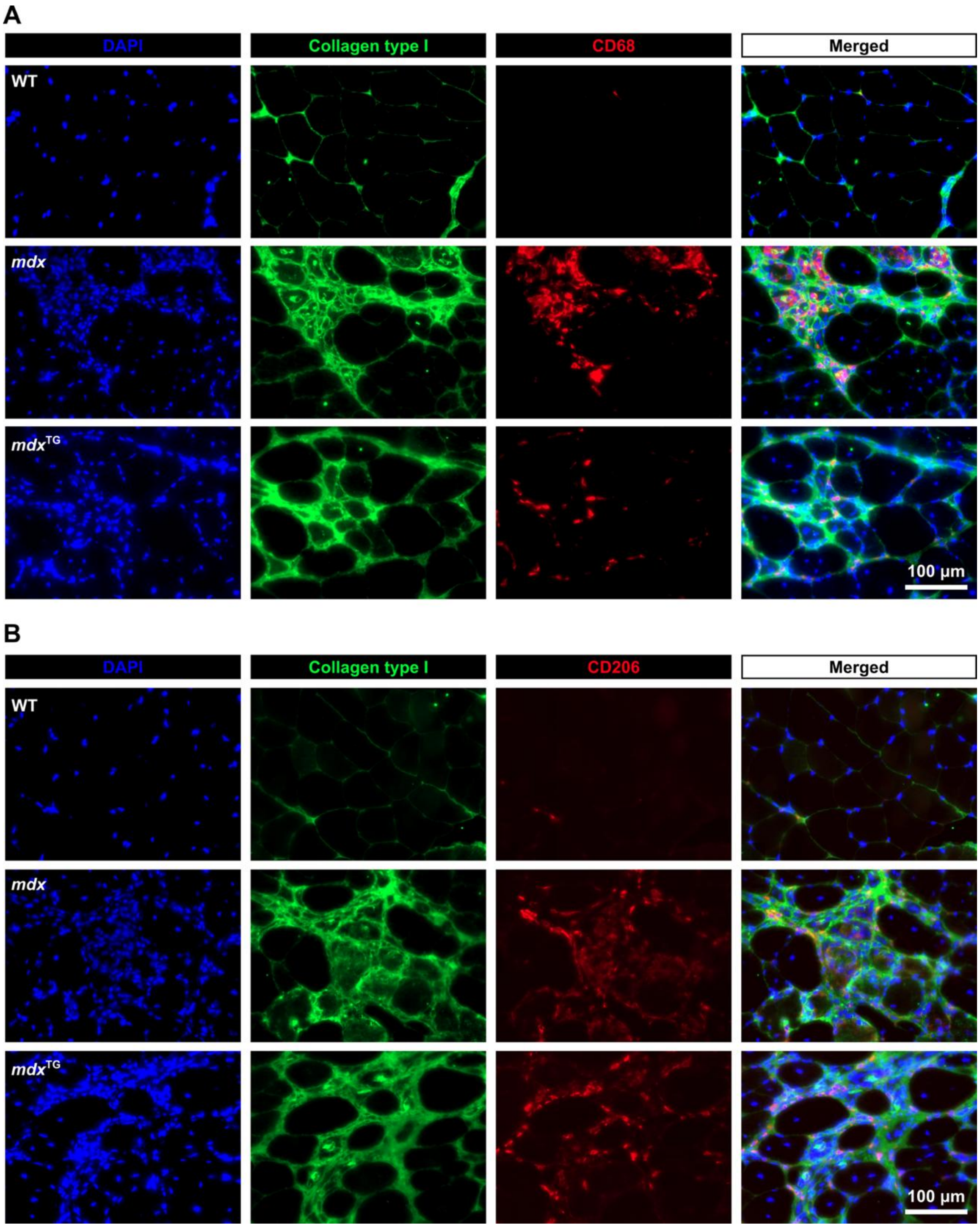


**Supplemental Figure 6. Macrophages localize to highly collagenous regions in *mdx* and *mdx*$^{TG}$ muscle.** (**A**) Immunofluorescence images of CD68$^{+}$ macrophages in wild-type, *mdx* and *mdx*$^{TG}$ quadriceps muscle. (**B**) immunofluorescence images of CD206$^{+}$ macrophages in wild-type, *mdx* and *mdx*$^{TG}$ quadriceps muscle.

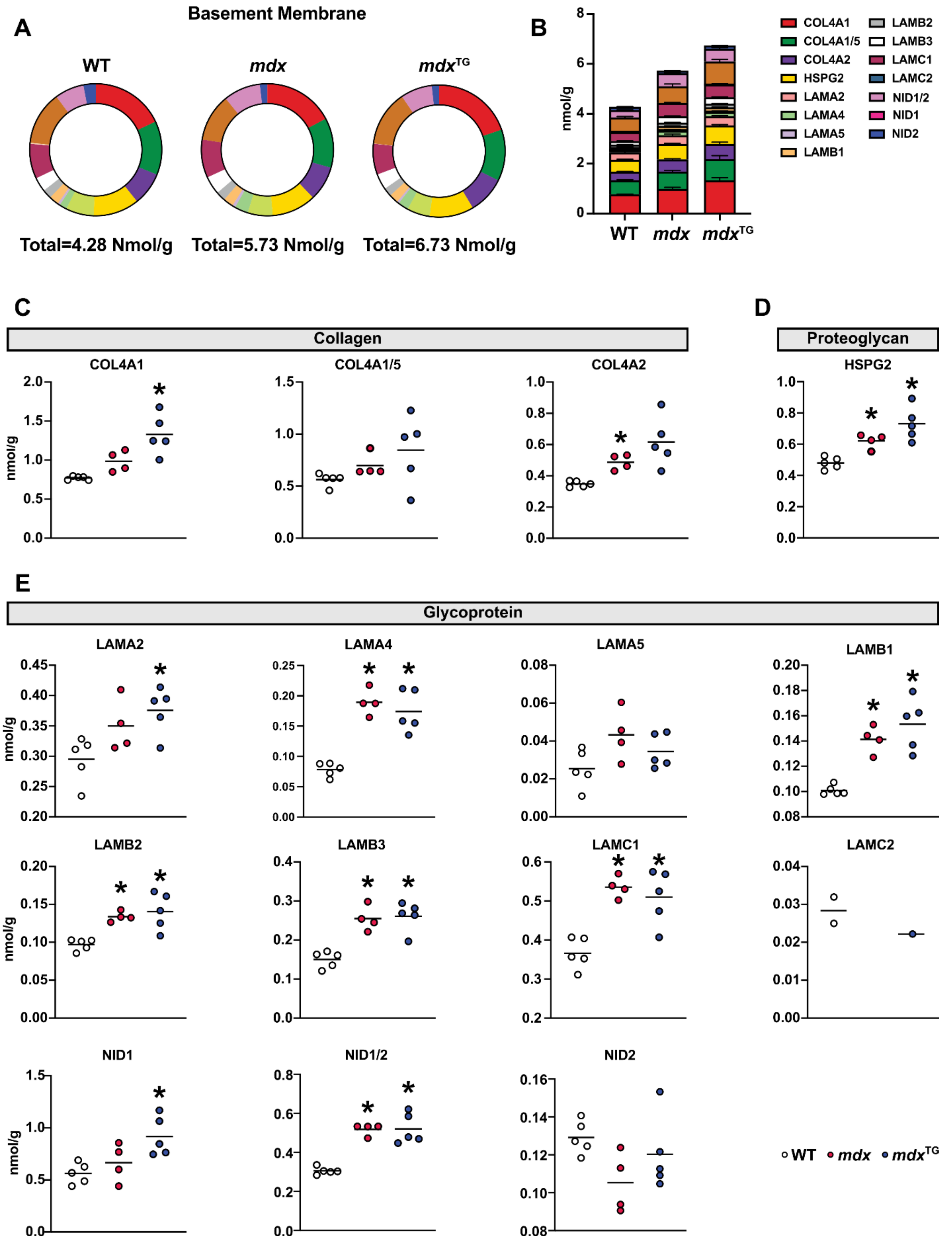
A
Basement Membrane
WT
mdx
mdxTG
Total=4.28 Nmol/g
Total=5.73 Nmol/g
Total=6.73 Nmol/g
B
nmol/g
WT
mdx
mdxTG
COL4A1
COL4A1/5
COL4A2
HSPG2
LAMA2
LAMA4
LAMA5
LAMB1
LAMB2
LAMB3
LAMC1
LAMC2
NID1/2
NID1
NID2
C
Collagen
COL4A1
COL4A1/5
COL4A2
D
Proteoglycan
HSPG2
E
Glycoprotein
LAMA2
LAMA4
LAMA5
LAMB1
LAMB2
LAMB3
LAMC1
LAMC2
NID1
NID1/2
NID2
WT
mdx
mdxTG

**Supplemental Figure 7. Overview of basement membrane proteins using quantitative proteomics.** (**A**) Proportion of basement membrane proteins and (**B**) absolute quantification. Individual protein separated by (**C**) collagens, (**D**) proteoglycans and (**E**) glycoproteins. *$P$<0.05 versus wild-type. Statistical analysis by Welch *t*-test with Benjamini-Hochberg correction.

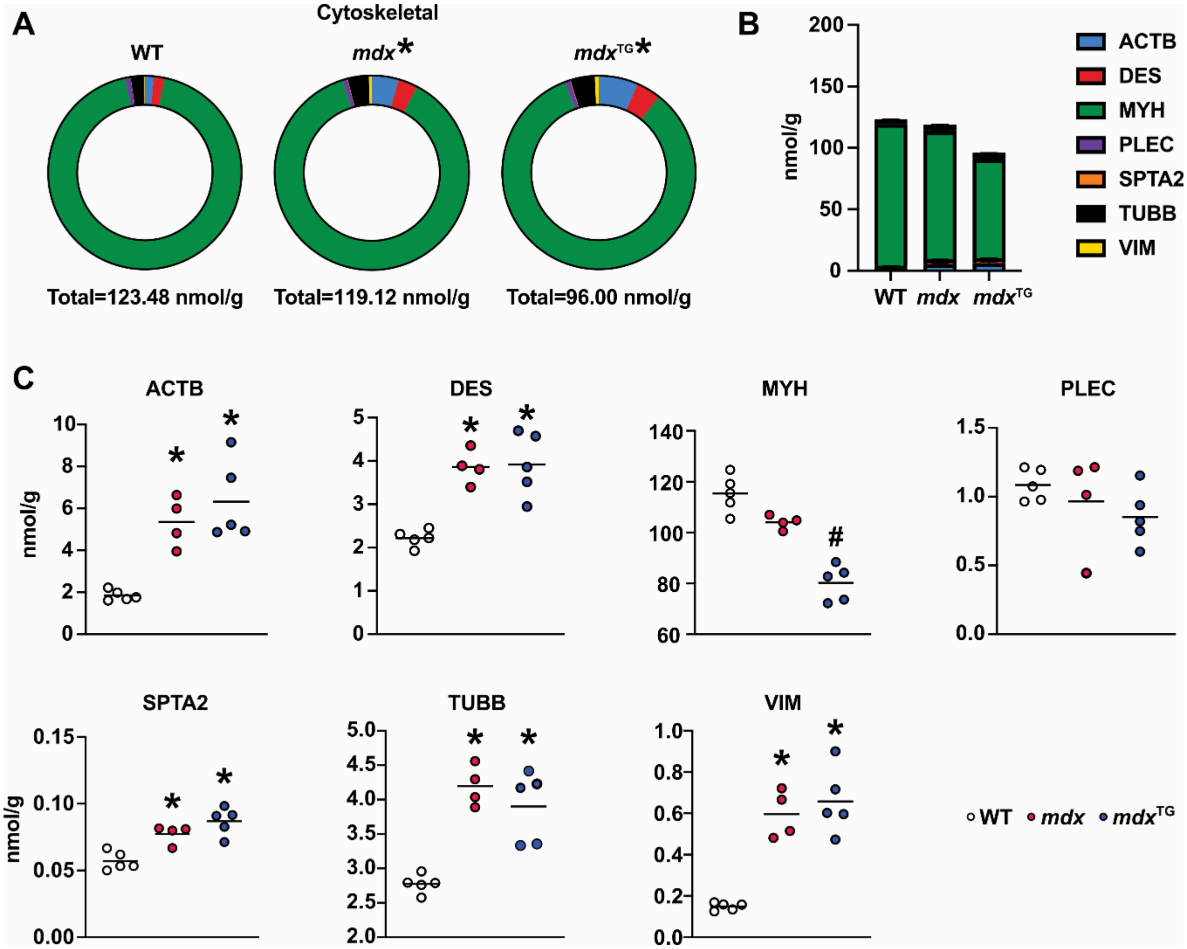


**Supplemental Figure 8. Overview of cytoskeletal proteins using quantitative proteomics.** (**A**) Proportion of cytoskeletal proteins and (**B**) absolute quantification. (**C**) Plots of individual protein abundances. *$P$<0.05 versus wild-type, #$P$<0.05 versus wild-type and *mdx*. Statistical analysis in A by centered log ratio transformation followed by pairwise PERMANOVA and Holm correction; in C by Welch $t$-test with Benjamini-Hochberg correction.

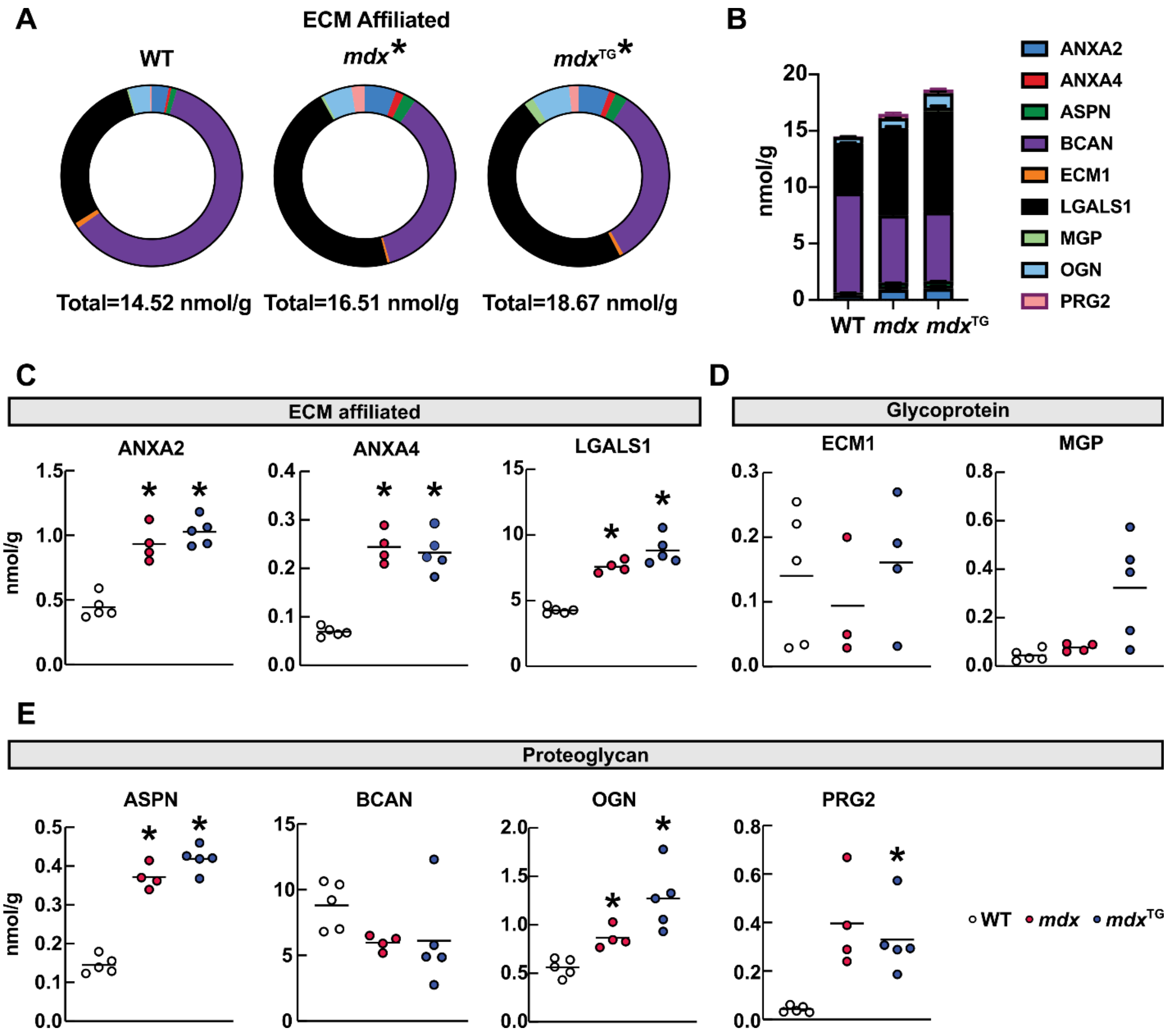


**Supplemental Figure 9. Overview of ECM affiliated proteins using quantitative proteomics.** (**A**) Proportion of ECM affiliated proteins and (**B**) absolute quantification. Individual protein separated by (**C**) ECM affiliated, (**D**) glycoproteins and (**E**) proteoglycans. *$P$<0.05 versus wild-type. Statistical analysis in A by centered log ratio transformation followed by pairwise PERMANOVA and Holm correction; in C-E by Welch $t$-test with Benjamini-Hochberg correction.

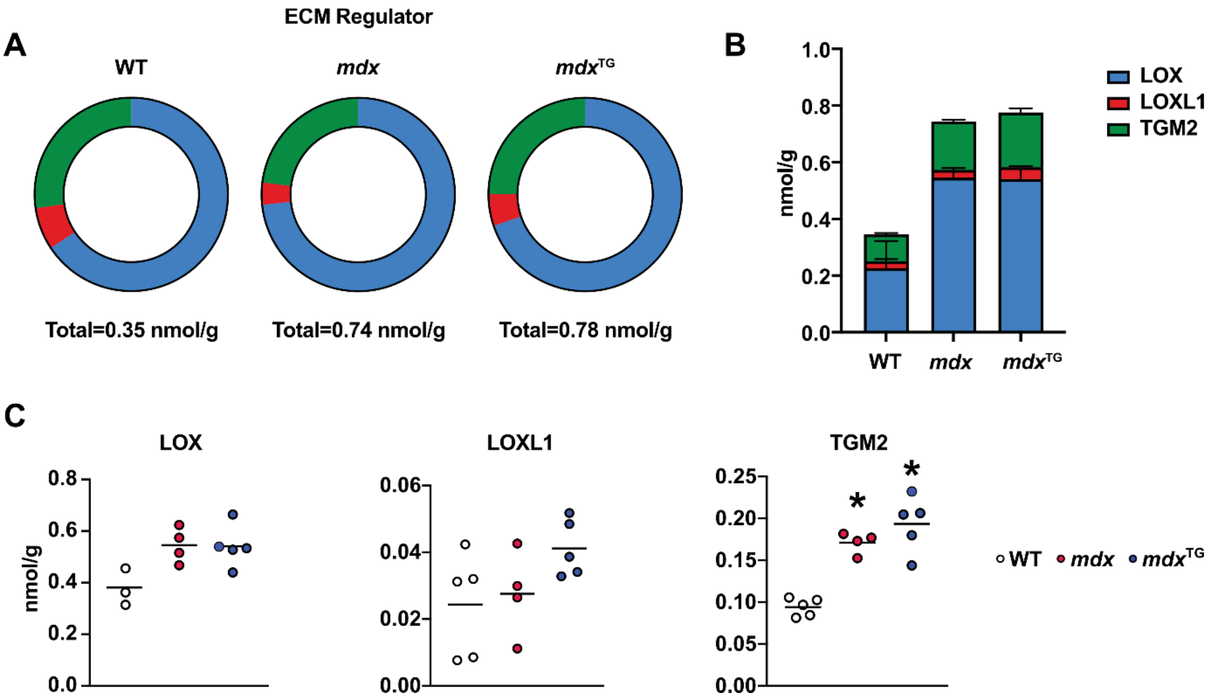


**Supplemental Figure 10. Overview of ECM regulator proteins using quantitative proteomics.** (**A**) Proportion of ECM regulator proteins and (**B**) absolute quantification. (**C**) Plots of individual protein abundances. *$P$<0.05 versus wild-type. Statistical analysis by Welch $t$-test with Benjamini-Hochberg correction.

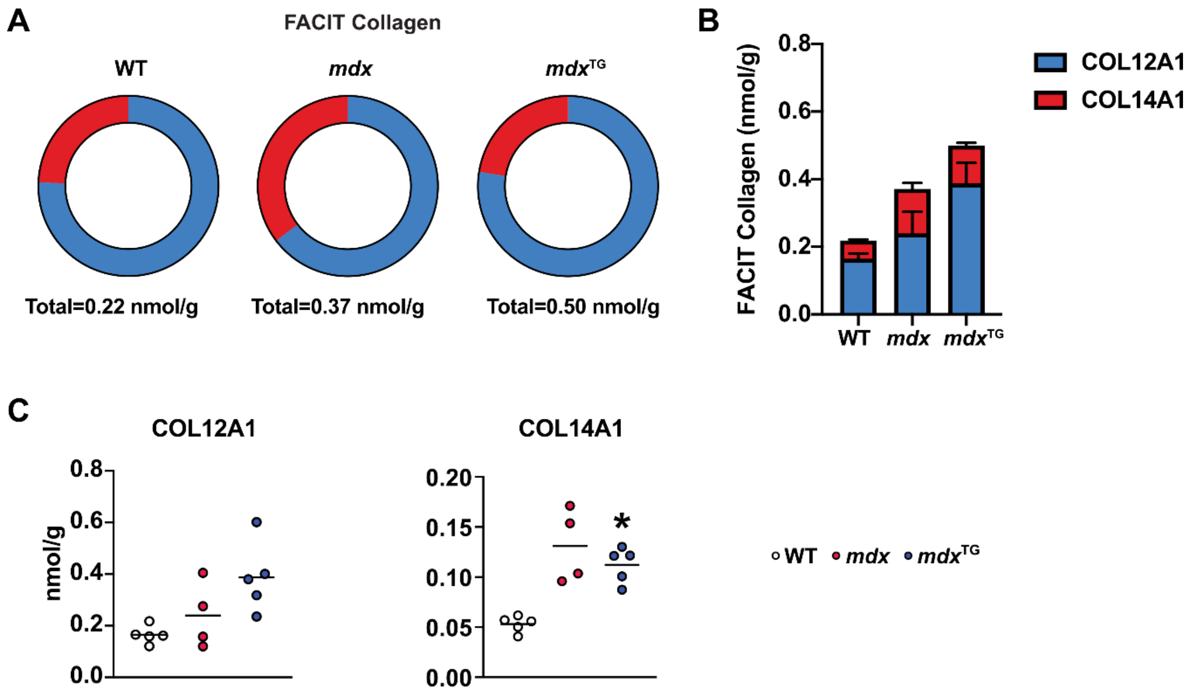


**Supplemental Figure 11. Overview of FACIT collagens using quantitative proteomics.** (**A**) Proportion of FACIT collagens and (**B**) absolute quantification. (**C**) Plots of individual protein abundances. **P*<0.05 versus wild-type. Statistical analysis by Welch *t*-test with Benjamini-Hochberg correction.

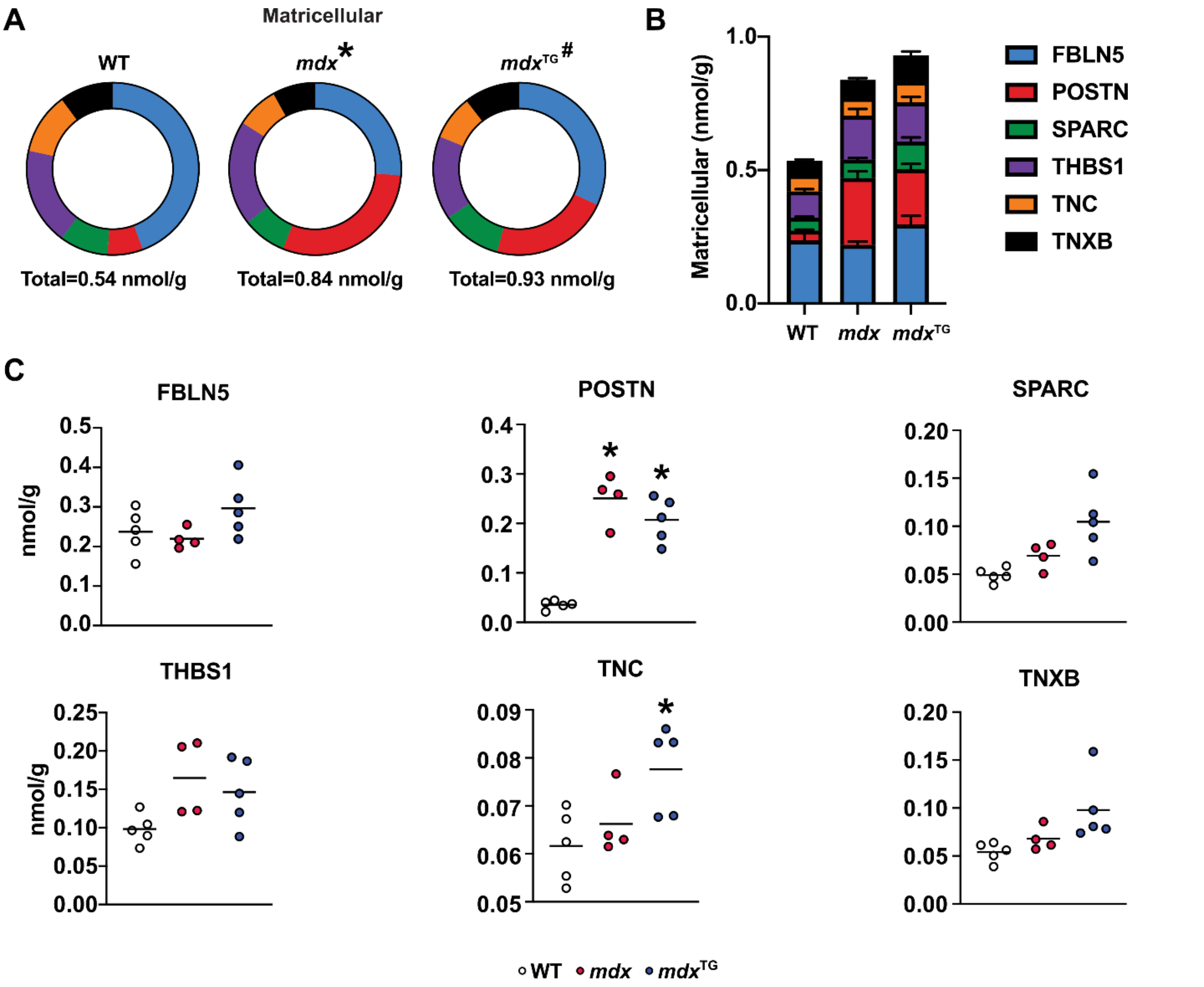


**Supplemental Figure 12. Overview of matricellular proteins using quantitative proteomics.** (**A**) Proportion of matricellular proteins and (**B**) absolute quantification. (**C**) Plots of individual protein abundances. **P*<0.05 versus wild-type, #*P*<0.05 versus wild-type and *mdx*. Statistical analysis in A by centered log ratio transformation followed by pairwise PERMANOVA and Holm correction; in C by Welch *t*-test with Benjamini-Hochberg correction.

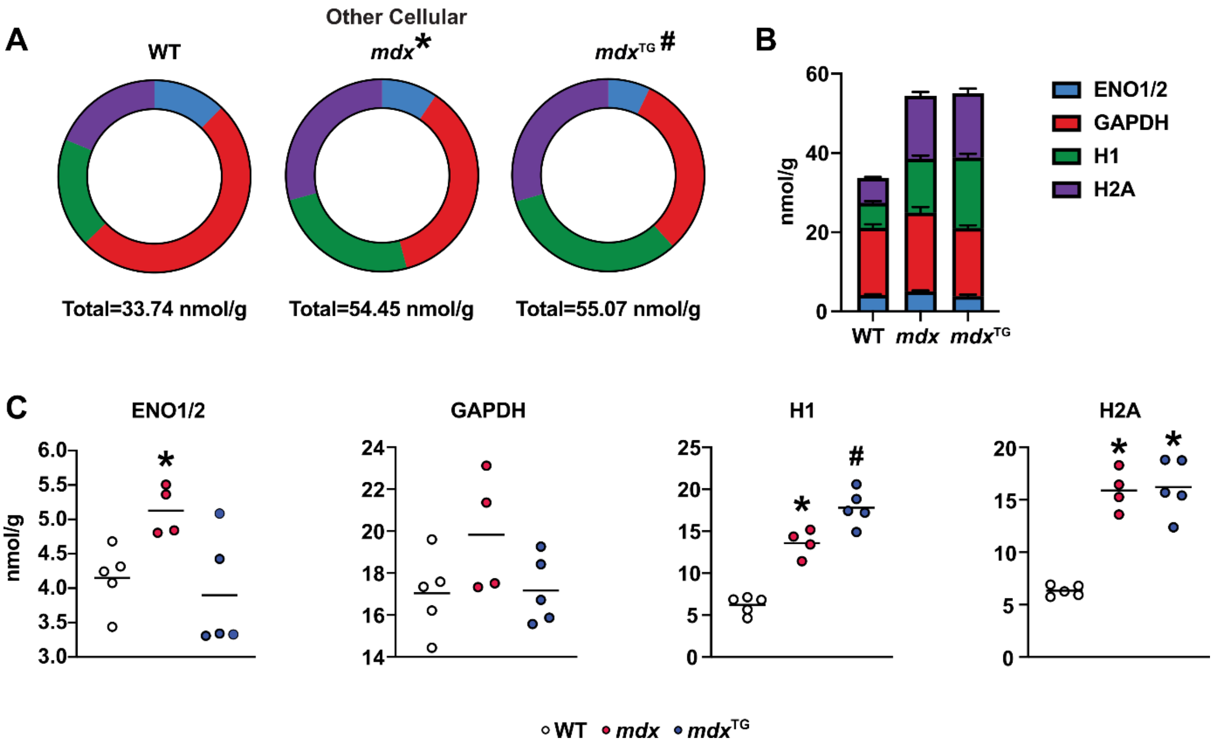


**Supplemental Figure 13. Overview of other cellular proteins using quantitative proteomics.** (**A**) Proportion of other cellular proteins and (**B**) their absolute quantification. (**C**) Plots of individual protein abundances. *$P$<0.05 versus wild-type, $^{\#}P$<0.05 versus wild-type and *mdx*. Statistical analysis in A by centered log ratio transformation followed by pairwise PERMANOVA and Holm correction; in C by Welch *t*-test with Benjamini-Hochberg correction.

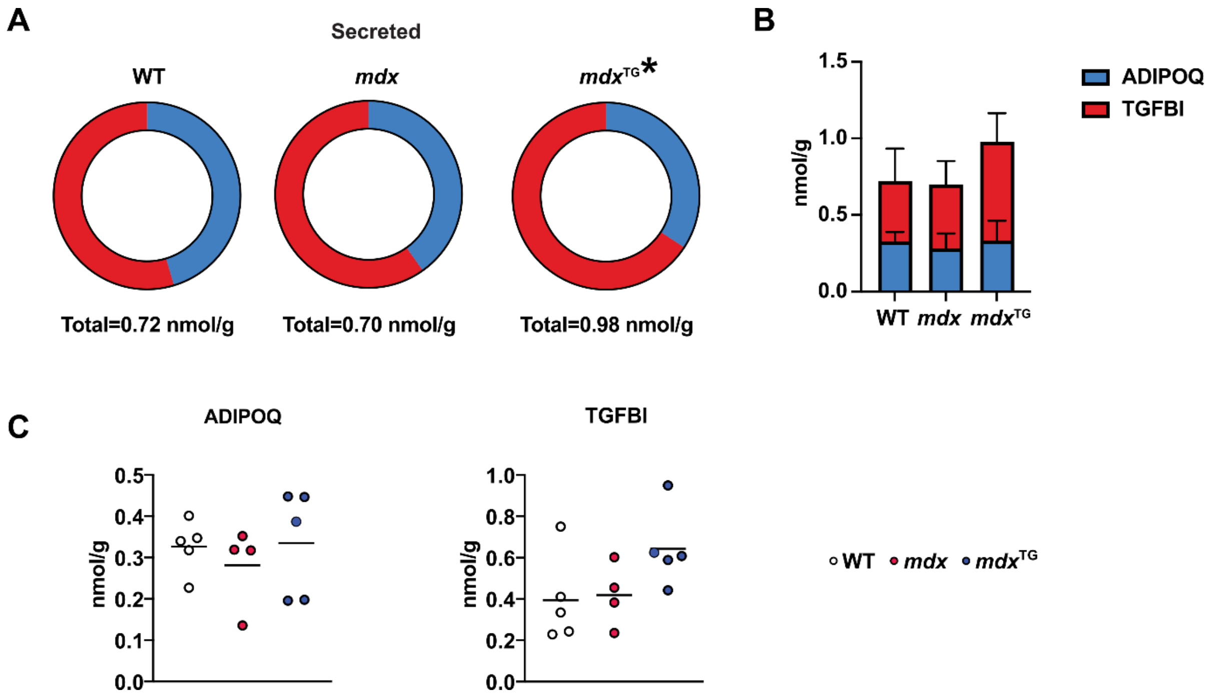


**Supplemental Figure 14. Overview of secreted proteins using quantitative proteomics.** (**A**) Proportion of secreted proteins and (**B**) absolute quantification. (**C**) Plots of individual protein abundances. *$P$<0.05 versus wild-type. Statistical analysis by centered log ratio transformation followed by pairwise PERMANOVA and Holm correction.

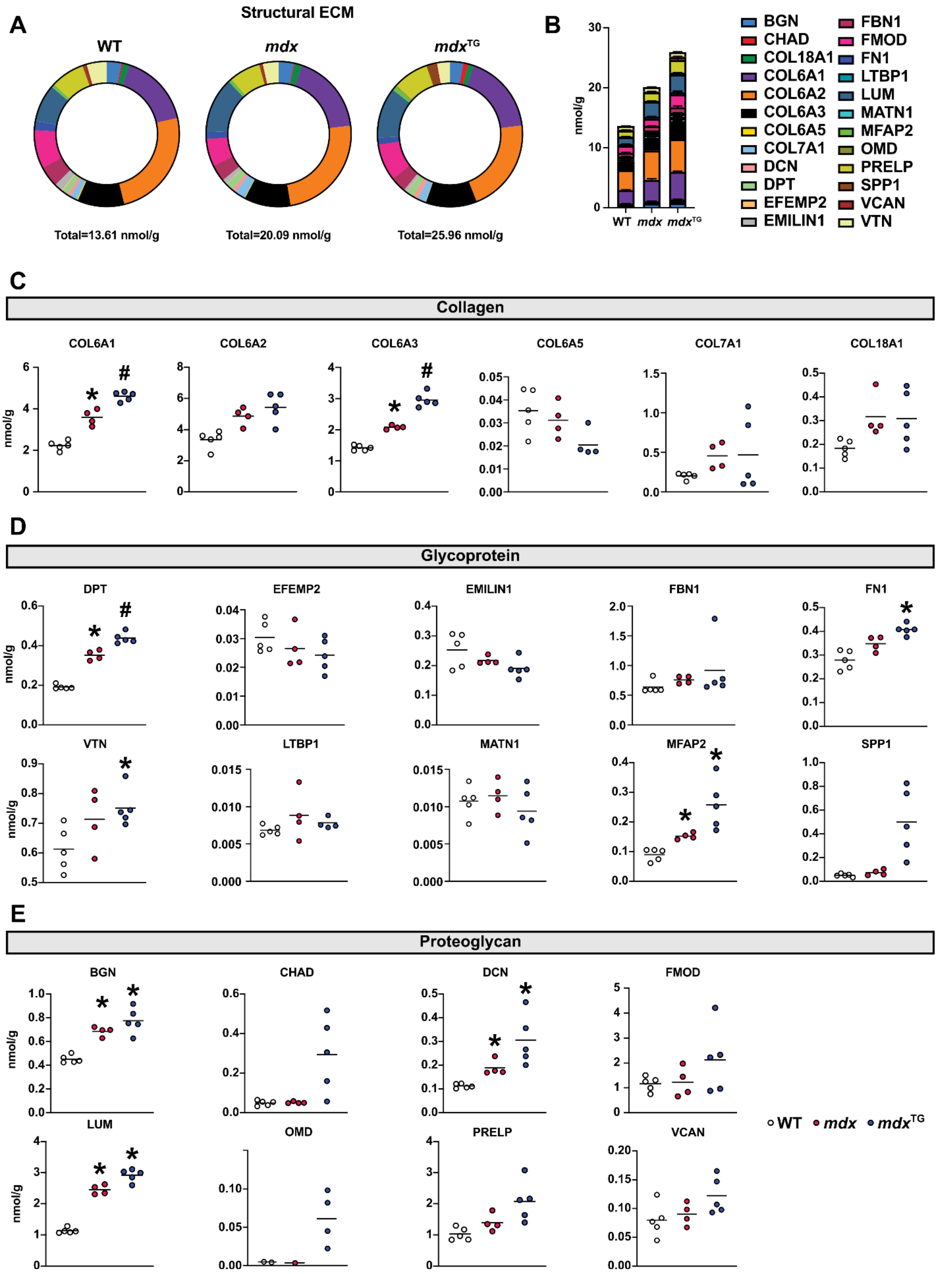

A
Structural ECM
WT
mdx
mdxTG
Total=13.61 nmol/g
Total=20.09 nmol/g
Total=25.96 nmol/g
B
nmol/g
WT mdx mdxTG
BGN
CHAD
COL18A1
COL6A1
COL6A2
COL6A3
COL6A5
COL7A1
DCN
DPT
EFEMP2
EMILIN1
FBN1
FMOD
FN1
LTBP1
LUM
MATN1
MFAP2
OMD
PRELP
SPP1
VCAN
VTN
C
Collagen
COL6A1
COL6A2
COL6A3
COL6A5
COL7A1
COL18A1
D
Glycoprotein
DPT
EFEMP2
EMILIN1
FBN1
FN1
VTN
LTBP1
MATN1
MFAP2
SPP1
E
Proteoglycan
BGN
CHAD
DCN
FMOD
LUM
OMD
PRELP
VCAN
WT mdx mdxTG

**Supplemental Figure 15. Overview of structural ECM proteins using quantitative proteomics.** (**A**) Proportion of structural ECM proteins and (**B**) absolute quantification. Individual protein separated by (**C**) collagens, (**D**) glycoproteins and (**E**) proteoglycans. *$P$<0.05 versus wild-type, $^{\#}P$<0.05 versus wild-type and *mdx*. Statistical analysis by Welch *t*-test with Benjamini-Hochberg correction.

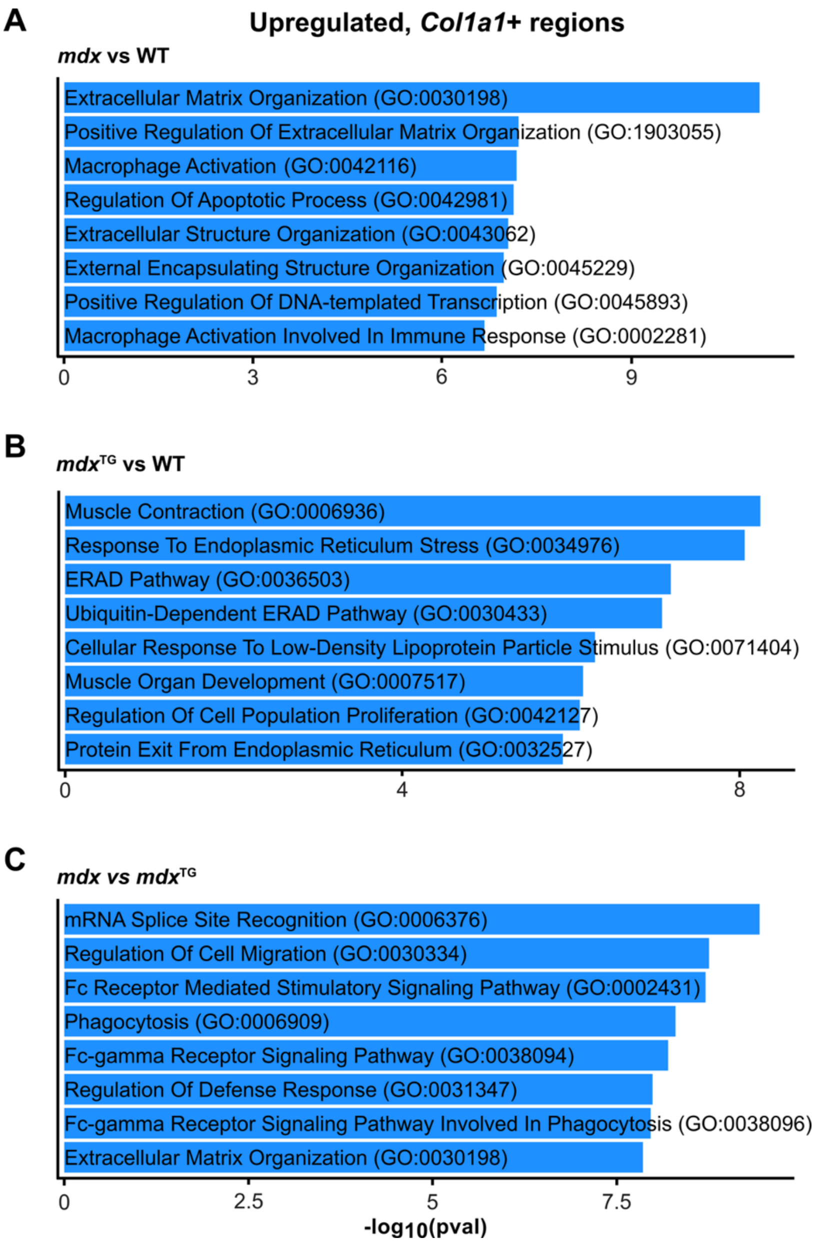


**Supplemental Figure 16. Collagen-expressing regions are associated with inflammation in *mdx* muscle.** GO term enrichment in regions highly expressing *Col1a1* in spatial RNA-seq comparing (**A**) *mdx* versus wild-type (**B**) *mdx*$^{TG}$ versus wild-type and (**C**) *mdx* versus *mdx*$^{TG}$.

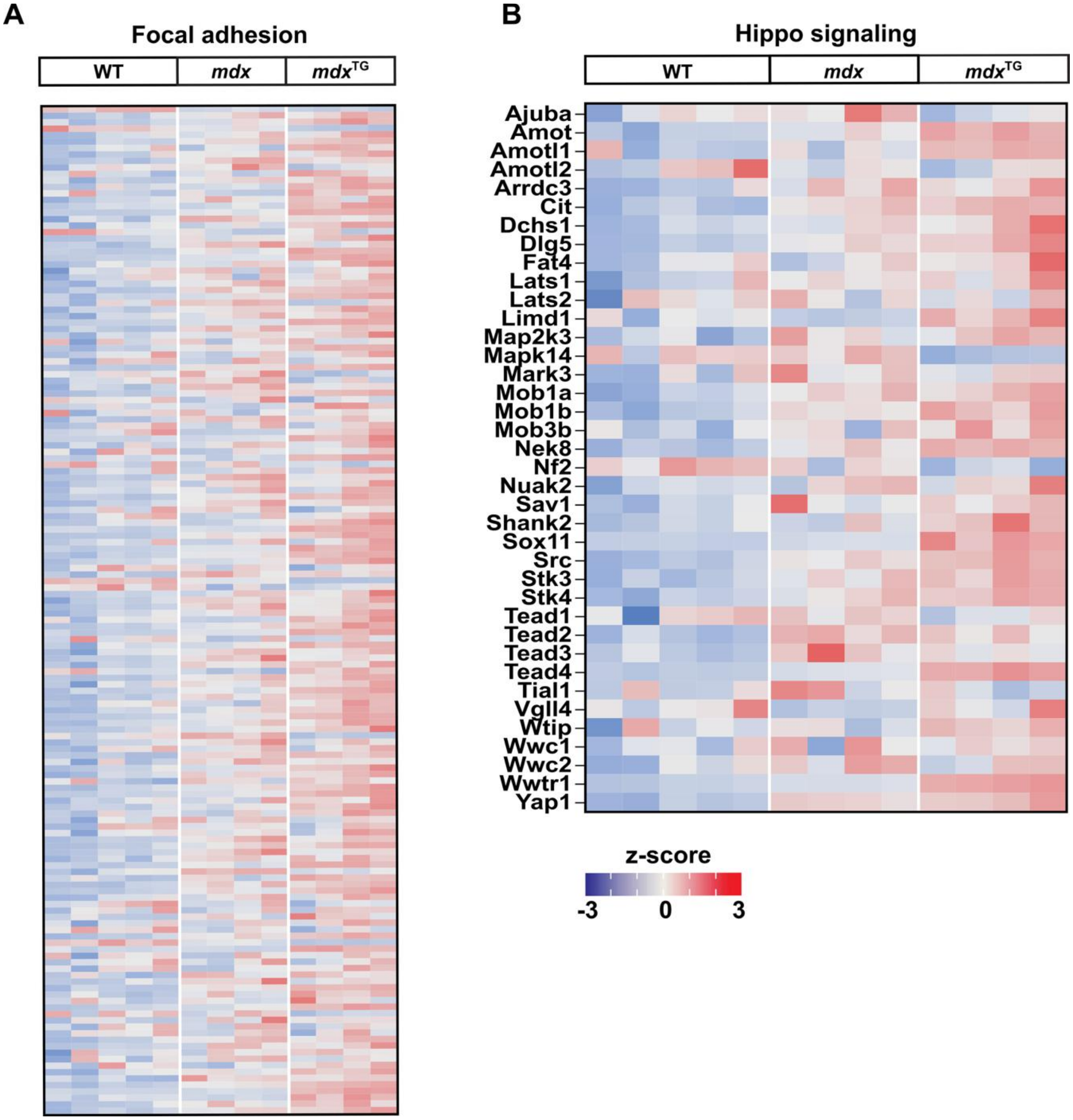


**Supplemental Figure 17. Focal adhesion and Hippo signaling genes are upregulated in *mdx* mice.** Heatmaps showing expression of (**A**) focal adhesion genes and (**B**) Hippo signaling genes from bulk RNA sequencing analysis of tibialis anterior muscle.

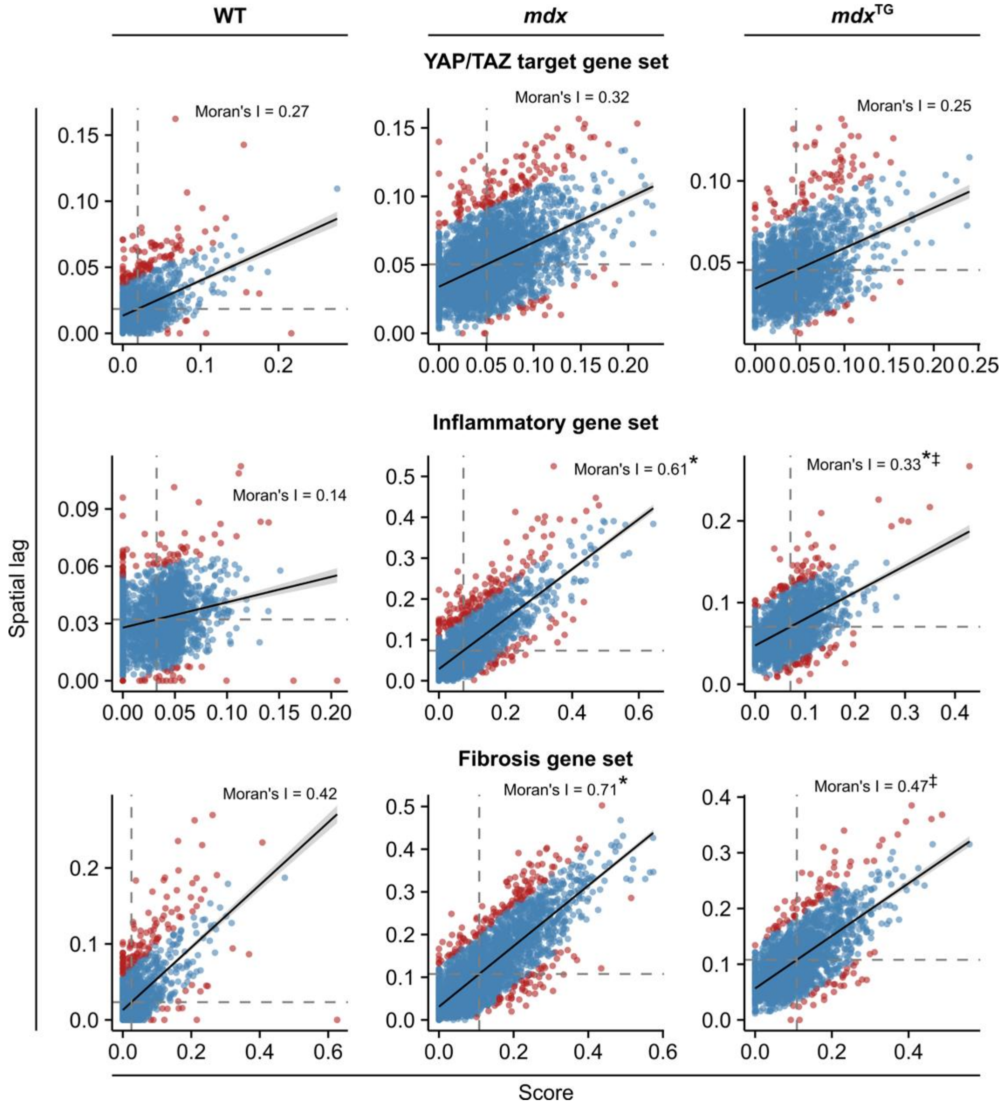


**Supplemental Figure 18. Inflammatory and fibrotic gene sets are highly spatially localized in *mdx* muscle.** Plots of raw score (x-axis) vs spatial lag of raw scores (y-axis; average score of all surrounding spots) for YAP/TAZ targets, inflammatory, and fibrosis gene sets using spatial RNA-seq data from wild-type, *mdx*, and *mdx*$^{TG}$ quadriceps muscle. The degree of spatial autocorrelation was determined by Moran's I, which is approximately equal to the slope of each fit curve. Statistically significant differences in Moran's I are indicated as *versus wild-type and ‡versus *mdx* determined by a z-test described in detail in the methods.

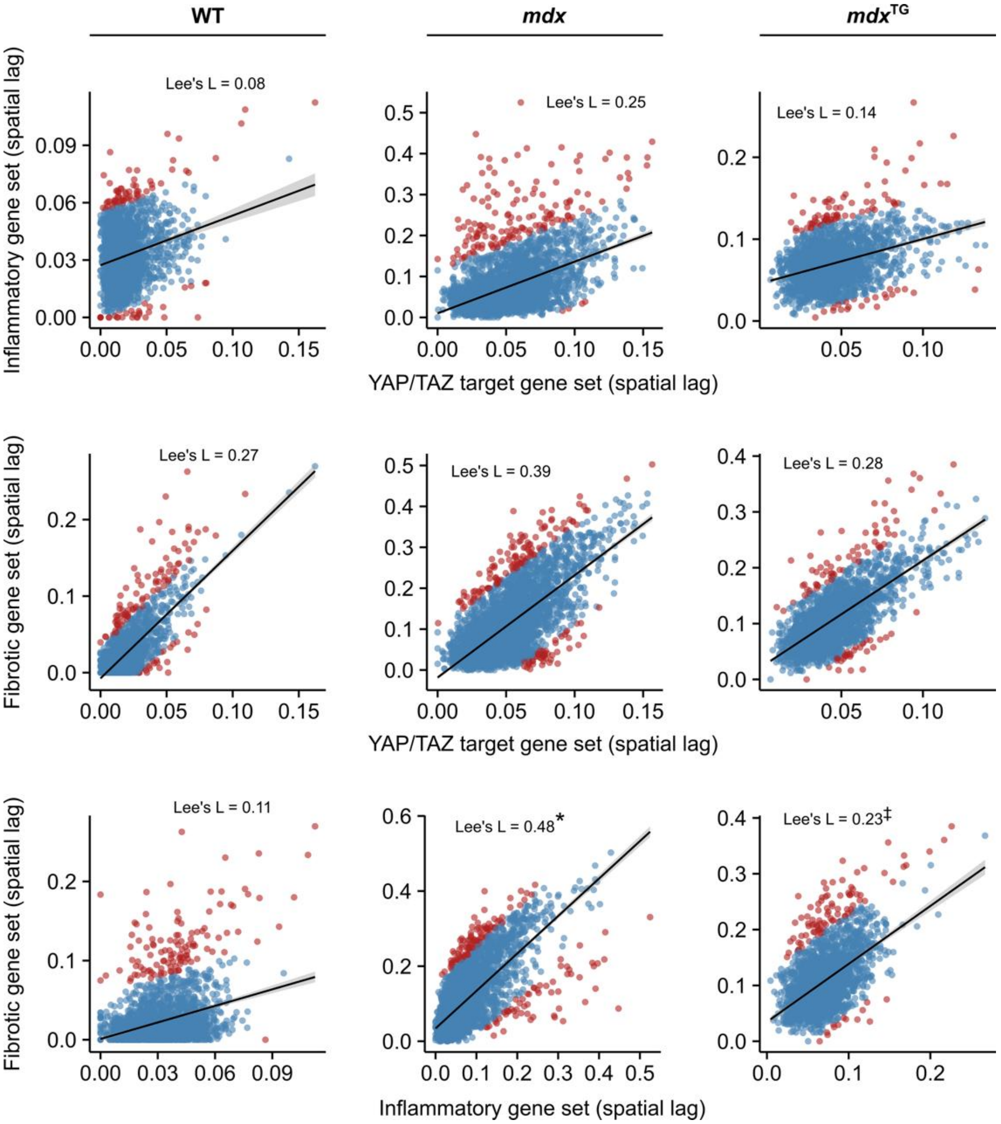


**Supplemental Figure 19. Fibrotic and inflammatory gene sets are highly spatially co-localized in *mdx* muscle.** Plots of spatial lag for each gene set score pair in wild-type, *mdx*, and *mdx*$^{TG}$ quadriceps muscle from spatial RNA-seq. Bivariate spatial correlation was determined by calculating Lee's L. Note that Lee's L does not correspond to the slope of the fit line, and plots are solely visual aids. Statistically significant differences in Lee's L are indicated as *versus wild-type and ‡versus *mdx* determined by a z-test described in detail in the methods.

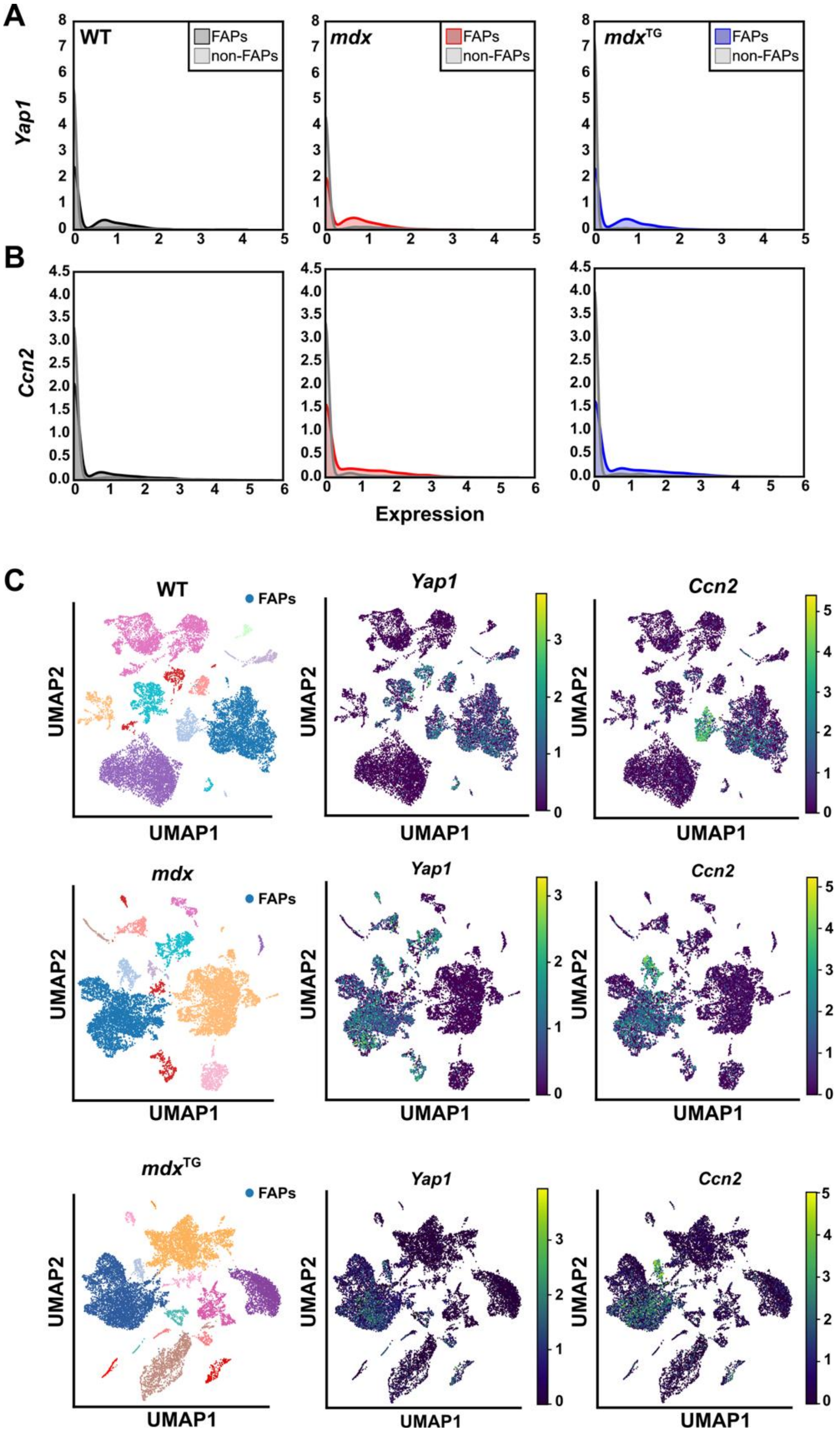


**Supplemental Figure 20. *Yap1* is expressed in FAPs.** (**A**) Kernel density plots of *Yap1* and (**B**) *Ccn2* in FAPs compared to all other cells using scRNA-seq. (**C**) UMAP visualization of *Yap1* and *Ccn2* expression in scRNA-seq datasets from wild-type, *mdx*, and *mdx*$^{TG}$.

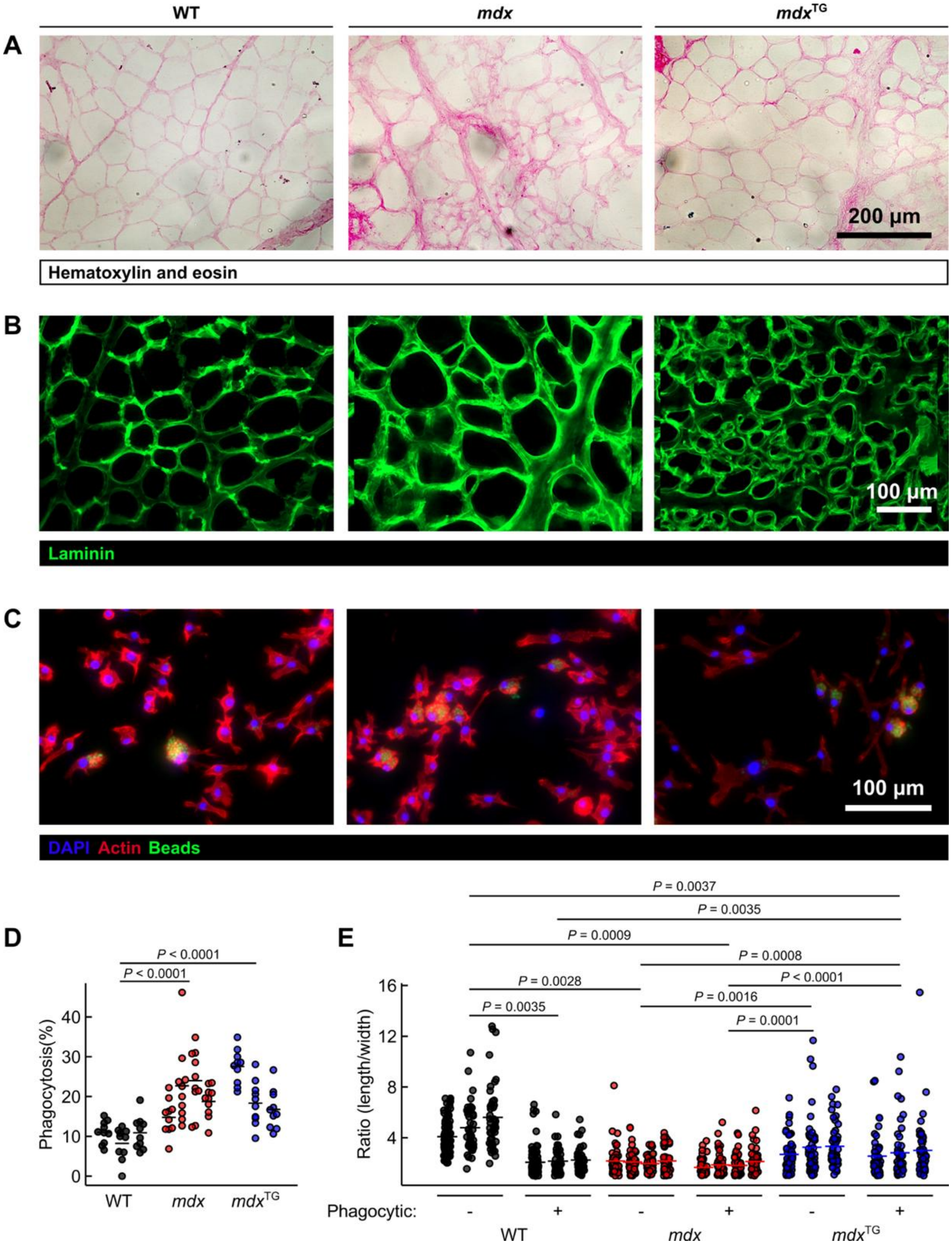


**Supplemental Figure 21. Myoscaffolds influence phagocytosis capacity and macrophage morphology.** (**A**) H&E staining after muscle cryosection decellularization. (**B**) Laminin staining after muscle cryosection decellularization. (**C**) Bone-marrow derived macrophages cultured on myoscaffolds generated from wild-type, *mdx*, and *mdx*$^{TG}$ quadriceps muscle were incubated with Zymosan A bioparticles conjugated with Alexa Fluor 488 and imaged. (**D**) Quantification of the percentage of macrophages with positive staining for Zymosan A bioparticles (n = 3-4 per genotype). Multiple images were captured per myoscaffold and the percentage of macrophages positive for internalized Zymosan A per image are represented as individual data points. Mean values of all images per myoscaffold are represented as horizontal lines. Statistical analysis by fitting a regression model followed by wild-cluster bootstrap. (**E**) Length-to-width ratio of non-phagocytic and phagocytic macrophages cultured on wild-type, *mdx*, and *mdx*$^{TG}$ myoscaffolds. Phalloidin was used to visualize actin. Individual datapoints represent a single macrophage. Data are separated by myoscaffold (n = 3-4 per genotype), with horizontal bars representing the average ratio for all macrophages within a single myoscaffold. Statistical analysis by generalized linear model with cluster-robust standard errors.

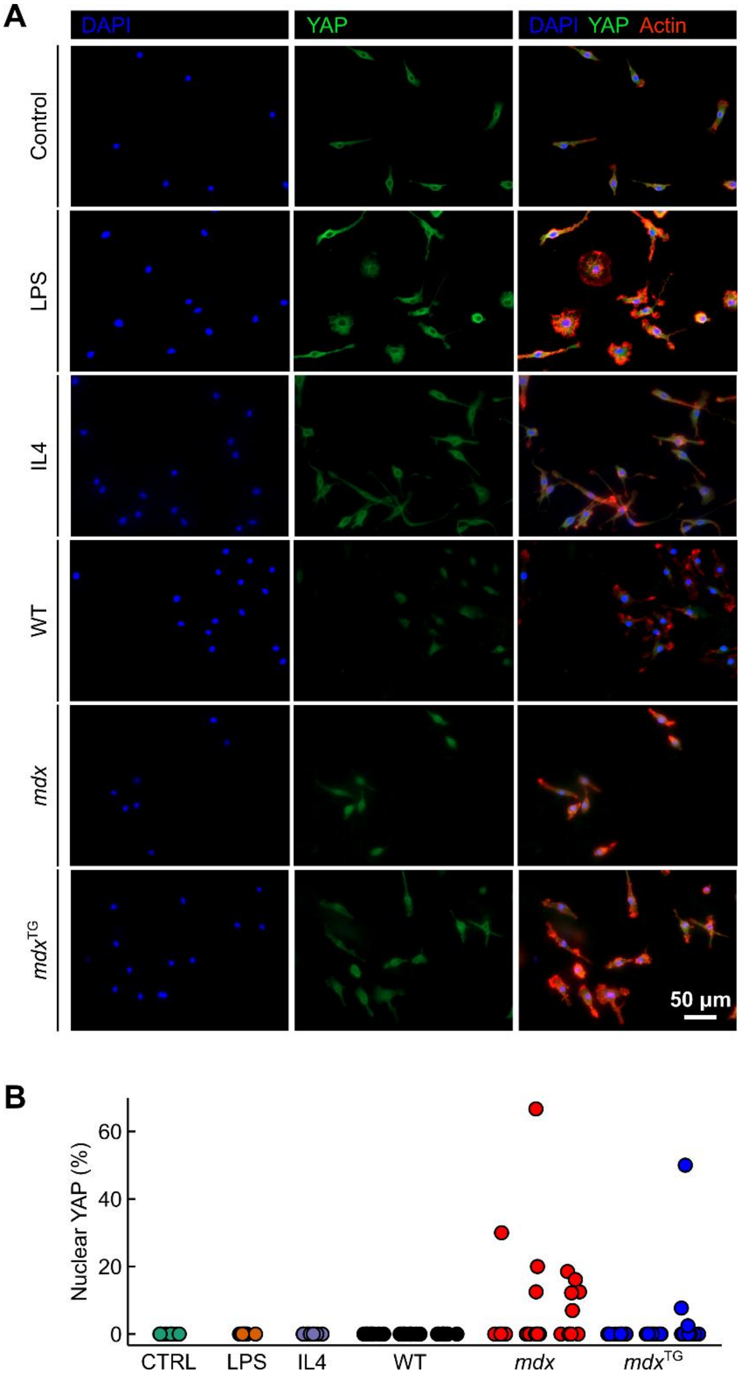


**Supplemental Figure 22. YAP1 localization in macrophages is not affected by myoscaffolds.** (**A**) Immunofluorescence images of YAP1 in bone marrow-derived macrophages cultured on the glass alone (control or with LPS or IL4; all n=1) or on slides coated with wild-type, *mdx*, or *mdx*$^{TG}$ myoscaffolds (n=3). (**B**) The percentage of macrophages with nuclear YAP1. Each datapoint represents the percentage of macrophages with nuclear YAP1 per image. Datapoints are grouped according to their corresponding myoscaffold within each genotype.
